# Bni5 tethers myosin-II to septins to enhance retrograde actin flow and the robustness of cytokinesis

**DOI:** 10.1101/2023.11.07.566094

**Authors:** Hiroki Okada, Xi Chen, Kangji Wang, Joseph Marquardt, Erfei Bi

## Abstract

The collaboration between septins and myosin-II in driving processes outside of cytokinesis remains largely uncharted. Here, we demonstrate that Bni5 in the budding yeast *S. cerevisiae* interacts with myosin-II, septin filaments, and the septin-associated kinase Elm1 via distinct domains at its N- and C-termini, thereby tethering the mobile myosin-II to the stable septin hourglass at the division site from bud emergence to the onset of cytokinesis. The septin and Elm1-binding domains, together with a central disordered region, of Bni5 control timely remodeling of the septin hourglass into a double ring, enabling the actomyosin ring constriction. The Bni5-tethered myosin-II enhances retrograde actin cable flow, which contributes to the asymmetric inheritance of mitochondria-associated protein aggregates during cell division, and also strengthens cytokinesis against various perturbations. Thus, we have established a biochemical pathway involving septin-Bni5-myosin-II interactions at the division site, which can inform mechanistic understanding of the role of myosin-II in other retrograde flow systems.

**Summary:** Okada et al. have determined the molecular mechanism underlying the Bni5 interactions with septins and myosin-II at the cell division site and uncovered its roles in promoting retrograde actin flow and the robustness of cytokinesis in budding yeast.

## Introduction

Septins and non-muscle myosin-IIs are known to act together to drive cytokinesis in fungal and animal cells. In the budding yeast *S. cerevisiae*, the septin hourglass scaffolds the actomyosin ring (AMR) assembly in anaphase (Bi et al., 1998; Lippincott and Li, 1998; Schneider et al., 2013). At the onset of cytokinesis, the septin hourglass is remodeled into a double ring that sandwiches the AMR, allowing it to access the plasma membrane (PM) to initiate constriction (Chen et al., 2020; Cid et al., 2001; Lippincott et al., 2001; Ong et al., 2014; Palani et al., 2012; Tamborrini et al., 2018). In mammalian cells, septins and myosin-IIs (IIA, IIB, and IIC) act together to drive both furrow ingression and abscission, two distinct stages of cytokinesis that are thought to involve distinct contributions from different septin complexes and myosin-II isoforms (Estey et al., 2010; Hickson and O’Farrell, 2008; Joo et al., 2007; Karasmanis et al., 2019; Oegema et al., 2000; Renshaw et al., 2014; Wang et al., 2019). Septins interact with myosin-II either directly (Joo et al., 2007) or indirectly via anillin, a scaffolding protein that also binds RhoA, actin, myosin-II, septin, and IQGAP3 at the cleavage furrow and the intercellular bridge during cytokinesis (Adachi et al., 2014; El Amine et al., 2013; Gai et al., 2011; Garno et al., 2021; Hickson and O’Farrell, 2008; Maddox et al., 2007; Piekny and Glotzer, 2008; Straight et al., 2005; Wang et al., 2023). Thus, the septin-myosin-II interaction and its function in cytokinesis has been extensively analyzed.

In contrast, very little is known about how septins and myosin-II interact and function to drive processes outside of cytokinesis. In the budding yeast *S. cerevisiae*, Myo1 is targeted to the division site via a two-step process, with Bni5 mediating its targeting from bud emergence to the onset of cytokinesis and Iqg1 from the onset of anaphase to the completion of cytokinesis (Fang et al., 2010). These two targeting mechanisms overlap from the onset of anaphase to the onset of cytokinesis during which Bni5 gradually dissociates from the bud neck whereas Iqg1 gradually increases at the bud neck (Fang et al., 2010; Lee et al., 2002; Okada et al., 2021b). Thus, Bni5 defines the sole linker between Myo1 and the septin hourglass before anaphase. How Bni5 interacts with these proteins and functions during the cell cycle have remained elusive. Myo1 has been shown to facilitate the retrograde actin cable flow (RACF) that is initiated by actin polymerization at the bud tip, and this role of Myo1 requires its motor activity (Huckaba et al., 2006). RACF, a widely conserved process, plays a critical role in the asymmetric inheritance of mitochondrial fitness in budding yeast, with the less oxidized mitochondria preferentially segregated into the daughter cell during bud growth (Higuchi et al., 2013). Whether Bni5 is involved in RACF via its interaction and recruitment of Myo1 at the bud neck has not been determined.

In mammalian cells, retrograde flow is driven by actin polymerization and myosin-II in the lamellipodia and lamella, respectively, to control the inward movement of T cell receptor microclusters at the immunological synapse (Babich et al., 2012; Yi et al., 2012), cell migration (Cai et al., 2006; Maiuri et al., 2015; Swaminathan et al., 2017), and growth cone dynamics (Lin et al., 1996; Medeiros et al., 2006). In all these systems, how the myosin-II is positioned in the lamella region and whether it drives the retrograde flow through the inward contraction of a concentric actomyosin ring, as hypothesized in the case of the immunological synapse (Yi et al., 2012), and/or by gliding on the flowing actin filaments as proposed in the case of cell migration (Gardel et al., 2010; Swaminathan et al., 2017) and growth cone dynamics (Lin et al., 1996; Medeiros et al., 2006) remains unknown. It is also unknown whether septins are involved in any of these retrograde systems. Given the geometric and molecular conservation (actin polymerization in the lamellipodia/at the bud tip and myosin-II in the lamella/at the bud neck) between the retrograde flow systems in budding yeast and mammalian cells, a deep mechanistic understanding of the RACF in budding yeast may inform similar analysis in other systems.

In this study, we have performed a comprehensive structure-function analysis on Bni5 using an integrative approach involving structure prediction by AlphaFold, gene editing, quantitative live-cell imaging, and biochemistry. This analysis has defined the specific domains of Bni5 necessary and sufficient for its interactions with the septin filaments, the septin-associated kinase Elm1 (Bouquin et al., 2000; Marquardt et al., 2020), and Myo1. Importantly, we have discovered three functions for Bni5. First, it regulates the timely remodeling of the septin hourglass to a double ring, which enables AMR constriction (Chen et al., 2020; Tamborrini et al., 2018). Second, Bni5 is responsible for the increase of Myo1 at the bud neck before the onset of cytokinesis, which strengthens the AMR against various genetic and chemical perturbations. Finally, Bni5 enhances RACF by binding to a specific region in Myo1 tail, which contributes to the asymmetric segregation of mitochondria-tethered protein aggregates that is related to cellular aging and health (Song et al., 2014; Zhou et al., 2014). Thus, we have established a specific biochemical pathway (septin-Bni5-Myo1) at the division site, which operates to control the efficiency, fidelity, and robustness of various cellular processes.

## Results

### C-terminal tagging of Bni5 compromises its septin-related function, timing of recruitment, and turnover kinetics at the division site

To understand how Bni5 tethers the dynamic myosin-II heavy chain (Myo1) to the stable septin hourglass before cytokinesis (Fang et al., 2010; Wloka et al., 2013), we must address an apparent conundrum: the endogenous Bni5 was previously shown to be co-purified stoichiometrically with the septin complexes suggesting a stable association (Mortensen et al., 2002; Renz et al., 2016), whereas fluorescence-recovery-after-photobleaching (FRAP) analyses using Bni5-C-GFP (C-terminally tagged) suggested a dynamic association with the septin hourglass (Schneider et al., 2013; Wloka et al., 2013). The C-, but not, N-terminal tagging of *BNI5* was shown to display a synthetic growth defect with a *cdc11* truncation allele, suggesting that the C-terminal tagging compromised Bni5 function (Finnigan et al., 2015a). This observation provided a clue to the problem and prompted us to further examine the positional impact of GFP tagging on Bni5 behavior and function. To this end, we constructed two plasmids to express *BNI5-C-GFP* or *BNI5-N-GFP* (N-terminally tagged) from the *MET17* promoter (induced by methionine depletion from the culture medium) in a *bni5Δ* strain to compare their localization and functionality (**Fig. 1 A**). We found that both tagged proteins, under the non-inducing condition, co-localized with septins (Cdc3-mCherry) at the bud neck (**Fig. 1 B**, left), as reported previously (Lee et al., 2002). Importantly, myosin-II (Myo1-mScarlet) localized at the bud neck of small- and medium-budded cells (G1/S/G2 stages) of both strains, in contrast to its loss of localization in *bni5Δ* strain (**Fig. 1 B**, right, arrowheads), suggesting a proper function of both tagged proteins as a septin-myosin-II linker. To examine their functionality further, we tested their ability as an overexpression suppressor of a temperature-sensitive septin mutant (*cdc12-6*) (Adams and Pringle, 1984; Lee et al., 2002) by introducing the plasmids into the *cdc12-6 bni5Δ* strain (**Fig. 1, C and D**). Strikingly, *BNI5-N-GFP*, under the inducing condition, was able to suppress the growth and morphological defects of *cdc12-6* at both the permissive (25°C) and the restrictive (32°C) temperatures to the same degree as the untagged *BNI5*. In contrast, *BNI5-C-GFP* was able to suppress the defects only at 25°C and behaved like the empty vector at 32°C (**Fig. 1, C and D**). These results indicate that C-terminal tagging of Bni5 does not overtly affect its localization or its role as a septin-myosin-II linker but compromises its function in septin regulation.

**Figure 1.**
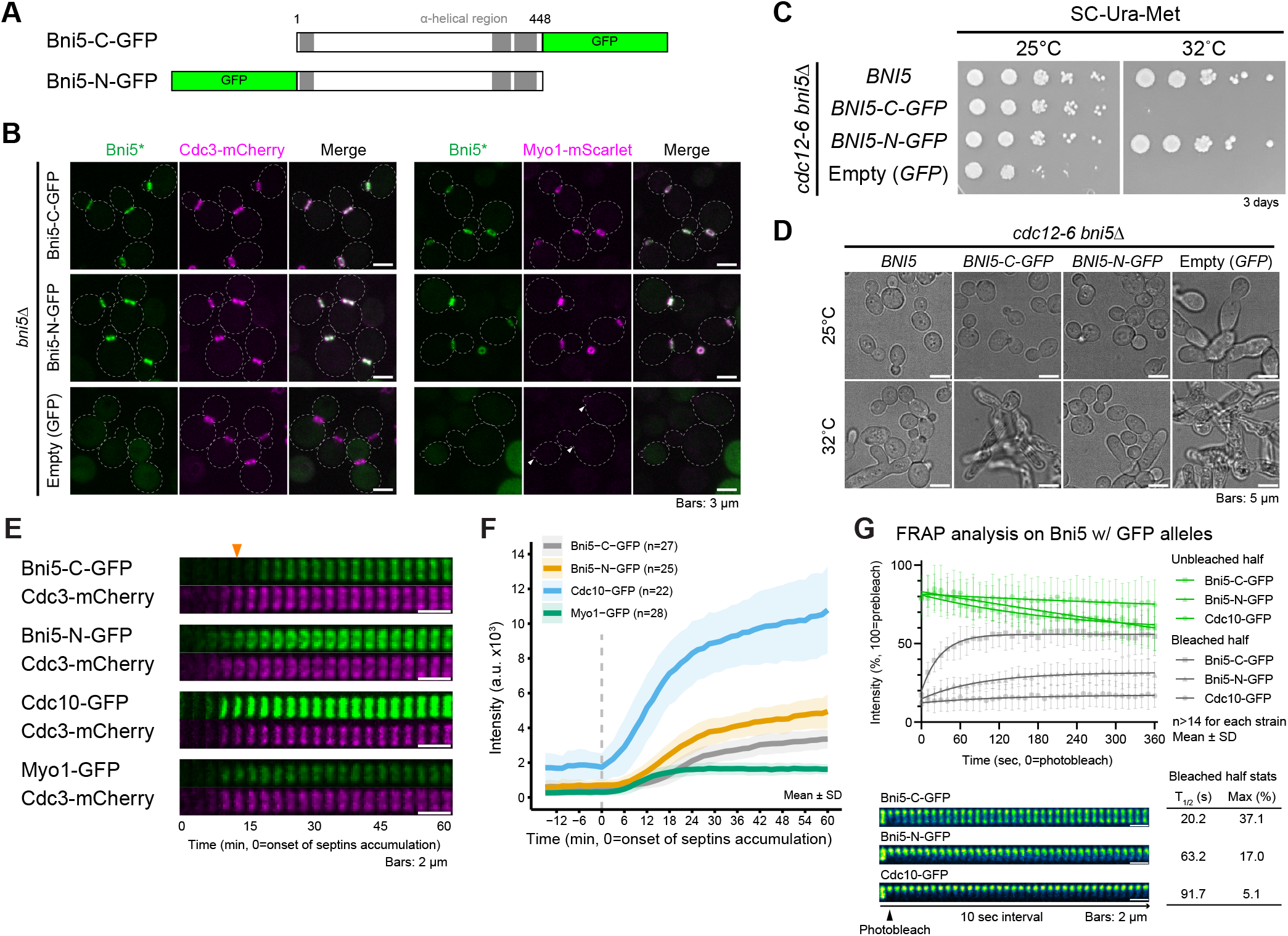
Distinct impact of N- and C-terminal GFP-tagging of Bni5 on its septin-related function, timing of recruitment, and turnover kinetics at the division site. (A) Diagram of Bni5-GFP constructs. Bni5-C-GFP and Bni5-N-GFP indicate GFP inserted at their C- and N-terminus, respectively. Numbers indicate amino acid residues. Gray boxes represent predicted α-helical regions. (B) Localization of Bni5-C-GFP and Bni5-N-GFP in relation to septins (Cdc3-mCherry) and myosin-II (Myo1-mScarlet). Cells cultured in SC-Ura at 25°C until the exponential growth phase were subjected to imaging. Asterisks represent variants of Bni5. Strains used are as follows: YEF11745 (*bni5Δ CDC3-mCherry* [*BNI5-C-GFP*]), YEF11053 (*bni5Δ CDC3-mCherry* [*BNI5-N-GFP*]), YEF11048 (*bni5Δ CDC3-mCherry* [*GFP*]), YEF11744 (*bni5Δ MYO1-mScarlet* [*BNI5-C-GFP*]), YEF11029 (*bni5Δ MYO1-mScarlet* [*BNI5-N-GFP*]), and YEF11033 (*bni5Δ MYO1-mScarlet* [*GFP*]). (C) Effect of induced expression of *BNI5* on the temperature sensitivity of *cdc12-6*. Cells with induced expression of different *BNI5* alleles were incubated for three days at 25°C or 32°C. Strains used are as follows: YEF11362 (*cdc12-6 bni5Δ* [*BNI5*]), YEF11363 (*cdc12-6 bni5Δ* [*BNI5-C-GFP*]), YEF11312 (*cdc12-6 bni5Δ* [*BNI5-N-GFP*]), and YEF11313 (*cdc12-6 bni5Δ* [*GFP*]). (D) Effect of induced expression of *BNI5* on cell morphology of *cdc12-6*. Cells harboring different alleles of *BNI5* were grown in SC-Ura-Met liquid medium to induce their expression at 25°C or 32°C for 16 hours before imaging. Strains used are the same as (C). (E) Time-lapse analysis of Bni5-C-GFP and Bni5-N-GFP accumulation at the budding site. Montages of GFP-tagged proteins with respect to Cdc3-mCherry were created from selected frames of time-lapse series taken with a 1.5-min interval. Arrowhead indicates the onset of Bni5-C-GFP accumulation. Strains used are as follows: YEF9336 (*BNI5-C-GFP CDC3-mCherry*), YEF10293 (*BNI5-N-GFP CDC3-mCherry*), YEF11750 (*CDC10-GFP CDC3-mCherry*), and YEF6108 (*MYO1-GFP CDC3-mCherry*). (F) Kinetics of the GFP-tagged proteins imaged in (E). Bold lines and associated shaded bands represent mean and SD values, respectively. (G) FRAP analysis of Bni5-C-GFP and Bni5-N-GFP to determine their turnover kinetics at the bud neck. Half of the ring from each cell was photo-bleached, and fluorescence recovery in the bleached region (gray symbol) and unbleached region (green symbol) was followed over time. Top: Lines, symbols, and error bars represent regression curves, mean values, and SD values, respectively. Bottom: Montages of cells were created from selected frames of time-lapse series taken with a 10-sec interval. Strains used are as follows: YEF11624 (*BNI5-C-GFP mScarlet-TUB1*), YEF11546 (*BNI5-N-GFP mScarlet-TUB1*), and YEF11753 (*CDC10-GFP mScarlet-TUB1*).

To gain further insights into the positional impact of GFP tagging on the functionality of Bni5, we constructed strains expressing either Bni5-C-GFP or Bni5-N-GFP from the native promoter at the endogenous locus and performed time-lapse imaging to acquire the localization kinetics (**Fig. 1, E and F**). Unless otherwise noted, all the kinetics analyzed in this study were obtained from GFP/RFP-tagged proteins that were chromosomally expressed by their own promoters, and different plots from each cell were aligned based on cell cycle-specific events, i.e., the start of the cell cycle in late G1 and the onset of cytokinesis in late anaphase/telophase, marked by the accumulation of septins at the budding site and the spindle breakage, respectively (Cid et al., 2001; Okada et al., 2021b; Woodruff et al., 2010). We found that Bni5-N-GFP arrived at the presumptive bud site at the same time as the septins (Cdc10-GFP and Cdc3-mCherry) and myosin-II (Myo1-GFP), but, strikingly, Bni5-C-GFP began to accumulate at the presumptive bud site ∼12 min after the septins, right around bud emergence (**Fig. 1, E and F**). Furthermore, Bni5-N-GFP accumulated at the bud neck ∼32% more than Bni5-C-GFP (intensities at 60 min: 4944 ± 545 vs. 3360 ± 981, **Fig. 1, E and F**). Thus, the C-terminal tagging affects both the timing and magnitude of Bni5 accumulation at the bud neck. Moreover, FRAP analysis revealed markedly reduced turnover of Bni5-N-GFP at the bud neck, which was similar to the septin (Cdc10-GFP) but in sharp contrast to the more dynamic Bni5-C-GFP (**Fig. 1 G**). The resemblance of Bni5-N-GFP with septins in recruitment timing and dynamics suggests that Bni5-N-GFP has a stronger association with septins than Bni5-C-GFP, thereby, explaining its efficient suppression of the septin mutant (*cdc12-6*) as well as its stable association with the septin complexes purified from yeast cells (Mortensen et al., 2002; Renz et al., 2016). Together, these data indicate that the C-terminal tagging of Bni5 compromises its septin-related function, timing of recruitment, and turnover kinetics at the division site.

### Bni5-C-GFP localizes to the division site via the septin hourglass-associated kinase Elm1

The delayed recruitment, dampened accumulation, and higher turnover of Bni5-C-GFP and its deficiency in septin function led us to hypothesize that the localization and/or maintenance of Bni5-C-GFP at the bud neck might be mediated by a dynamic septin-associated protein(s). The LKB1-like kinase Elm1, which specifically associates with the septin hourglass, appeared to be an attractive candidate, as it localized to the bud neck around bud emergence, similar to Bni5-C-GFP (**Fig. 2 A cf. Fig. 1 E**) (Bouquin et al., 2000; Lee et al., 2002; Marquardt et al., 2020). In addition, Elm1 is known to interact with Bni5 (Marquardt et al., 2020; Patasi et al., 2015).

**Figure 2.**
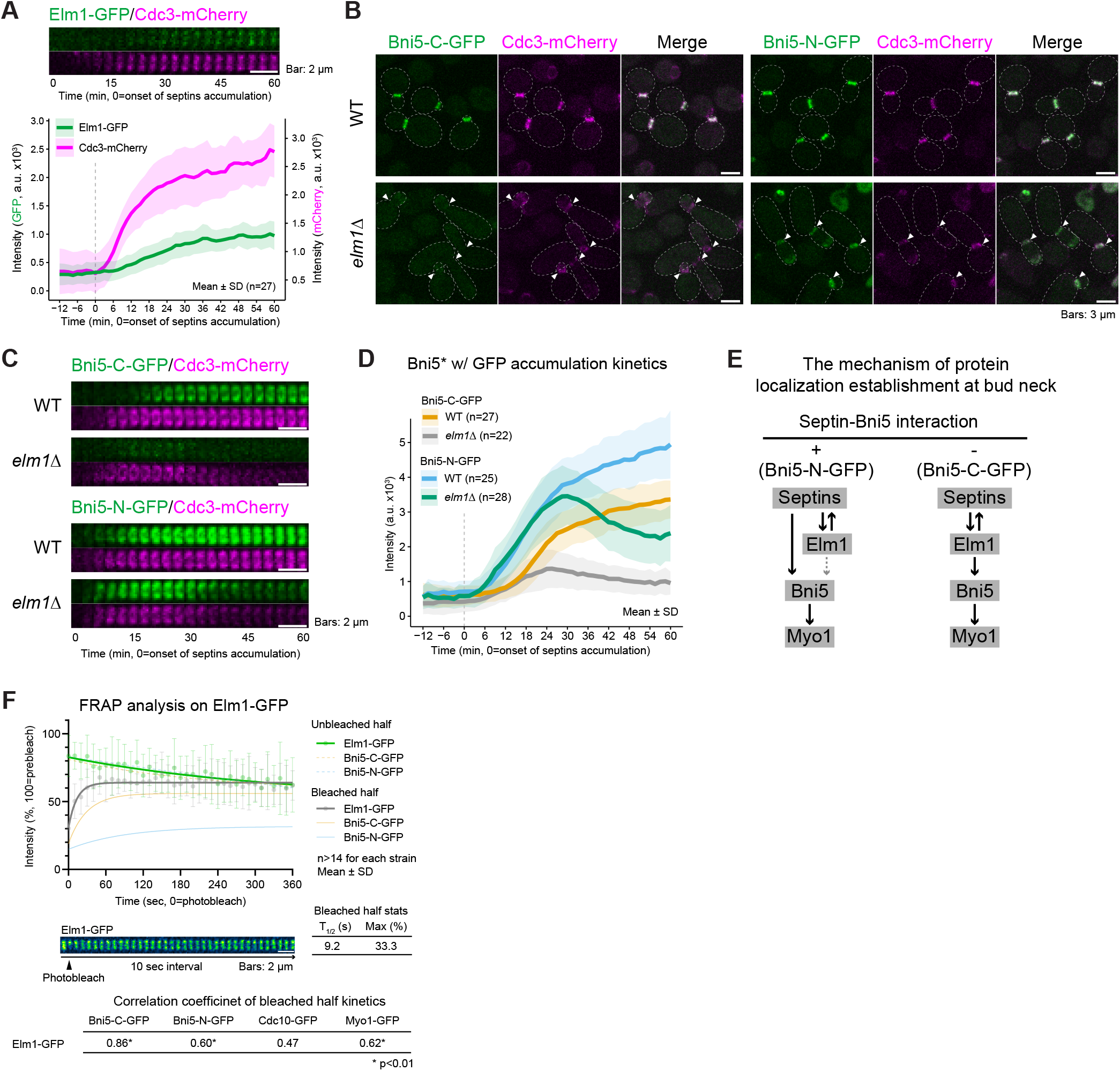
Bud neck localization of Bni5-C-GFP depends on the septin hourglass-associated kinase Elm1. (A) Elm1-GFP accumulation at the budding site. Top: Montages of Elm1-GFP with respect to Cdc3-mCherry were created from selected frames of time-lapse series taken with a 1.5-min interval. Bottom: Kinetics of the Elm1-GFP and Cdc3-mCherry. Bold lines and associated shaded bands represent mean and SD values, respectively. Strain used is YEF10440 (*ELM1-GFP CDC3-mCherry*). (B) Localization of Bni5-C-GFP and Bni5-N-GFP in *elm1Δ* cells. Cells cultured in SC-Ura at 25°C until the exponential growth phase were subjected to imaging. Strains used are as follows: YEF9336 (*BNI5-C-GFP CDC3-mCherry*), YEF9369 (*elm1Δ BNI5-C-GFP CDC3-mCherry*), YEF10293 (*BNI5-N-GFP CDC3-mCherry*), and YEF10315 (*elm1Δ BNI5-N-GFP CDC3-mCherry*). Arrowheads indicate bud necks determined by Cdc3-mCherry localization. (C) Time-lapse analysis of Bni5-C-GFP and Bni5-N-GFP accumulation at the budding site in *elm1Δ* cells. Montages of GFP-tagged Bni5 with respect to Cdc3-mCherry were created from selected frames of time-lapse series taken with a 1.5-min interval. Strains used are the same as in (B). See also **Figure S1**. (D) Kinetics of the GFP-tagged proteins imaged in (C). Bold lines and associated shaded bands represent mean and SD values, respectively. An asterisk represents variants of Bni5. See also **Figure S1**. (E) A model for the bud neck localization of Bni5-N-GFP and Bni5-C-GFP. Solid and dashed arrows represent high- and low-confidence localization dependencies, respectively. For example, the arrow from Bni5 to Myo1 indicates that Bni5 is the localization determinant for Myo1, supported by in vivo and in vitro evidence. (F) FRAP analysis on Elm1-GFP. Half of the ring from each cell was photo-bleached, and fluorescence recovery in the bleached region (gray symbol) and unbleached regions (green symbol) was followed over time. Top: Lines, symbols, and error bars represent regression curves, mean values, and SD values, respectively. The reference plots of Bni5-C-GFP and Bni5-N-GFP were modified from Figure 1G. Middle: Montages of cells were created from time-lapse series taken with a 10-sec interval. Strain used is YEF8437 (*ELM1-GFP mRuby2-TUB1*). Bottom: Correlation coefficients of bleached half kinetics between Elm1-GFP and other proteins. See also **Figure S1**.

To test this hypothesis, we examined the localization of Bni5-C-GFP and Bni5-N-GFP expressed from plasmids in *elm1*Δ cells and found that Bni5-C-GFP was absent at both the bud neck and the bud tip, despite the presence of septins (Cdc3-mCherry) at both locations – a stereotypical phenotype for *elm1*Δ cells (**Fig. 2 B**, white arrowheads) (Marquardt et al., 2020). In contrast, Bni5-N-GFP co-localized with the septins at both locations (**Fig. 2 B**, white arrowheads). Time-lapse analysis of both Bni5-C-GFP and Bni5-N-GFP expressed from the native promoter at the endogenous locus confirmed this observation, and additionally showed that the peak intensity of Bni5-C-GFP in *elm1*Δ cells was significantly reduced [61% of the peak intensity in wild-type (WT), time = 25.5 min, p < 0.01] whereas the peak intensity of Bni5-N-GFP was virtually unchanged (91% of that in WT, time = 30 min, p = 0.13) (**Fig. 2, C and D**). These data indicate that the neck localization of Bni5-C-GFP, but not Bni5-N-GFP, largely depends on Elm1. As expected, septin accumulation at the bud neck in *elm1*Δ cells peaked at ∼30 min and then decreased due to the migration of destabilized septins from the bud neck to the bud tip (**Fig. S1 A)** (Marquardt et al., 2020). Importantly, septin accumulation in Bni5-C-GFP-expressing WT and *elm1*Δ cells remained unchanged until the peak (25.5 min, **Fig. S1 A**), which demonstrates that the decreased localization of Bni5-C-GFP in *elm1*Δ cells is not due to decreased septin localization at the bud neck. The simplest interpretation for these results is that C-terminal tagging disrupts the Bni5-septin interaction but not the Bni5-Elm1 interaction, and that Bni5-N-GFP, presumably like the untagged Bni5, is targeted to the division site mainly via the Bni5-septin interaction. However, when this interaction is compromised, as is apparently the case for Bni5-C-GFP, the Bni5-Elm1 interaction becomes essential for its localization to the bud neck (**Fig. 2 E**).

To test our hypothesis further, we performed FRAP analysis on Elm1-GFP, anticipating that the dynamic characteristics of Bni5-C-GFP (**Fig. 1 G**) reflect the dynamic characteristics of its localization determinant, Elm1 (**Fig. 2 E**). Indeed, Elm1 was dynamic and showed the highest correlation with Bni5-C-GFP (r = 0.86) among other components examined (**Fig. 2 F, and Fig. S1, B and C**). In contrast, septins, the localization determinant for Bni5-N-GFP (**Fig. 2 E**), showed the highest correlation with Bni5-N-GFP (Cdc10-GFP, r = 0.88, **Figs. S1 C**).

Collectively, these analyses reveal a close association of Bni5-N-GFP and Bni5-C-GFP with the septins and Elm1, respectively, and demonstrate an essential role of Elm1 in targeting Bni5-C-GFP to the division site.

### Bni5 interacts with myosin-II and septins via distinct regions

To determine how Bni5 functions as the septin-myosin-II linker at the molecular level, we conducted a domain analysis on Bni5 to identify the essential regions for its interactions with myosin-II and septins, respectively. As the crystal structure of Bni5 is not available, we used its AlphaFold-predicted structure (Jumper et al., 2021; Varadi et al., 2022) to guide our analysis (**Fig. 3 A**, right). Bni5 is predicted to consist of three alpha-helical regions that we designated as CC1 (<u>c</u>oiled-<u>c</u>oil #1, aa5-33), CC2 (aa355-390), and CC3 (aa396-436). Additionally, a long-disordered region (aa 41-339), as predicted by IUPred3 (Erdos et al., 2021), exists between CC1 and CC2. Sequence analysis by BLAST showed that the CC1-3, but not the disordered region, are specifically conserved among budding yeast species (**Fig. S 2**).

**Figure 3.**
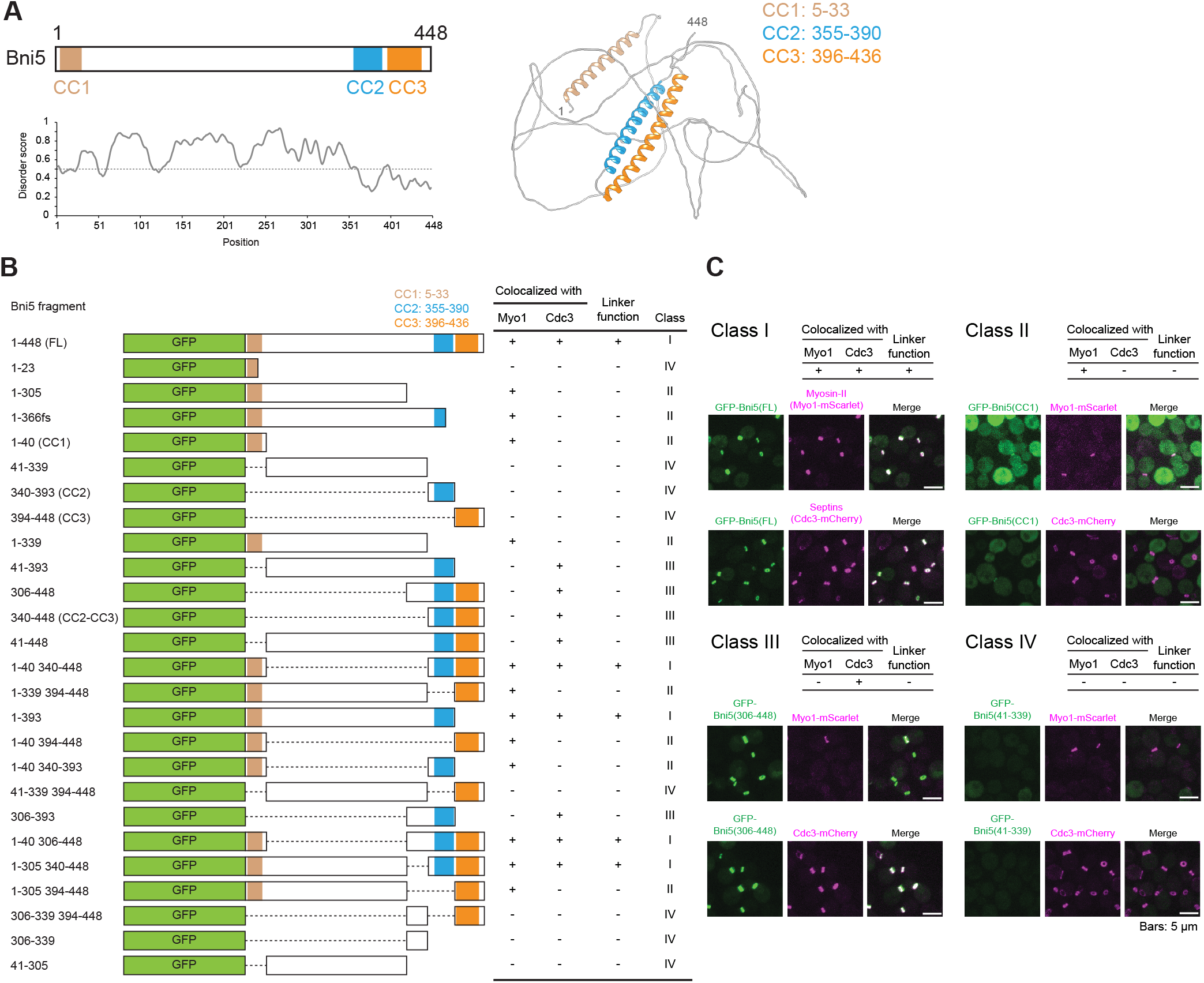
Bni5 interacts with myosin-II and septins via distinct domains. (A) Diagram of the molecular structure of Bni5. Top left: Secondary structure of Bni5. Brown, blue, and orange boxes represent coiled-coil domains designated CC1, CC2, and CC3, respectively. Bottom left: Result of IUPred3 prediction of domain disorder. The score above the dotted line indicates the higher probability of each residue being disordered. Right: AlphaFold model of Bni5. The CC1, CC2, and CC3 are highlighted as the same color code used in the top left panel. (B) Summary of colocalization between different Bni5 fragments and myosin-II or septins. Localization of different fragments of Bni5 is analyzed using the pUG36-Bni5* plasmid series (**Table S2**). Boxes and dashed lines indicate the presence and absence of the region of Bni5 in relevant alleles. In the first two columns, + and – signs indicate the presence and absence of colocalization with Myo1-mScarlet and Cdc3-mCherry. In the third column, + and – signs indicate the presence and absence of the linker function, i.e., + means colocalization with both Myo1 and Cdc3 is observed in relevant alleles. In the fourth column, the class is determined based on the colocalization patterns described in (C). Strains used are summarized in **Table S1**, 56 strains with denotation with ^p^ and ^r^. (C) Class and pattern of colocalization of Bni5 with myosin-II or septins. Class I includes Bni5 fragments that colocalize with both myosin-II and septins. Class II or Class III includes Bni5 fragments that colocalize with only myosin-II or septins, respectively. Class IV includes Bni5 fragments that did not colocalize with myosin-II and septins. Strains selected as class-representative are as follows: Class I: YEF11029 (*bni5Δ MYO1-mScarlet* [*GFP-BNI5(FL)*] and YEF11053 (*bni5Δ CDC3-mCherry* [*GFP-BNI5(FL)*], Class II: YEF11058 (*bni5Δ MYO1-mScarlet* [*GFP-bni5(1-40)*] and YEF11038 (*bni5Δ CDC3-mCherry* [*GFP-bni5(1-40)*]), Class III YEF11020 (*bni5Δ MYO1-mScarlet* [*GFP-bni5(306-448)*]) and YEF11039 (*bni5Δ CDC3-mCherry* [*GFP-bni5(306-448)*], and Class IV: YEF11025 (*bni5Δ MYO1-mScarlet* [*GFP-bni5(41-339)*] and YEF11044 (*bni5Δ CDC3-mCherry* [*GFP-bni5(41-339)*]).

We constructed a set of plasmids carrying different fragments of Bni5 with GFP fused to their N-termini (**Fig. 3 B**). These GFP-Bni5 constructs were expressed in *bni5*Δ strains carrying either Myo1-mScarlet or Cdc3-mCherry and subjected to live-cell imaging analysis. Based on their colocalization patterns with Myo1 and/or Cdc3, we classified the GFP-Bni5 fragments into four categories (Class I-IV, **Fig. 3 B and C**). Class I showed colocalization with both Myo1 and Cdc3, indicating that these fragments are sufficient for binding to both targets and serve as a molecular linker (**Fig. 3 C**). In contrast, Class II and III showed colocalization with either Myo1 or Cdc3, respectively, suggesting that they retain their ability to bind to their specific target (**Fig. 3 C**). Class IV did not colocalize with Myo1 and Cdc3, indicating a loss of their ability to bind either target and/or a defect in their expression (**Fig. 3 C**).

Remarkably, all fragments that exhibited colocalization with Myo1 (Class I and II) contained intact CC1 (e.g., aa1-305, class II, **Fig. 3 B**), whereas none of the fragments that lost partial (aa1-23) or entire CC1 (e.g., aa41-448) was able to colocalize with Myo1 (**Fig. 3 B**). Similarly, all fragments that colocalized with Cdc3 (Class I and III) contained intact CC2 (e.g., aa41-393, class III, **Fig. 3 B**), whereas none of the fragments that lost partial (aa1-366fs) or entire CC2 (e.g., aa1-339) colocalized with Cdc3 (**Fig. 3 B**). These results strongly suggest that CC1 and CC2 are the essential functional domains for myosin-II and septin interaction, respectively. The presence or absence of the disordered region did not affect the colocalization patterns (e.g., FL vs. 1-40 340-448, both class I, **Fig. 3 B**), suggesting that the disordered region does not play a significant role in the binding of Bni5 to either Myo1 or septins.

### The CC1 domain of Bni5 is necessary and sufficient for its interaction with Myosin-II

To further define the role of CC1 in myosin-II binding, we first determined its role in Myo1 recruitment during bud formation by time-lapse analysis (**Fig. 4, A and B**). In WT cells, GFP-Bni5(FL) (full length, equivalent of Bni5-N-GFP) and Myo1-mScarlet co-localized as a ring at the bud neck. As expected, in *bni5Δ* cells, the neck localization of Myo1-mScarlet was abolished (Fang et al., 2010). Strikingly, in cells expressing GFP-Bni5-(CC1)Δ, Myo1-mScarlet failed to localize to the bud neck as seen in *bni5Δ* cells, despite its own normal accumulation at the bud neck (**Fig. S3 A**). These results indicate that CC1 is required for Myo1 binding but not for its localization at the bud neck.

**Figure 4.**
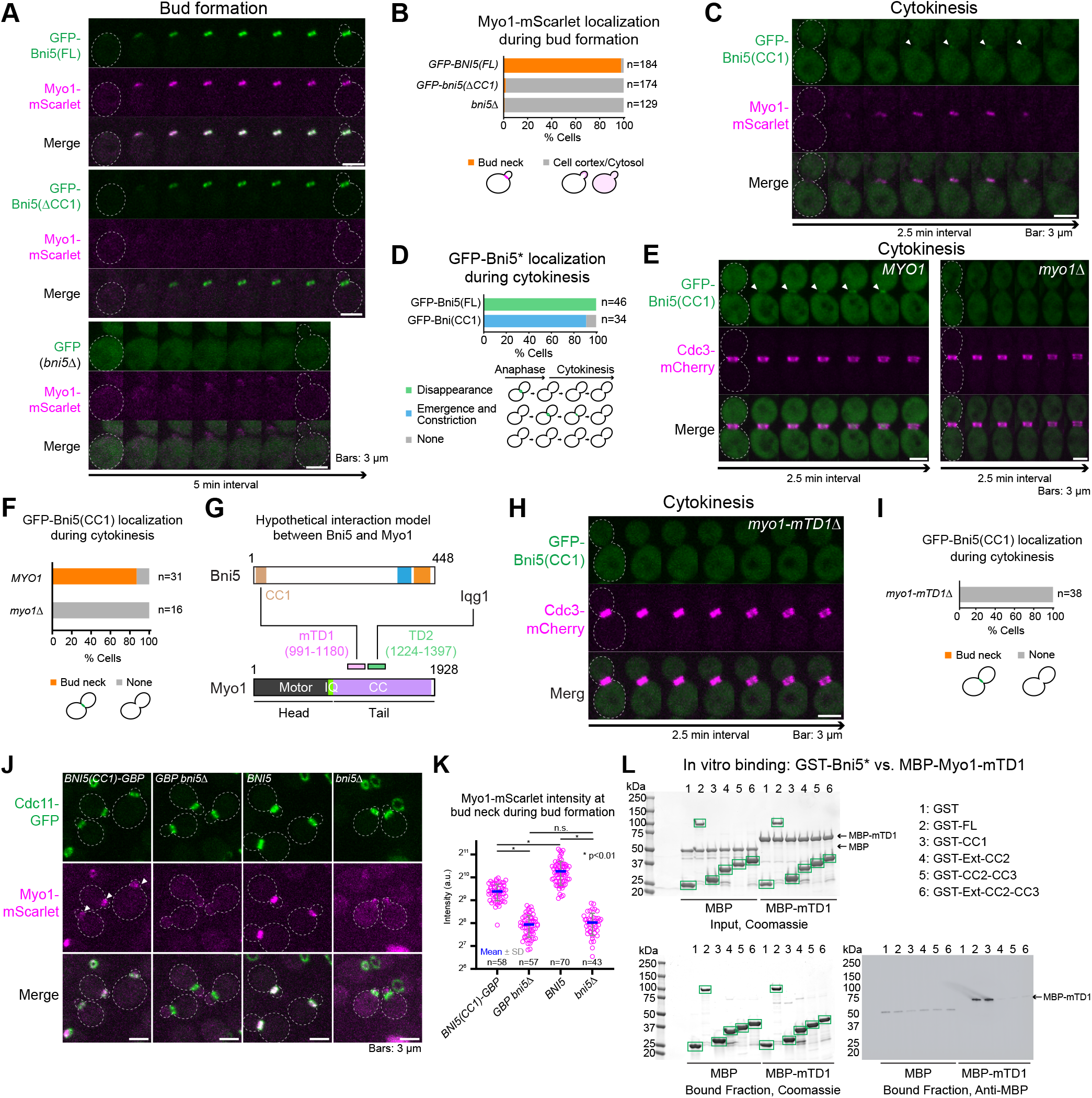
The CC1 domain of Bni5 is necessary and sufficient for binding to a targeting domain in Myosin-II. (A) Myo1 accumulation at the budding site depends on the CC1 of Bni5. Montages of GFP-tagged Bni5 fragments with respect to Myo1-mScarlet were created from selected frames of time-lapse series taken with a 2.5-min interval. Strains used are as follows: YEF11029 (*bni5Δ MYO1-mScarlet* [*GFP-BNI5(FL)*], YEF11027 (*bni5Δ MYO1-mScarlet* [*GFP-bni5(41-448)*], and YEF11033 (*bni5Δ MYO1-mScarlet* [*GFP*]). See also **Figure S4**. (B) Bud neck localization of Myo1 in *bni5(ΔCC1)* cells. Myo1 localization was scored from the imaging data in (A). (C) Bni5(CC1) localization at the division site during cytokinesis. Montages of GFP-Bni5(CC1) with respect to Myo1-mScarlet were created from time-lapse series taken with a 2.5-min interval. Strain used is YEF11058 (*bni5Δ MYO1-mScarlet* [*GFP-bni5(1-40)*]). (D) Distinct patterns of bud neck localization of Bni5 FL vs its CC1 fragment. Localization of GFP-Bni5(FL) and GFP-Bni5(CC1) at the bud neck during anaphase and cytokinesis was scored from the imaging data in (C). See also **Figure S4**. (E) The bud neck localization of Bni5(CC1) depends on Myo1. Montages of GFP-Bni5(CC1) with respect to Cdc3-mCherry in WT and *myo1Δ* cells were created from time-lapse series taken with a 2.5-min interval. Strains used are as follows: YEF11038 (*bni5Δ CDC3-mCherry* [*GFP-bni5(1-40)*] and YEF11114 (*myo1Δ CDC3-mCherry* [*GFP-bni5(1-40)*]). (F) Quantitative analysis of Bni5(CC1) localization at the bud neck in WT and *myo1Δ* cells. The imaging data in (E) were used for this analysis. (G) Diagram of putative and demonstrated interactions among Bni5, Myo1, and Iqg1. Minimal targeting domain 1 (mTD1) and targeting domain 2 (TD2) as the binding sites for Bni5 and Iqg1, respectively, were determined in (Fang et al., 2010). Motor, IQ, and CC represent the motor domain, IQ motifs, and coiled-coil domain, respectively. (H) Time-lapse analysis of Bni5(CC1) localization at the bud neck in *myo1-mTD1Δ* cells. Montages were created from time-lapse series taken with a 2.5-min interval. Strain used is YEF11757 (*myo1-mTD1Δ CDC3-mCherry* [*GFP-bni5(1-40)*]). (I) Quantitative analysis of Bni5(CC1) localization at the bud neck in *myo1-mTD1Δ* cells. The imaging data in (H) were used for this analysis. See also **Figure S4**. (J) Forced tethering of Bni5(CC1) to the septin hourglass restores Myo1 localization at the bud neck. Cells in the exponential growth phase were subjected to imaging. Strains used are as follows: YEF11212 (*CDC11-GFP MYO1-mScarlet bni5(1-40)-GBP*), YEF11211 (*CDC11-GFP MYO1-mScarlet bni5Δ::GBP*), YEF11088 (*CDC11-GFP MYO1-mScarlet*), and YEF11248 (*CDC11-GFP MYO1-mScarlet bni5Δ*). (K) Quantitative analysis of the Bni5(CC1)-restored Myo1 at the bud neck. Myo1-mScarlet intensity at the bud neck in cells with small- and medium-bud (S/G2 cells) was scored from the imaging data in (J). Magenta circles, blue bars, and gray error bars represent all data points, mean values, and SD values, respectively. See also **Figure S4**. (L) Bni5 binds to the mTD1 of Myo1 via CC1. Top left: SDS-PAGE stained with Coomassie Blue depicting amounts of each indicated purified protein used as the input for the in vitro binding assay. Bottom left: In vitro binding assay results (Coomassie Blue stained gel) for the indicated GST-tagged proteins bound to glutathione resin and their ability to pull down MBP-mTD1. Green boxes represent GST and GST-tagged Bni5 fragments as described on the top right. Bottom right: immunoblotted membrane with antibody against MBP; kDa = kiloDalton. This experiment was repeated three times, and a representative immunoblot is shown.

We then examined the localization of the GFP-CC1 fragment during the cell cycle and found that it localized in the cytoplasm before cytokinesis and accumulated as a weak signal and constricted with Myo1-mScarlet at the bud neck during cytokinesis (**Fig. 4 C**, arrowheads). This pattern of localization is in sharp contrast to that of the full-length Bni5, which was rapidly removed from the division site before cytokinesis (**Fig. 4 D and Fig. S3, B and C**). The weak signal of GFP-CC1 accounted for only 14% of the maximum value of GFP-Bni5 (**Fig. S3 C**). As expected, Myo1 localization during cytokinesis was normal in *bni5Δ* as this phase of Myo1 localization is mediated by Iqg1 and not Bni5 (**Fig. S3 B**) (Fang et al., 2010). Strikingly, the localization of GFP-CC1 during cytokinesis was not observed in *myo1Δ* cells (**Fig. 4, E and F**), suggesting that CC1 is sufficient for Myo1 binding.

The fact that the full-length Bni5 interacts with the minimal targeting domain 1 (mTD1: aa991-1180) of Myo1 in vitro (**Fig. 4 G**) (Fang et al., 2010) raises the possibility that CC1 may interact with mTD1 (**Fig. 4 G**). To explore this possibility, we first tested the requirement of mTD1 for the neck localization of GFP-CC1 during cytokinesis and found that GFP-CC1 signal at the bud neck was abolished in *myo1-mTD1Δ* cells (**Fig. 4, H and I**). Reciprocally, we tested the requirement of CC1 for the neck localization of Myo1-mTD1-GFP, which is known to be entirely dependent on Bni5 (Fang et al., 2010). As expected, Myo1-mTD1-GFP failed to localize to the bud neck in *bni5Δ* cells, but this defect was rescued by a plasmid expressing *BNI5* (**Fig. S3 D**). However, the rescue was not observed with a plasmid expressing the CC1-deleted *BNI5* (*bni5-CC1Δ*) (**Fig. S3 D**). We also used GBP (GFP-binding protein) (Rothbauer et al., 2008) to forcibly tether CC1 to the septins (Cdc11-GFP) (**Fig. S3 E**). Strikingly, this septin-tethered CC1 was able to recruit Myo1-mScarlet to the bud neck during bud formation (**Fig. 4 J**, arrowheads, **and Fig. 4 K**). Finally, recombinant Bni5-CC1 and Myo1-mTD1 interacted directly in vitro (**Fig. 4 L**). Taken together, these data demonstrate that the CC1 domain of Bni5 is necessary and sufficient for its binding to the mTD1 of Myo1.

### CC2 and its adjacent regions in Bni5 interact with the septins and Elm1 to control its localization and septin-related functions at the division site

Our analysis revealed that the CC2 region was required for the colocalization of Bni5 with septins (**Fig. 3 B and Fig. 5, A and B**), suggesting its possible involvement in septin binding. However, CC2 alone or its adjacent regions by themselves did not show any neck localization under the normal expression condition (**Fig. 5 A and Fig. S4 A**). In contrast, combining CC2 (aa340-393) with its upstream and/or downstream adjacent regions, which we designated as Ext (<u>ext</u>ended sequence required for CC2 localization, aa306-339) and CC3 (aa394-448), respectively, was sufficient for colocalization with the septin (Cdc3-mCherry) (**Fig. 5, A and B**). Surprisingly, when overexpressed under the induced condition, Ext localized weakly to the bud neck (**Fig. S4 B**, arrowheads), whereas CC2 was exclusively cytosolic, and CC3 did not show an enhanced signal compared to the uninduced condition, suggesting poor expression or rapid degradation (**Fig. S4 B**). Taken together, these data indicate that Ext and CC3 play distinct roles in establishing the CC2 neck localization. Indeed, our time-lapse analysis showed that Ext-CC2 co-appeared with the septins at the budding site (**Fig. 5 C**), similar to Bni5-N-GFP (**Fig. 1 E**), whereas CC2-CC3 exhibited a delay in bud neck recruitment during bud formation (**Fig. 5 C**, arrowhead), similar to Bni5-C-GFP (**Fig. 1 E,** arrowhead), whose neck localization depended on Elm1 (**Fig. 2, B-D**). These observations suggest that Ext-CC2 may bind to the septins whereas CC2-CC3 may bind to Elm1. This conclusion is supported by the observation that the neck localization of overexpressed Ext depended on the septins (Cdc11 and Shs1) but not Elm1 (**Fig. S4 B**, arrowheads) whereas the neck localization of CC2-CC3, but not Ext-CC2, almost entirely depended on Elm1 (**Fig. 5, D and E**). Using recombinant proteins, we found that both Ext-CC2 and CC2-CC3 interacted with Elm1-C-terminus directly in vitro (**Fig. 5 F,** lanes 4 and 5). Taken together, these data suggest that Ext-CC2 localize to the division site by interacting with both septins and Elm1 whereas CC2-CC3 do so by interacting with Elm1 mainly, if not exclusively.

**Figure 5.**
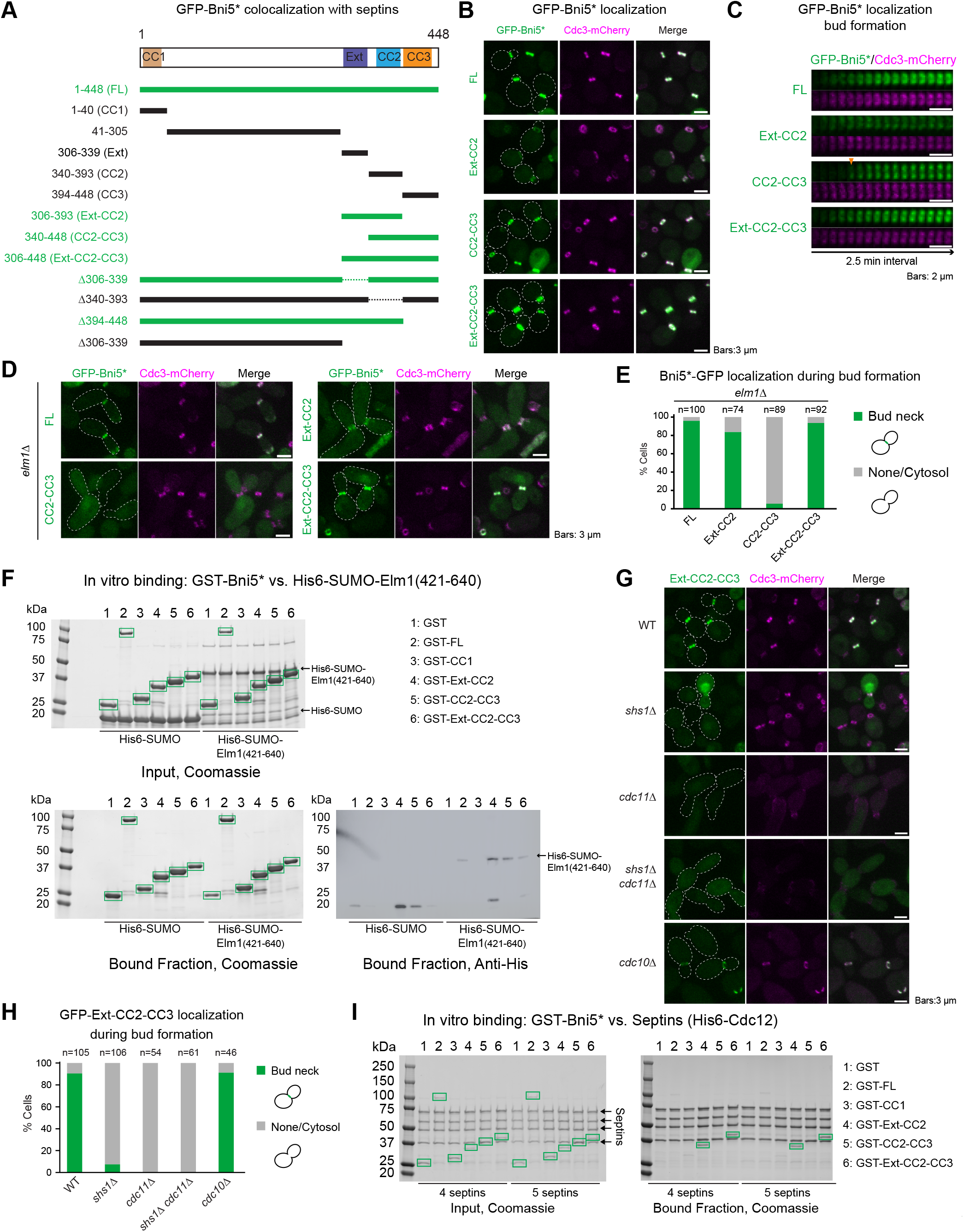
The CC2 domain and adjacent regions of Bni5 interact with septin filaments and Elm1. (A) Summary of colocalization between different Bni5 fragments and septins. Selected and modified data from the results of Figure 3B are presented. Alleles in green and black indicate colocalization and non-colocalization with septins (Cdc3-mCherry), respectively. Bold lines and dashed lines indicate the presence and absence of the region of Bni5 in relevant alleles. An asterisk represents variants of Bni5. Strains used are as follows: YEF11053 (*bni5Δ CDC3-mCherry* [*GFP-BNI5(FL)*]), YEF11038 (*bni5Δ CDC3-mCherry* [*GFP-bni5(1-40)*]), YEF11521 (*bni5Δ CDC3-mCherry* [*GFP-bni5(41-305)*]), YEF11181 (*bni5Δ CDC3-mCherry* [*GFP-bni5(306-339)*]), YEF11041 (*bni5Δ CDC3-mCherry* [*GFP-bni5(340-393)*]), YEF11042 (*bni5Δ CDC3-mCherry* [*GFP-bni5(394-448)*]), YEF11182 (*bni5Δ CDC3-mCherry* [*GFP-bni5(306-393)*]), YEF11040 (*bni5Δ CDC3-mCherry* [*GFP-bni5(340-448)*]), YEF11039 (*bni5Δ CDC3-mCherry* [*GFP-bni5(306-448)*]), YEF11184 (*bni5Δ CDC3-mCherry* [*GFP-bni5(Δ306-339)*]), YEF11049 (*bni5Δ CDC3-mCherry* [*GFP-bni5(Δ340-393)*]), YEF11051 (*bni5Δ CDC3-mCherry* [*GFP-bni5(1-393)*]), and YEF11055 (*bni5Δ CDC3-mCherry* [*GFP-bni5(1-305)*]). (B) Colocalization of indicated Bni5 fragments with septins at the bud neck. Cells in the exponential growth phase were subjected to imaging. Asterisks represent variants of Bni5. Strains used are as follows: YEF11053, YEF11181, YEF11040, and YEF11039. (C) Time-lapse analysis of colocalization between Bni5 fragments and septins at the bud neck. Montages of Bni5 fragments with respect to Cdc3-mCherry were created from time-lapse series taken with a 2.5-min interval. Arrowhead indicates the onset of GFP-CC2-CC3 localization at the budding site. An asterisk represents variants of Bni5. Strains used are the same as in (B). (D) Localization of Bni5 fragments in *elm1Δ* cells. Cells in the exponential growth phase were subjected to imaging. Strains used are as follows: YEF11190 (b*ni5Δ elm1Δ CDC3-mCherry* [*GFP-BNI5(FL)*]), YEF11197 (*bni5Δ elm1Δ CDC3-mCherry* [*GFP-bni5(306-393)*]), YEF11194 (*bni5Δ elm1Δ CDC3-mCherry* [*GFP-bni5(340-448)*]), and YEF11193 (*bni5Δ elm1Δ CDC3-mCherry* [*GFP-bni5(306-448)*]). (E) Quantitative analysis of Bni5 fragment localization in *elm1Δ* cells during bud formation. Localization of the indicated GFP-Bni5 fragments in *elm1Δ* cells with a small or medium bud (S/G2 cells) was scored from the imaging data in (D). An asterisk represents variants of Bni5. (F) The CC2-containing fragments of Bni5 interact with Elm1. Top left: SDS-PAGE stained with Coomassie Blue depicting amounts of each indicated purified protein used as the input for the in vitro binding assay. Bottom left: In vitro binding assay results (Coomassie Blue stained gel) for the indicated GST-tagged proteins bound to glutathione resin and their ability to pull down His-SUMO-Elm1(421-640). Green boxes represent GST and GST-tagged Bni5 fragments as described on the top right. Bottom right: immunoblotted membrane with antibody against His; kDa = kiloDalton. An asterisk represents variants of Bni5. This experiment was repeated three times, and a representative immunoblot is shown. (G) Localization of Bni5(Ext-CC2-CC3) in different septin mutants. Cells in the exponential growth phase were subjected to imaging. Strains used are as follows: YEF11039 (*bni5Δ CDC3-mCherry* [*GFP-bni5(306-448)*]), YEF11121 (*shs1Δ CDC3-mCherry* [*GFP-bni5(306-448)*]), YEF11310 (*cdc11Δ CDC3-mCherry* [*GFP-bni5(306-448)*]), YEF11790 (*shs1Δ cdc11Δ CDC3-mCherry* [*GFP-bni5(306-448)*]), and YEF11117 (*cdc10Δ CDC3-mCherry* [*GFP-bni5(306-448)*]). (H) Quantitative analysis of Bni5(Ext-CC2-CC3) localization in different septin mutants. Localization of the indicated GFP-Bni5 fragments in cells with a small or medium bud (S/G2 cells) was scored from the imaging data in (G). (I) Bni5 binds to septin filaments via Ext-CC2. Left: SDS-PAGE stained with Coomassie Blue depicting the amount of each indicated purified protein used as the input for the in vitro binding assay. Right: In vitro binding assay results (Coomassie Blue stained gel) for indicated GST-tagged proteins co-sedimented with septin filaments by ultracentrifugation; kDa = kiloDalton. Green boxes represent GST-tagged proteins. An asterisk represents variants of Bni5. This experiment was repeated three times, and a representative immunoblot is displayed. See also **Figure S4**.

At the functional level, we found that overexpression of Ext-CC2 or CC2-CC3 did not suppress the temperature sensitivity of the *cdc12-6* strain, whereas overexpression of a fragment containing all three regions (Ext-CC2-CC3, aa306-448) displayed a strong suppression (**Fig. S4 C**). Thus, all three regions of Bni5 are required for its role in septin regulation.

The ability of Ext-CC2-CC3 to suppress the septin mutant (*cdc12-6*) (**Fig. S4 C**) and its concomitant recruitment with Cdc3-mCherry at the budding site (**Fig. 5 C**) raise the possibility that Ext-CC2-CC3 may define the “minimal domains” for septin binding and function. Bni5 is thought to interact with the terminal subunit(s) of septin octamers (Booth et al., 2016; Finnigan et al., 2015a; Finnigan et al., 2016; Lee et al., 2002). Thus, we expressed Ext-CC2-CC3 in the septin terminal subunit deletion strains (*shs1Δ, cdc11Δ,* and *shs1Δ cdc11Δ*) and found that more than 90% of *shs1Δ* cells and 100% of *cdc11Δ* and *shs1Δ cdc11Δ* cells lost the bud neck signal (**Fig. 5, G and H**). In contrast, the localization of Ext-CC2-CC3 was unchanged in a core septin subunit deletion strain (*cdc10Δ*). These results suggest that Ext-CC2-CC3 accounts for the localization dependency of Bni5 on the septin terminal subunits. To test this hypothesis, we directly tested interaction between recombinant Ext-CC2-CC3 and septin filaments made of either Cdc11-capped octamers (4 septins) or both Cdc11- and Shs1-capped octamers (5 septins) (**Fig. 5 I and Fig. S4 D**). Because Bni5 is known to interact only with septin filaments (Booth et al., 2016; Patasi et al., 2015; Renz et al., 2013) and Shs1-capped octamers alone do not form filaments in vitro (Garcia et al., 2011), we did not include a reaction containing the Shs1-capped octamers alone. Surprisingly, Ext-CC2-CC3 showed an equally strong interaction with the septin filaments regardless of the presence of the Shs1 (**Fig. 5 I and Fig. S4 D**), suggesting that Cdc11 could be the direct target of Bni5 whereas Shs1 may play a regulatory role in vivo (see next paragraph). We also found that Ext-CC2, but not CC2-CC3 or CC1, interacted robustly with the septin filaments, similar to Ext-CC2-CC3 (**Fig. 5 I and Fig. S4 D)**. This result is in remarkable agreement with our in vivo localization data and highlights the Ext as a key determinant for septin interaction.

Given the in vitro interaction data, it remains a mystery why Ext-CC2-CC3 failed to localize in *shs1Δ* cells (**Fig. 5 G**), despite the presence of Cdc11-containing septin filaments at the bud neck (**Fig. S4 E**). One possible explanation is that Ext-CC2-CC3 may interact with Shs1 first in order to generate a confirmation that is competent for its interaction with Cdc11. Alternatively, Ext-CC2-CC3 may favor a heterotypic interface (Cdc11-Shs1) formed by an end-to-end interaction of Shs1- and Cdc11-capped octamers, which likely exists in vivo (Booth et al., 2015; Finnigan et al., 2015b). The loss of either terminal subunit leads to the formation of a homotypic interface (Shs1-Shs1 or Cdc11-Cdc11), which could prevent a robust interaction with Ext-CC2-CC3 in vivo. These putative regulatory roles of Shs1 are likely absent in the in vitro assays. Regardless of the explanation, our analyses still strongly suggest that Ext-CC2-CC3 likely mediates the targeting and function of Bni5 through its interaction with the terminal subunits of septin complexes/octamers.

### The Ext, CC2, and CC3 regions in Bni5 cooperate to form a structural and functional module at the division site

Because of its vital importance for the localization and function of Bni5 with respect to the septins, we attempted to gain further mechanistic insight into the structure of Ext-CC2-CC3. AlphaFold predicts that CC3 folds back over CC2 to form an antiparallel coiled-coil structure (**Fig. 6 A**). To test this prediction, we conducted an “in vivo reconstitution” experiment by co-expressing GFP-tagged fragments of Bni5 and mScarlet-CC3 from two different plasmids in the same strain under induced condition (**Fig. 6 B**). If two different fragments can interact in vivo, they should exhibit a colocalization signal of GFP and mScarlet at the bud neck. Remarkably, we detected a clear signal of mScarlet-CC3 at the bud neck only when it was co-expressed with fragments containing CC2 but lacking CC3 (**Fig. 6, B and C**). Notably, Ext-CC2, but not CC2 alone, strongly attracted mScarlet-CC3 to the division site, indicating that the Ext region plays an important role in promoting an in-trans interaction between CC2 and CC3. We also found that any GFP-tagged Bni5 fragment containing both CC2 and CC3 prevented the localization of mScarlet-CC3 to the division site (**Fig. 6, B and C**). This is likely due to an intramolecular interaction of CC2 and CC3, which prevents the access of mScarlet-CC3 to CC2 on the GFP-tagged fragment. These results strongly support the AlphaFold prediction that CC2 and CC3 interact to form an anti-parallel coiled-coil structure.

**Figure 6.**
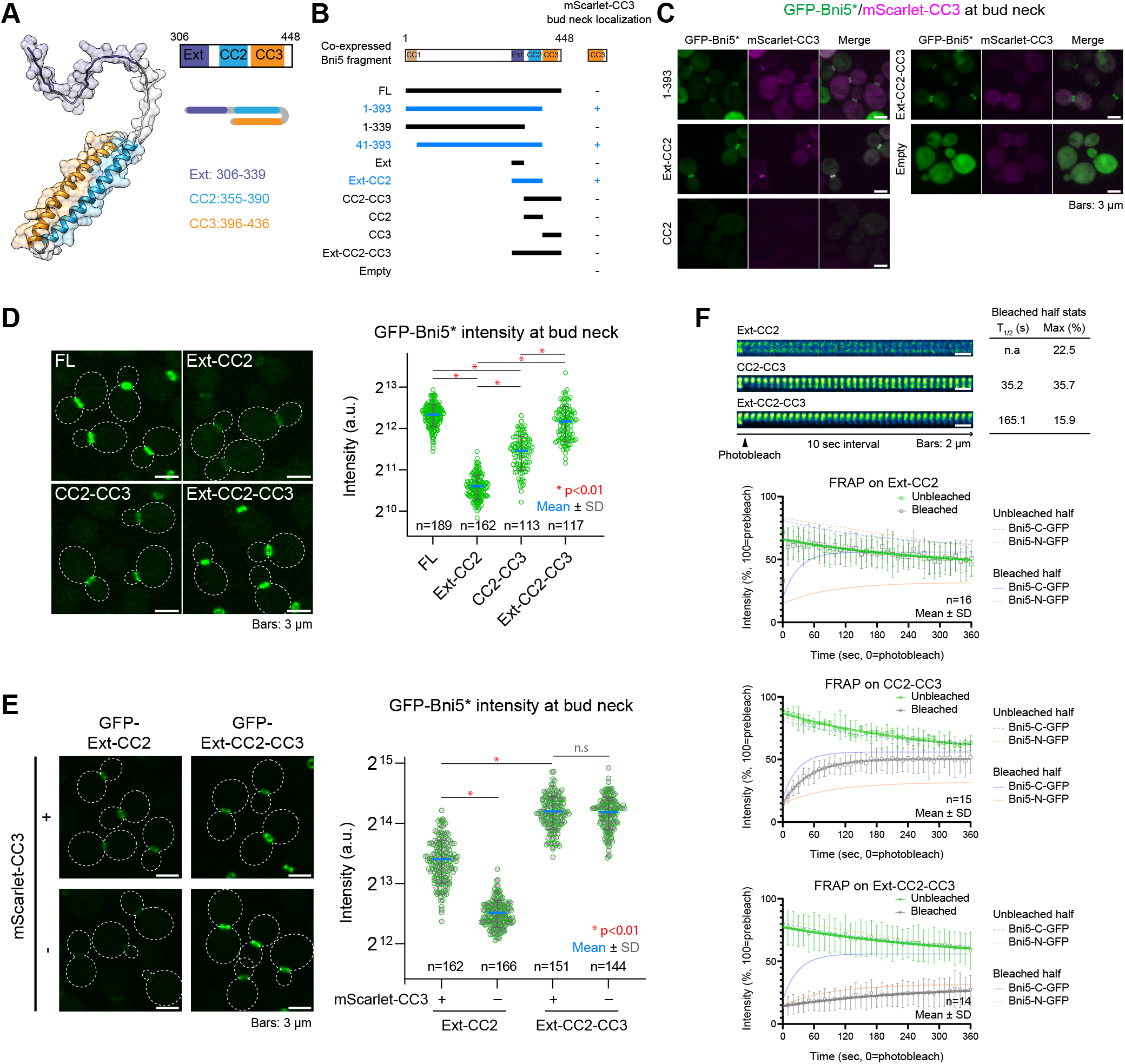
The Ext, CC2, and CC3 domains cooperate to form a structural unit for the stable localization of Bni5 at the division site. (A) AlphaFold model of the Ext-CC2-CC3 module of Bni5. Color codes used are as follows: Ext (306-339) in purple, CC2 (355-390) in blue, and CC3 (306-436) in orange. (B) Summary of the ability of different Bni5 fragments to recruit CC3 to the bud neck. Expression of both GFP-Bni5 fragments and mScarlet-CC3 was induced by culturing in SC-Ura-His-Met medium for five hours. Alleles in blue indicate colocalization with mScarlet-CC3 at the bud neck, which are shown in (C). Bold lines indicate the region of Bni5 in relevant alleles. Strains used are as follows: YEF11834 (*bni5Δ* [*mScarlet-bni5(394-448)*] [*GFP-BNI5(FL)*]), YEF11835 (*bni5Δ* [*mScarlet-bni5(394-448)*] [*GFP-bni5(1-393)*]), YEF11836 (*bni5Δ* [*mScarlet-bni5(394-448)*] [*GFP-bni5(1-339)]*), YEF11837 (*bni5Δ* [*mScarlet-bni5(394-448)*] [*GFP-bni5(41-393)*]), YEF11838 (*bni5Δ* [*mScarlet-bni5(394-448)*] [*GFP-bni5(306-339)*]), YEF11839 (*bni5Δ* [*mScarlet-bni5(394-448)*] [*GFP-bni5(306-393)*]), YEF11840 (*bni5Δ* [*mScarlet-bni5(394-448)*] [*GFP-bni5(340-448)*]), YEF11841 (*bni5Δ* [*mScarlet-bni5(394-448)*] [*GFP-bni5(340-393)*]), YEF11842 (*bni5Δ* [*mScarlet-bni5(394-448)*] [*GFP-bni5(394-448)]*), YEF11843 (*bni5Δ* [*mScarlet-bni5(394-448)*] [*GFP-bni5(306-448)*]), and YEF11844 (*bni5Δ* [*mScarlet-bni5(394-448)*] [*GFP*]). (C) Colocalization of the GFP-tagged Bni5 fragments with mScarlet-CC3 at the bud neck. Asterisks represent variants of Bni5. Strains used are as follows: YEF11835, YEF11839, YEF11841, YEF11843, and YEF11844. (D) Quantitative analysis of different Bni5 fragments at the bud neck. Left: Cells in the exponential growth phase were subjected to imaging. Right: the intensities of the indicated GFP-Bni5 fragments at the bud neck in cells with a small or medium bud (S/G2 cells) were scored. Green circles, blue bars, and gray error bars represent all data points, mean values, and SD values, respectively. An asterisk represents variants of Bni5. Strains used are as follows: YEF11821 [*GFP-BNI5(FL)*], YEF11823 [*GFP-bni5(306-393)*], YEF11824 [*GFP-bni5(340-448)*], and YEF11825 [*GFP-bni5(306-448)*]. (E) Impact of the mScarlet-CC3 induction on the intensity of Bni5 fragments at the bud neck. Left: Cells in the exponential growth phase were cultures for five hours either in SC-His-Met medium or SC-His to or not to induce the expression of mScarlet-CC3, and then subjected to imaging. Right: Intensities of the indicated GFP-Bni5 fragments at the bud neck in cells with a small or medium bud (S/G2 cells) were scored. Green circles, blue bars, and gray error bars represent all data points, mean values, and SD values, respectively. An asterisk represents variants of Bni5. Strains used are as follows: YEF11826 (*GFP-bni5(306-393)* [*mScarlet-bni5(394-448)*]), YEF11827 (*GFP-bni5(306-393)* [*mScarlet*]), YEF11828 (*GFP-bni5(306-448)* [*mScarlet-bni5(394-448)*]), and YEF11829 (*GFP-bni5(306-448)* [*mScarlet*]). (F) FRAP analysis on Bni5 fragments. Half of the ring from each cell was photo-bleached, and fluorescence recovery in the bleached region (gray symbol) and unbleached region (green symbol) was followed over time. Top: Montages of cells were created from time-lapse series taken with a 10-sec interval. Bottom: Lines, symbols, and error bars represent regression curves, mean values, and SD values, respectively. The reference plots of Bni5-C-GFP and Bni5-N-GFP were modified from Figure 1G. Strains used are as follows: YEF11549 (*GFP-bni5(306-339) mScarlet-TUB1*), YEF11551 (*GFP-bni5(340-448) mScarlet-TUB1*), and YEF11552 (*GFP-bni5(306-448) mScarlet-TUB1*).

To investigate the impact of compromising the structural integrity for the Ext-CC2-CC3 module, we measured the targeting efficiency for various constructs (**Fig. 6 D**). The full module exhibited a strong signal intensity comparable to the FL protein (**Fig. 6 D**). Strikingly, when either the Ext or CC3 region was deleted, the signal reduced by 45% or 70% (vs. FL), respectively, (**Fig. 6 D**), indicating that these regions are differentially required for efficient localization at the bud neck. The significant increase in the neck signal of GFP-Ext-CC2 upon the co-expression of mScarlet-CC3 further demonstrated this point (**Fig. 6 E**). Additionally, our FRAP analysis showed that the partial modules, namely, Ext-CC2 and CC2-CC3, were significantly more dynamic compared to the full module of Ext-CC2-CC3 (**Fig. 6 F**), which exhibited a stable signal comparable to FL (Bni5-N-GFP).

Collectively, these data provide a strong support for the AlphaFold prediction that CC2 and CC3 are packed into an antiparallel coiled-coil, which, together with Ext, form a structural and functional module that largely accounts for the stable localization of Bni5 at the division site.

### The disordered MID region regulates the timely removal of Bni5 to facilitate the onset of septin hourglass-to-double ring transition

The long-disordered region between CC1 and Ext-CC2, which we termed MID (<u>mi</u>ddle <u>d</u>isordered region, aa41-305), accounts for ∼60% of the protein, yet its deletion did not cause any obvious defect in Bni5 localization or its function as a septin-myosin-II linker (**Fig. 3, A and B**, see construct of aa1-40 306-448). The signal intensity and turnover rate of GFP-ΔMID at the bud neck was comparable to that of

Bni5-N-GFP (i.e., GFP-FL) (**Fig. 7, A and B**). However, time-lapse analysis revealed a delay in the signal drop of GFP-ΔMID at the bud neck before cytokinesis (**Fig. 7, C and D**). The GFP-FL signal started to drop rapidly about 12 minutes before the spindle break (-12 min) and disappeared completely at the time of the spindle break (0 min), whereas GFP-ΔMID began to drop sharply about -6 min and reached the bottom three minutes after the spindle break (3 min). Because the kinetic behavior of GFP-ΔMID is similar to that of Myo1 (Okada et al., 2021b), we thought that the CC1-mediated interaction with Myo1 might be involved. However, deleting CC1 from GFP-ΔMID (i.e., GFP-ΔCC1ΔMID) did not affect its disappearing kinetics at the bud neck (**Fig. 7, C and D**). Thus, the MID region appears to control the timely removal of Bni5 from the division site independently of Myo1.

**Figure 7.**
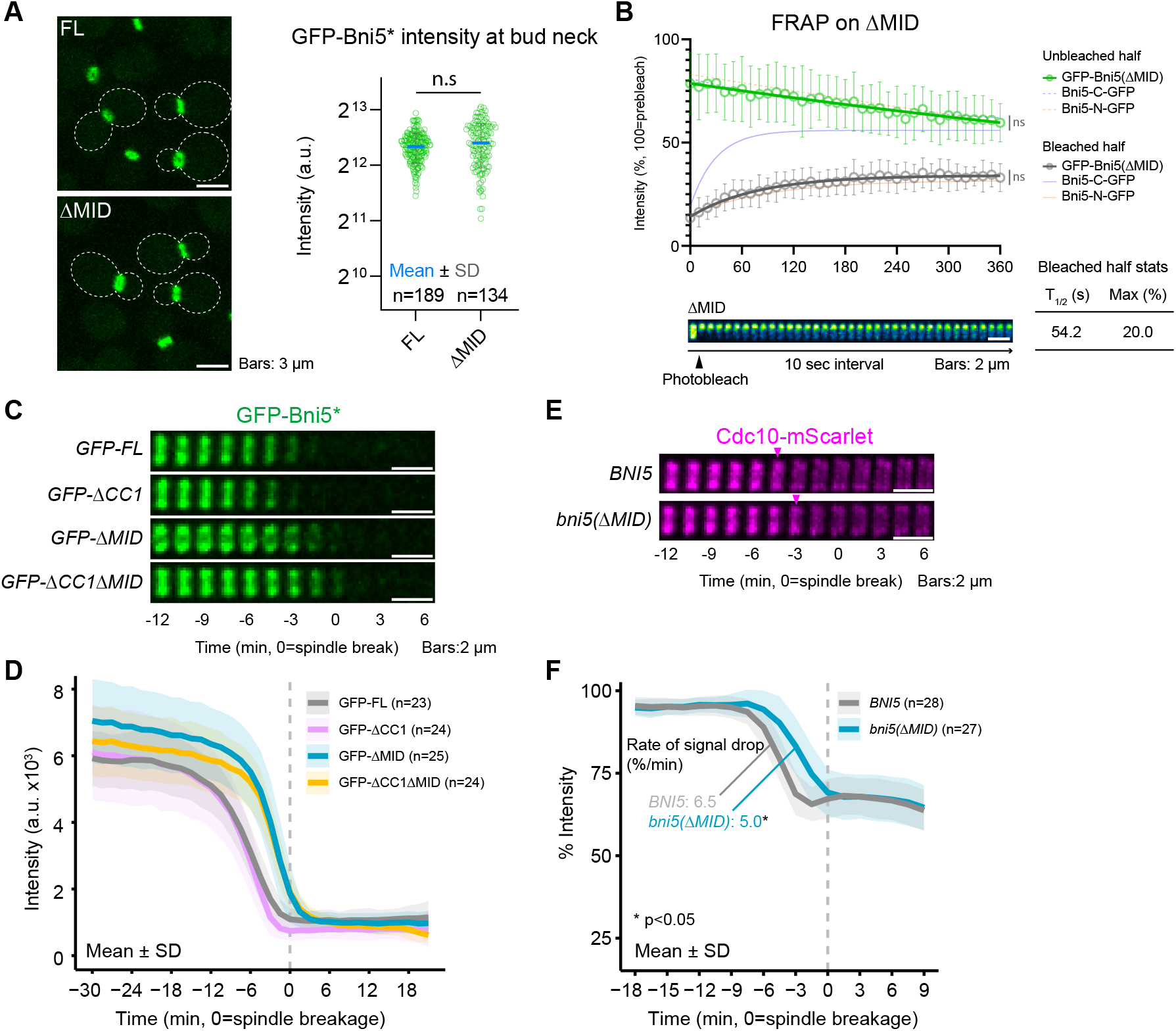
The disordered MID region regulates the timely removal of Bni5 to facilitate the onset of septin hourglass-to-double ring transition. (A) The MID is dispensable for the bud neck localization of Bni5. Left: Cells in the exponential growth phase were subjected to imaging. Right: GFP-Bni5(ΔMID) intensity at the bud neck was scored from cells with a small or medium bud (S/G2 cells). Green circles, blue bars, and gray error bars represent all data points, mean values, and SD values, respectively. An asterisk represents variants of Bni5. Strains used are as follows: YEF11546 [*mScarlet-TUB1 GFP-BNI5(FL)*] and YEF11553 [*mScarlet-TUB1 GFP-bni5(Δ41-305)*]. (B) FRAP analysis of Bni5(ΔMID) at the bud neck. Half of the ring from each cell was photo-bleached, and fluorescence recovery in the bleached region (gray symbol) and unbleached region (green symbol) was followed over time. Top: Lines, symbols, and error bars represent regression curves, mean values, and SD values, respectively. The reference plots of Bni5-C-GFP and Bni5-N-GFP were modified from Figure 1G. Bottom: Montages of cells were created from time-lapse series taken with a 10-sec interval. Strain used is YEF11553. (C) Time-lapse analysis of the impact of MID1 deletion on Bni5 localization during the cell cycle. Montages of the indicated Bni5 fragments at the bud neck were created from selected frames of time-lapse series taken with a 1.5-min interval. An asterisk represents variants of Bni5. Strains used are as follows: YEF11546 [*mScarlet-TUB1 GFP-BNI5(FL)*], YEF11600 [*mScarlet-TUB1 GFP-bni5(41-448)*], YEF11553 [*mScarlet-TUB1 GFP-bni5(Δ41-305)*], and YEF11552 [*mScarlet-TUB1 GFP-bni5(306-448)*]. (D) Kinetics of the bud neck accumulation for the GFP-tagged Bni5 proteins imaged in (C). Bold lines and associated shaded bands represent mean and SD values, respectively. (E) Time-lapse analysis of the septin hourglass-to-double ring transition in WT and *bni5(ΔMID)* cells. Montages of Cdc10-mScarlet at the bud neck were created from selected frames of time-lapse series taken with a 1.5-min interval. Strains used are as follows: YEF11772 [*CDC10-mScarlet Venus-TUB1 GFP-BNI5(FL)*] and YEF11773 [*CDC10-mScarlet Venus-TUB1 GFP-bni5(Δ41-305)*] (F) Kinetics of the neck-localized Cdc10-mScarlet quantified from the imaging data in (E). Bold lines and associated shaded bands represent mean and SD values, respectively.

Because the FL Bni5 disappeared during the septin hourglass-to-double ring (HDR) transition at the onset of cytokinesis (**Fig. 7 D and Fig. 7 E**, arrowhead), we examined the possible effect of the delayed removal of GFP-ΔMID on this septin remodeling event. The HDR transition is controlled by the mitotic exit network (MEN, an equivalent pathway of Hippo and SIN in human and fission yeast, respectively), which allows the PM access and constriction of the AMR sandwiched by the septin double ring (Lippincott et al., 2001; Tamborrini et al., 2018). The HDR transition is accompanied by a sharp drop in septin intensity in the WT strain (-6 to -3 min, *BNI5,* **Fig. 7 F**). Strikingly, the onset of the transition was delayed (**Fig. 7 E**, arrowhead) and the rate of signal drop was decreased (**Fig. 7 F**) in *ΔMID* strain than in WT. These data suggest that the MID region controls the timely removal of Bni5, which, in turn, controls the timely execution of the septin HDR transition to allow AMR constriction.

### Bni5 tethers myosin-II to the septin hourglass at the division site before cytokinesis to facilitate the retrograde actin cable flow

The function of myosin-II (Myo1), as part of the AMR, in cytokinesis has been extensively studied, but its role at the division site prior to actin ring assembly in anaphase has remained poorly understood. One such role for Myo1 is thought to facilitate the rate of RACF via its motor activity (**Fig. 8 A**) (Huckaba et al., 2006), which is important for asymmetric segregation of mitochondrial fitness to control cellular lifespan (Higuchi et al., 2013). Since Bni5 links Myo1 to the septins, it likely plays a similar role as Myo1 in controlling the rate of RACF.

**Figure 8.**
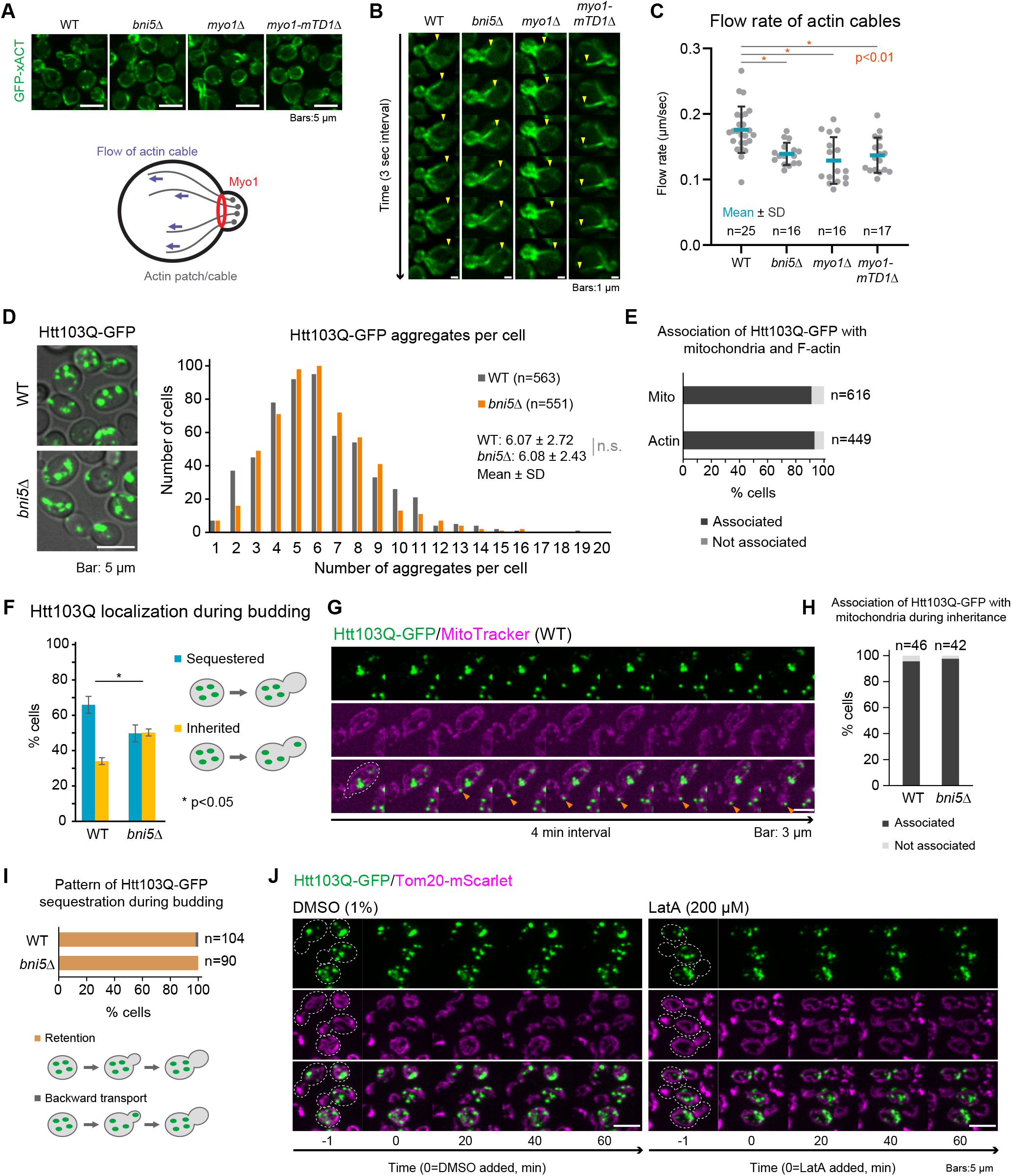
Bni5-mediated localization of Myo1 at the division site before cytokinesis facilitates retrograde actin cable flow and asymmetric segregation of protein aggregates. (A) Images of actin cables marked by the GFP-xACT probe in different strains. Cells were cultured in SC medium at 25°C for 18 hours before imaging. Prior to culturing, the *myo1Δ* strain was grown on a 5-FOA plate to cure the *MYO1* cover plasmid. Strains used are as follows: YEF10201 (*GFP-xACT = ECM25[aa536-588]*), YEF10173 (*bni5Δ GFP-xACT*), YEF11489 (*myo1-mTD1Δ GFP-xACT*), YEF10170 (*myo1Δ GFP-xACT*). (B) Images of RACF in *bni5* and *myo1* mutants. Montages of actin cables undergoing RACF were created from selected frames of a time-lapse series taken with a 1-sec interval. Yellow arrowheads represent the ends of extending actin cables during RACF. Strains used are the same as in (A). (C) Rate of RACF in *bni5* and *myo1* mutants. The rates of RACF in the indicated strains were quantified from the imaging data in (B). (D) Formation of Htt103Q-GFP aggregates in WT and *bni5Δ* cells. The *HTT103Q-GFP* gene carried on a plasmid was overexpressed in the indicated cells by four-hour culturing in SC-Ura+2% galactose medium. Strains used are as follows: YEF11921 ([*proGAL1-HTT103Q-GFP*]) and YEF11920 (*bni5Δ* [*proGAL1-HTT103Q-GFP*]). (E) Association of Htt103Q-GFP with mitochondria and F-actin in WT. The association of Htt103Q-GFP with mitochondria (marked by Tom20-mScarlet) in live cells and F-actin (marked by phalloidin) in fixed cells was separately scored by examining each z-series of confocal images. Strains used are as follows: YEF11949 (*TOM20-mScarlet* [*proGAL1-HTT103Q-GFP*]) and YEF11921. (F) Asymmetric segregation of Htt103Q-GFP during budding in WT and *bni5Δ* cells. Cells induced to form Htt103Q-GFP aggregates were subjected to live-cell imaging. Data from > 150 cells for each strain, acquired from three independent experiments, were used for quantification. Strains used are the same as in (D). (G) Inheritance of mitochondria-associated Htt103Q-GFP aggregates by bud. Cells induced to form Htt103Q-GFP aggregates were subjected to live-cell imaging. Mitochondria were visualized using MitoTracker. Arrowheads indicate Htt103Q-GFP aggregates during inheritance. Strains used are the same as in (D). (H) Association of Htt103Q-GFP aggregates with mitochondria during inheritance in the indicated strains were quantified from the imaging data in (G). (I) Patterns of Htt103Q-GFP sequestration during bud growth in the indicated strains were quantified from the imaging data in (F). (J) Impact of F-actin disruption on Htt103Q-GFP aggregates and mitochondria. Cells induced to form Htt103Q-GFP aggregates were treated with either 200 µM LatA or 1% DMSO and then subjected to live-cell imaging. Montages were created from selected frames of a time-lapse series taken with a 2-min interval. Strain used is YEF11949.

To examine this possibility, we used GFP-xACT, a recently developed actin cable probe derived from the budding yeast Ecm25 (Duan et al., 2021), to visualize and track the movement of actin cables. GFP-xACT successfully decorated actin cables in both WT and mutants (**Fig. 8 A**). Time-lapse imaging was performed to trace the extension of actin cables (**Fig. 8 B**). As expected, deletion of *MYO1* decreased the rate of RACF by 27% (**Fig. 8 C**). Similarly, deletion of *BNI5* decreased the rate of RACF by 21%. Remarkably, deletion of the Bni5-binding site in Myo1 (i.e., *myo1*-*mTD1Δ*) caused a nearly identical decrease (22%) as the deletion of *BNI5* did (**Fig. 8, B and C**). These data indicate that Bni5 tethers myosin-II to the septin hourglass before cytokinesis to facilitate the RACF.

Since RACF is involved in the asymmetric segregation of mitochondrial fitness (Higuchi et al., 2013) and mitochondria in budding yeast can serve as an organelle for the spatial sequestration of protein aggregates in mother cells (Zhou et al., 2014), it raises the possibility that the reduction of RACF impacts not only the asymmetric segregation of mitochondria but also the sequestration of protein aggregates. To test this possibility, we overexpressed Htt103Q-GFP, a mutant fragment of huntingtin, a protein that causes Huntington’s disease and leads to the formation of protein aggregation in the yeast system (Meriin et al., 2002; Song et al., 2014). A four-hour induction of galactose-mediated gene overexpression led to an average of ∼6 aggregates per cell in both WT and *bni5Δ* (**Fig. 8 D**). These aggregates are mostly (>90%) associated with mitochondria (Tom20-mScarlet) and F-actin structures (Alexa Fluor-594 Phalloidin), as observed through dual-color imaging (**Fig. 8 E**), confirming previous observations (Song et al., 2014; Zhou et al., 2014).

Remarkably, as revealed by time-lapse imaging, Htt103Q-GFP aggregates were more likely to be inherited by the daughter cells of *bni5Δ* (50.2 ± 2.0%, **Fig. 8 F**) than of the WT (34.1 ± 4.8%, **Fig. 8 F**), supporting our hypothesis. Notably, the majority of aggregates inherited by the bud were associated with mitochondria (**Fig. 8, G and H**), suggesting that the daughter inheritance is not caused by the diffusion of free aggregates from mother to daughter but rather by defects in an active sorting and transport mechanism of mitochondria to daughter cells. Time-lapse analysis of bud formation in both WT and *bni5Δ* cells indicated that aggregates were strictly retained in mother cells (**Fig. 8 I**), suggesting that spatial retention, rather than backward movement of aggregates from daughter to mother, accomplishes this sequestration.

To gain further mechanistic insight, we performed time-lapse imaging of cells treated with latrunculin A (LatA), a toxin that disrupts actin filaments (Ayscough et al., 1997). Remarkably, neither Htt103Q-GFP nor mitochondria, nor their association, were affected in the presence of LatA (**Fig. 8 J**), suggesting that F-actin is not required for the maintenance of Htt103Q tethering to mitochondria. Strikingly, all the aggregates inherited by the bud (6 out of 57 cells) were associated with mitochondria. These results suggest that RACF mediates spatial sequestration of protein aggregates through the selective inheritance of aggregate-free mitochondria by the bud.

Taken together, these results demonstrate that Bni5 links Myo1 to the septins at the bud neck to mediate rapid RACF and the asymmetric segregation of protein aggregates before cytokinesis.

### Bni5-mediated localization of Myo1 at the division site before cytokinesis endows the AMR robustness against internal and external stress during cytokinesis

Since Bni5 is removed from the bud neck before the onset of cytokinesis (**Fig. S3 B**), it is unlikely to impact cytokinesis in a direct way. Deletion of *BNI5* was previously shown to cause a partially penetrant defect in cytokinesis and cell morphology at high temperatures (Lee et al., 2002). However, we were unable to reproduce these findings in four different laboratory strains, including our own, as we observed no obvious growth or morphological defects in the *bni5Δ* strains at either 25°C or 37°C (**Fig. 9, A and B, and Fig. S5, A and B**), as well as no significant difference in Myo1 constriction during cytokinesis (**Fig. 9 C**). The discrepancy may be attributed to the presence of *YFP-CDC10* in the previous strains (Lee et al., 2002), which may produce synthetic phenotypes with *bni5Δ*. Regardless of the reasons for the discrepancy, our data indicate that deletion of *BNI5* alone does not cause any obvious defect in cytokinesis.

**Figure 9.**
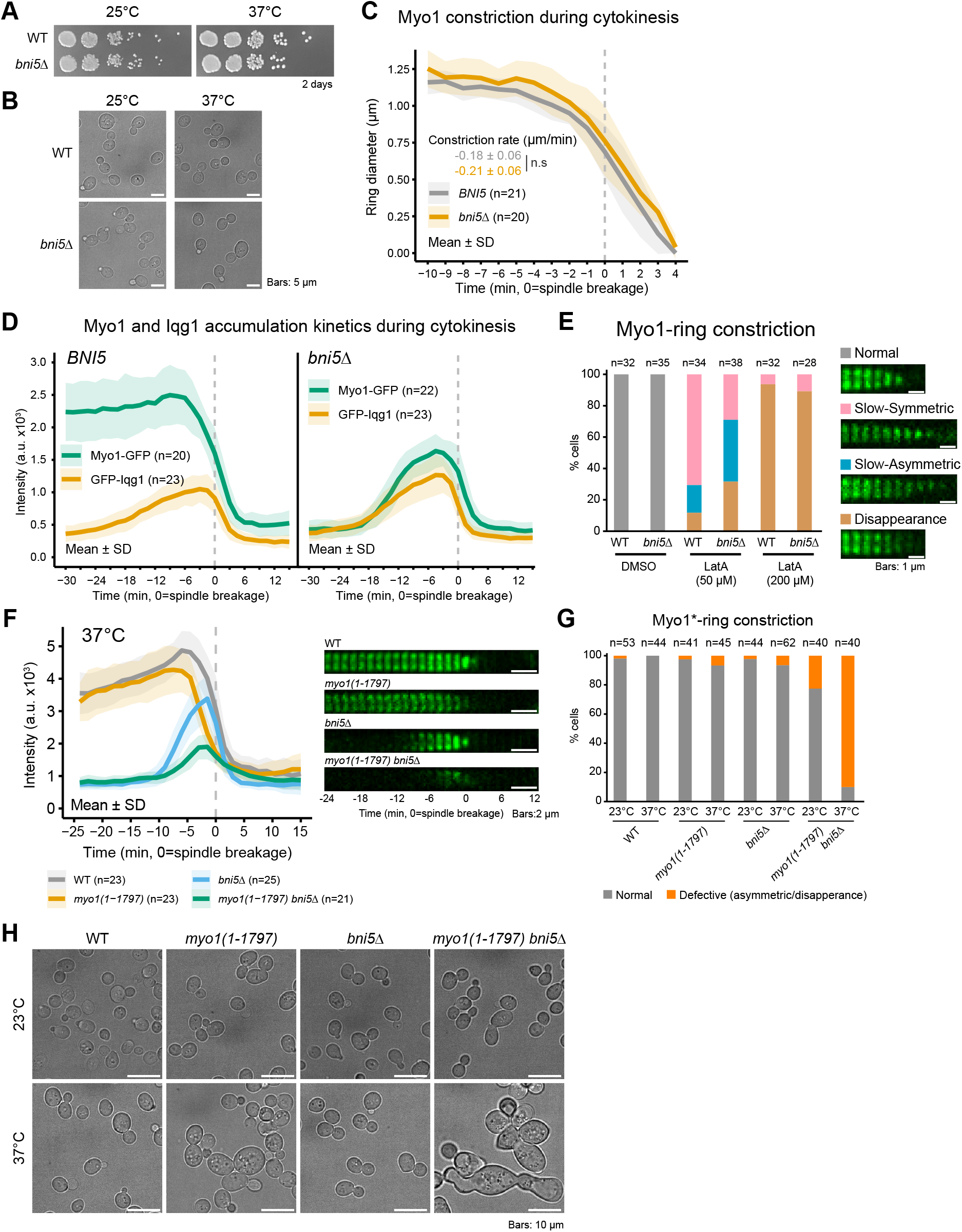
Bni5-mediated localization of Myo1 at the division site before cytokinesis endows the AMR robustness against internal and external stress during cytokinesis. (A) Growth of *bni5Δ* cells on YPD plates. Spot assay result documented after two days of incubation at 25°C or 37°C. Strains used are as follows: YEF473A (WT) and YEF9654 (*bni5Δ*). (B) Cell morphology of *bni5Δ* cells. Cells cultured in YM-1 medium at 25°C or 37°C for 24 hours and subjected to imaging. Strains used are the same as (A). (C) Constriction of Myo1 ring during cytokinesis in WT and *bni5Δ* cells. The temporal changes in diameter of the Myo1 ring during cytokinesis were quantified from time-lapse series taken with a 1-min interval. Bold lines and associated shaded bands represent mean and SD values, respectively. Strains used are as follows: YEF11385 (*MYO1-GFP mScarlet-TUB1*) and YEF11443 (*bni5Δ MYO1-GFP mScarlet-TUB1*). (D) Accumulation kinetics of Myo1 and Iqg1 in *bni5Δ* cells during cytokinesis. Intensities of indicated proteins at the bud neck were measured from time-lapse series taken with a 1.5-min interval. Bold lines and associated shaded bands represent mean and SD values, respectively. Strains used are as follows: YEF11385, YEF11443, YEF11412 (*GFP-IQG1 mScarlet-TUB1*), and YEF11464 (*bni5Δ GFP-IQG1 mScarlet-TUB1*). (E) Impact of F-actin disruption on AMR constriction in *bni5Δ* cells. AMR (Myo1-GFP) constriction in WT and *bni5Δ* cells in the presence of moderate (50 µM) or high (200 µM) concentrations of LatA was monitored. AMR constriction was scored based on the indicated category from the time-lapse series taken with a 2-min interval. Strains used are the same as in (C). (F) Impact of Bni5 deletion, the tail truncation of Myo1, and high temperature on the Myo1 accumulation kinetics at the bud neck. Left: Accumulation kinetics of Myo1 in indicated mutants during cytokinesis. Intensities of indicated proteins at the bud neck were measured from time-lapse series taken with a 1.5-min interval. Bold lines and associated shaded bands represent mean and SD values, respectively. Right: Montages of indicated proteins at the bud neck were created from selected frames. Strains used are as follows: YEF11529 (*GFP-MYO1 mScarlet-TUB1*), YEF11524 [*GFP-myo1(1-1797) mScarlet-TUB1)*], YEF11601 (*bni5Δ GFP-MYO1 mScarlet-TUB1*), and YEF11530 [*bni5Δ GFP-myo1(1-1797) mScarlet-TUB1)*]. See also **Figure S5**. (G) Quantitative analysis of the AMR constriction phenotypes in the indicated strains was scored based on the indicated categories using the imaging data in (F). See also **Figure S5**. (H) Cell morphology of the *myo1* tail-truncation allele and *bni5Δ* cells. Cells of indicated strains were cultured in YM-1 medium at 25°C or 37°C for 24 hours and subjected to imaging. Strains used are the same as in (F).

During our analysis of cytokinesis in *bni5Δ* cells, we unexpectedly found that the peak intensity of Myo1 at the division site was reduced by 35% (2496.5 ± 466.0 vs. 1635.1 ± 262.2, p<0.01, **Fig. 9 D**). However, the peak intensity of Iqg1, the sole mediator of Myo1 localization during cytokinesis, remained unchanged in *bni5Δ* cells, suggesting that the reduction of Myo1 intensity is not due to the reduction of Iqg1 at the division site (**Fig. 9 D**). Importantly, Myo1 lacking the Bni5-binding site (Myo1-mTD1Δ) showed a very similar accumulation profile at the division site as the full-length Myo1 did in *bni5Δ* cells (**Fig. S5, C and D**, r=0.98), suggesting that the binding of Bni5 to the mTD1 of Myo1 mediates its role in the accumulation of Myo1 at the division site before cytokinesis. Since Myo1 plays a scaffolding role in the assembly of a multi-component cytokinetic machine (Wloka et al., 2013), we also monitored the accumulation of other cytokinetic proteins in *bni5Δ* cells to determine whether they would decrease following the reduction of Myo1 at the division site. Surprisingly, the intensities of two major cytokinetic proteins, Iqg1 and Chs2, were not affected in *bni5Δ* cells (**Fig. 9 D and Fig. S5 E**), which explains why cytokinesis appeared to be normal when the *bni5Δ* cells were grown under standard conditions (**Fig. 9, A-C**).

Does the Bni5-mediated increase of Myo1 at the division site play a role in cytokinesis? We hypothesized that this increase of myosin-II in the AMR benefits the integrity and robustness of the ring structure against genetic and/or environmental insults. To test this hypothesis, we treated cells with LatA. A high dose of LatA (200 µM) abolished actin filaments in both WT and *bni5Δ* cells, causing Myo1 disappearance without constriction during cytokinesis (**Fig. 9 E**) (Bi et al., 1998; Okada et al., 2021b). A lower dosage (50 µM) caused a mild phenotype in most of WT cells (slow symmetric constriction, 71%, 24/34 cells, **Fig. 9E**). The rest of the cells showed a severe phenotype, i.e., slow asymmetric constriction (18%, 6/34 cells, **Fig. 9E**) or disappearance without constriction (12%, 4/34 cells, **Fig. 9E**). In comparison, the same dosage induced a more severe phenotype in *bni5Δ* cells (slow asymmetric, 39%, 15/38 cells, and disappearance, 32%, 12/38 cells, **Fig. 9E**). These results suggest that the AMR with reduced Myo1 amount is more susceptible to F-actin perturbation.

We also tested the role of Bni5 in cytokinesis when Myo1 was compromised. We have previously shown that Myo1 lacking a small tail region (aa1798-1928) became more mobile at the division site during cytokinesis (Wloka et al., 2013). Kinetic analysis of cells imaged at 23°C revealed that single mutants of *myo1(1-1797)* and *bni5Δ* lost their peak values of Myo1 by 5% and 37%, respectively, compared to the WT, whereas the double mutant of *myo1(1-1797) bni5Δ* lost by 54% (**Fig. S5 F**). A similar but more profound effect was observed in cells exposed to a temperature stress at 37°C. The single mutants, *myo1(1-1797)* and *bni5Δ*, and double mutant, *myo1(1-1797) bni5Δ*, lost their peak values of Myo1 by 13%, 31%, and 61%, respectively (**Fig. 9 F**). We also scored the Myo1 constriction phonotype: at 23°C, 23% (9/40 cells) of the double mutant cells showed the severe phenotype (asymmetric constriction or disappearance of the ring, **Fig. 9G**), whereas only 2% of both single mutant cells did so. Strikingly, at 37°C, 90% of the double mutant cells (36/40 cells) exhibited the severe phenotype (**Fig. 9 G**), accompanied by pronounced defects in cytokinesis (**Fig. 9 H**). These results suggest that the AMR can tolerate some amount of Myo1 structural defect but it becomes extremely vulnerable to a temperature stress when the Bni5-mediated increase of Myo1 at the division site is eliminated.

Taken together, the Bni5-mediated increase of Myo1 at the division site before cytokinesis endows the AMR, and therefore cytokinesis, the robustness against various perturbations.

## DISCUSSION

Bni5 has been shown to be the sole linker between the septin hourglass and the myosin-II heavy chain Myo1 at the division site before cytokinesis (Fang et al., 2010), but the molecular interactions and cellular functions underlying this linker role have remained largely unknown. In this study, guided by the AlphaFold-predicted structure of Bni5, we have performed a comprehensive structure-function analysis, and found that Bni5 binds to Myo1 via its N-terminal CC1 and binds to septins and the Elm1 kinase (a septin regulator) via its C-terminal Ext-CC2-CC3 (**Fig. 10**). Importantly, we have defined three functions for Bni5 (**Fig. 10**): 1) from G1 phase to anaphase, Bni5 tethers Myo1 to the septin hourglass at the division site to facilitate the retrograde flow of actin cables, which, in turn, regulates asymmetric segregation of toxic protein aggregates, 2) this tethering function leads to a local increase of Myo1 at the division site before cytokinesis, which endows the AMR robustness against various stresses during cytokinesis, and 3) the timely removal of Bni5 from the division site is regulated, at least in part, by its MID region, which contributes to the timely remodeling of the septin hourglass into a double ring that enables AMR constriction.

**Figure 10.**
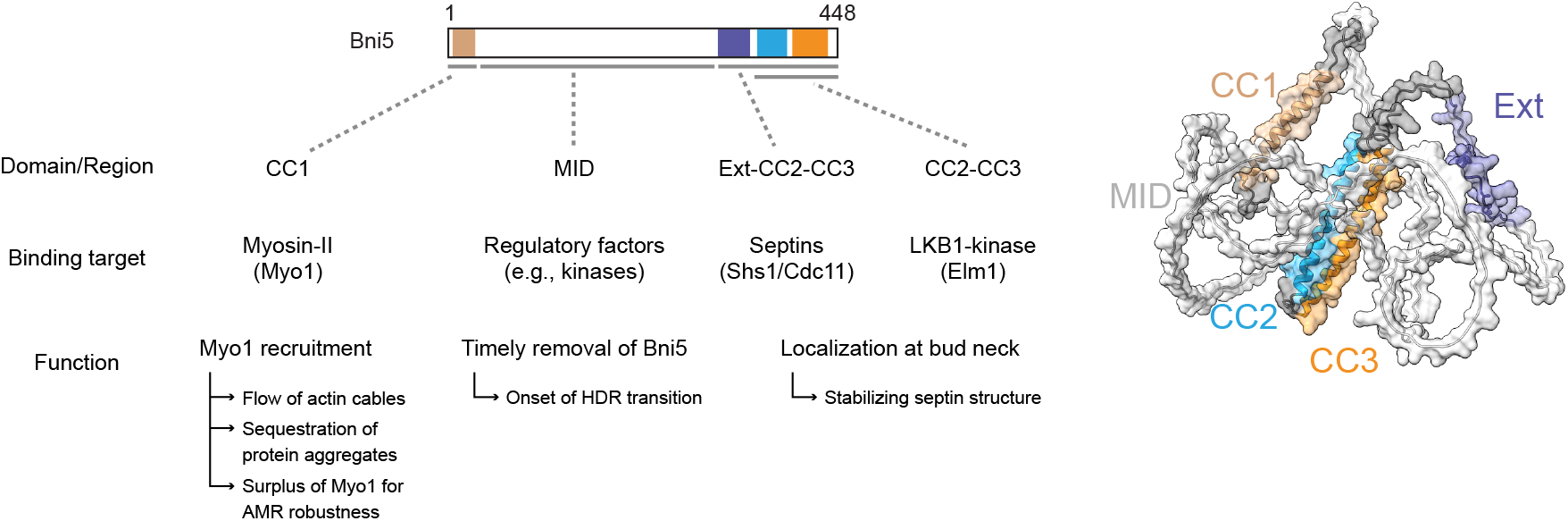
Distinct domains of Bni5 and their binding targets and specific functions. See details in the text.

### The Ext-CC2-CC3 region of Bni5 defines a structural and functional unit that stabilizes the septin hourglass and controls the timely remodeling of the hourglass into a double ring

We found that the CC2 region of Bni5 is necessary but not sufficient for its targeting to the bud neck. To fulfill sufficiency, an upstream (Ext) or a downstream (CC3) region is required. While both Ext-CC2 and CC2-CC3 are capable of targeting to the bud neck on their own, they differ in the timing, magnitude, and turnover kinetics of their localization. Ext-CC2 localizes to the polarization site before bud emergence, resembling the septins, whereas CC2-CC3 localizes to the bud neck during or shortly after bud emergence, resembling Elm1. The bud-neck signal of CC2-CC3 is stronger than that of Ext-CC2 but only Ext-CC2-CC3 reaches similar levels of localization as the FL Bni5. Additionally, both Ext-CC2 and CC2-CC3 display rapid turnover at the division site and cannot suppress the *cdc12-*6 septin mutant, but Ext-CC2-CC3 becomes stabilized and is able to suppress the septin mutant defects, akin to the FL Bni5. Thus, Ext-CC2-CC3 defines the structural and functional unit in Bni5 that is largely responsible for its localization at the bud neck as well as its role in septin regulation.

Overexpressed Ext alone can target to the bud neck weakly and this targeting depends on the terminal subunits (Cdc11 and Shs1) of the septin complexes but not Elm1, suggesting that Ext localize to the bud neck by interacting with the septins with low affinity. This localization is enhanced by adding CC2, which might be due to a structural stabilization, leading to the formation of a conformational state of Ext-CC2 that enables efficient interaction with the septins. This possibility is supported by the observation that Ext-CC2 interacts strongly with the septin filaments in vitro. Another non-mutually exclusive possibility is that the addition of CC2 to Ext provides a novel binding site for a non-septin protein at the bud neck. This is supported by the observation that Ext-CC2 also interacts with Elm1 in vitro.

AlphaFold predicts that CC2 and CC3 form an anti-parallel coiled-coil structure. The weak cytoplasmic signals of GFP-CC2 and mScarlet-CC3 suggest that CC2 and CC3 on their own are unstable. The predicted anti-parallel interaction between CC2 and CC3 is supported by the observation that the cytosolic mScarlet-CC3 is recruited to the neck localized GFP-Ext-CC2 (CC2 is available for interaction with mScarlet-CC3), but not by GFP-CC2-CC3 or GFP-Ext-CC2-CC3 (CC2 is not available for interaction with mScarlet-CC3). The enhanced signal of GFP-Ext-CC2 by mScarlet-CC3 at the bud neck, together with the observation that GFP-CC2-CC3 localizes to the bud neck in an Elm1-dependent manner, suggests that the interaction between CC2 and CC3 stabilizes each other, leading to a CC2-CC3 folding that is competent for Elm1 binding. The fact that both Ext-CC2 and CC2-CC3 are able to bind to Elm1 in vitro indicates a key role of CC2 in these interactions.

Collectively, our study suggests that Ext-CC2-CC3 is organized into a structure as predicted by AlphaFold2 (**Fig. 10**) and this structure most likely binds to septin filaments and Elm1 via Ext and CC2, respectively, with CC3 involved in structural stabilization by forming an anti-parallel dimer with CC2. This anti-parallel interaction would position the GFP moiety of Bni5-C-GFP in close proximity to the Ext region, potentially disrupting the interaction between the Ext and the septins, causing the FL Bni5-C-GFP to behave essentially like CC2-CC3. Indeed, the timing for the recruitment of both Bni5-C-GFP and CC2-CC3 to the bud neck and their turnover kinetics at the division site resemble those of Elm1 and their localization at the bud neck completely depends on Elm1. Thus, this study provides the most reasonable explanation for the specific defect of Bni5-C-GFP in septin binding and regulation.

Bni5 dissociates from the bud neck before the onset of cytokinesis (Lee et al., 2002). Our analysis indicates that Ext-CC2-CC3 retains this property. However, its removal from the bud neck is delayed in comparison to FL Bni5, and this delay appears to be due to the deletion of the disordered MID region. This interruption in Bni5 removal leads to a delay in the initiation of and a decrease in rate of the hourglass-to-double ring (HDR) transition, suggesting that Ext-CC2-CC3 stabilizes the septin hourglass. Strikingly, among the 34 phosphorylation sites identified in Bni5 (a total of 448 residues) (data available at https://www.yeastgenome.org/), ∼65% of the sites are either within or flanking the Ext region (with one in CC1 and the remainder in the MID region), suggesting that the septin binding capacity of Bni5 is highly regulated. Indeed, mutagenizing four flanking sites (Ser270, Thr274, Ser346, and Ser350) into non-phosphorylatable residues (4Ala) has been reported to cause Bni5 dissociate from the bud neck at earlier time points and attenuate its interaction with the septins (Nam et al., 2007a; Nam et al., 2007b), although another report questioned the impact of these phosphorylations on its septin-specific functions (Finnigan et al., 2015a). As Bni5 interacts with two septin hourglass-associated kinases, Gin4 and Elm1 (Marquardt et al., 2020; Mortensen et al., 2002) and is a direct substrate of Elm1 (Patasi et al., 2015), and the septin-binding Ext region is proximal to the Elm1-binding site within the CC2-CC3 region, it would be important to determine the impact of Gin4 and Elm1-mediated phosphorylation of Bni5 on its interaction with the septins.

### The CC1 region of Bni5 mediates its interaction with myosin-II at the division site to facilitate the retrograde actin flow for asymmetric inheritance and increase local concentration for the robustness of cytokinesis

Previously, we showed that Bni5 is the sole linker between the septin hourglass and myosin-II at the division site before the onset of anaphase, and that Bni5 interacts with the mTD1 (minimal Targeting Domain 1) in the N-terminal portion of Myo1 tail (Fang et al., 2010). Subsequently, the Myo1-interacting region in Bni5 was narrowed to a large fragment (aa1-341) of Bni5 (Schneider et al., 2013). In this study, we demonstrate that the CC1 region (aa1-40) of Bni5 is both necessary and sufficient for binding to the mTD1 of Myo1 in vivo and in vitro. As mTD1 is in the coiled-coil (CC) region of the Myo1 tail, the CC1 interacts with mTD1 most likely through a CC interface. Our FRAP analysis indicates that Myo1 at the division site is far more mobile than the septin hourglass and Bni5 before anaphase, suggesting that the binding affinity between the CC1 of Bni5 and the mTD1 of Myo1 is weaker than that between Bni5 and the septin filaments. The remarkably weak signal of GFP-CC1 at the bud neck during cytokinesis agrees with this possibility. A weak interaction between the stable anchor Bni5 and the dynamic Myo1 is ideally suited for the two cellular functions of Myo1 – RACF and AMR assembly – before the onset of cytokinesis as described below.

Myo1 is known to facilitate the RACF from the bud tip towards the mother cell, and this plays a critical role in the asymmetric inheritance of mitochondrial fitness, with the “young” and “less oxidized” mitochondria preferentially segregated to the daughter cell (Higuchi et al., 2013; Huckaba et al., 2006). Since Bni5 mediates Myo1 localization at the division site before the actin ring assembly in anaphase (Fang et al., 2010), it likely plays a role in RACF through Myo1, independent of the actin ring. Indeed, we found that deletion of *BNI5* slowed down RACF to the same degree as deleting the mTD1-coding sequence in *MYO1*, suggesting that Bni5 facilitates RACF by binding to mTD1 of Myo1. The weak interaction between the stable Bni5 and the dynamic Myo1 at the bud neck may provide spatial flexibility for the interactions of the flowing cables with the septins at the bud neck.

Protein aggregates (marked with Hsp104-GFP) are captured by mitochondria and preferentially retained in the mother cell (Zhou et al., 2014). Protein aggregates (Htt103Q-GFP and Hsp104-GFP) are also shown to be associated with actin cables (Liu et al., 2010; Song et al., 2014). The fact that RACF is involved in the asymmetric inheritance of mitochondrial fitness raises the possibility that RACF might contribute to the asymmetric retention of protein aggregates. Indeed, we found that the Htt103Q-GFP aggregates are tethered to mitochondria and transported to the daughter cell more frequently in *bni5Δ* cells than in WT, and this transport depends on actin filaments. Thus, Bni5 likely contributes to the biased retention of the mitochondria-tethered protein aggregates in the mother cell by promoting RACF.

Bni5 and Iqg1 mediate Myo1 localization at the bud neck before and during cytokinesis, respectively (Fang et al., 2010). These two targeting mechanisms overlap from the onset of anaphase to the onset of mitotic exit (or onset of cytokinesis), with Bni5 and Iqg1 at the bud neck being progressively decreased and increased, respectively (Fang et al., 2010; Lee et al., 2002; Okada et al., 2021b; Schneider et al., 2013). It is during this period, Myo1 is released from Bni5 and passed to Iqg1. Meanwhile, Bni5 is degraded while Myo1 becomes progressively immobilized, suggestive of filament assembly (Lee et al., 2002; Ong et al., 2014; Wloka et al., 2013). Thus, the weak interaction between the stable Bni5 and the dynamic Myo1 at the bud neck could enable an efficient transfer of Myo1 from Bni5 to Iqg1 for filament assembly. While Bni5 is completely dissociated from the bud neck and degraded by the onset of cytokinesis, its presence leads to an ∼35-40% of increase of Myo1 at the bud neck. We have demonstrated that this Bni5-mediated Myo1 increase at the division site endows the AMR with robustness needed to endure different insults, including LatA treatment and a small deletion in the C-terminus of Myo1. In fission yeast, increasing the cellular level of myosin-II (Myo2) also promotes the AMR assembly (Stark et al., 2010). In mice, the non-muscle myosin-IIB (NM-IIB) becomes essential for blood vessel formation when one copy of the gene encoding NM-IIA is ablated in endothelial cells (Ma et al., 2020), indicating that the cellular level of NM-IIA is important for sprouting angiogenesis. Thus, the cellular level of myosin-II is carefully regulated to drive distinct cellular processes including cytokinesis.

### Retrograde flow systems in mammalian cells could be informed by the septin-Bni5-myosin-II mechanism in asymmetrically dividing yeast cells

Retrograde flow is known to drive many biological processes in mammalian cells, including the inward movement of T cell receptor (TCR) microclusters at the immunological synapse (Babich et al., 2012; Yi et al., 2012), axon guidance in neuronal growth cones (Brown and Bridgman, 2003; Lin et al., 1996; Medeiros et al., 2006)), cell migration (Alexandrova et al., 2008; Cai et al., 2006; Maiuri et al., 2015), microtubule buckling and breakage (Waterman-Storer and Salmon, 1997), and the alignment of ligand-activated integrins at focal adhesions (Swaminathan et al., 2017)). All these systems share the general feature that the retrograde flow in the lamellipodia/leading-edge area is driven by actin polymerization occurring at or near the PM whereas the flow in the lamella/sub-leading-edge area is governed by a mechanism involving myosin-II and other types of F-actin structures. While the consensus on the role of actin polymerization is strong, the function and mechanism of myosin-II is less clear. For example, for the retrograde flow at the immunological synapse, one study suggests that myosin-II is not directly involved in driving the flow but is required for the long-term integrity of the synapse (Babich et al., 2012) whereas another study suggests that myosin-II and F-actin arcs at the lamella form concentric actomyosin bundles to power the flow (Yi et al., 2012). However, it is unclear how the actomyosin bundles are tethered to the PM to drive the retrograde flow. Similarly, the architecture and tethering mechanism of myosin-II in the lamella of neuronal growth cones and migrating cells remains unclear. It is also not fully understood whether and how different isoforms (IIA, IIB, and IIC) of myosin-II make distinct contributions to the retrograde flow in a cell type-specific manner, and whether the myosin heads gliding on the flowing actin filaments are from myosin-II bipolar filaments or even active monomers (Sanborn et al., 2011; Shutova et al., 2014).

The mechanism of RACF in the budding yeast *S. cerevisiae* offers an attractive model to understand the role of myosin-II in retrograde flow in other systems. Myo1, the sole myosin-II heavy chain in *S. cerevisiae*, was known to facilitate RACF by localizing at the bud neck and gliding on the flowing actin cables from the bud tip towards the mother (Huckaba et al., 2006), but the tethering mechanism of Myo1 and its precise timing of function during the cell cycle were not clearly understood. In this study, we have demonstrated that Bni5 links Myo1 to the septin hourglass before the onset of cytokinesis to facilitate RACF. Furthermore, our analysis indicates that Bni5 stably associates with the septin hourglass by binding to septin filaments via its C-terminal Ext-CC2-CC3 domain and this binding is likely mediated by the terminal septin subunits Cdc11 and Shs1. We have also demonstrated that Bni5 dynamically associates with Myo1 by binding to the mTD1 in Myo1 tail via its N-terminal CC1 (Fang et al., 2010). As Myo1 is mobile at the division site before anaphase and is progressively immobilized during anaphase and cytokinesis (Wloka et al., 2013), it is likely that Myo1 participates in RACF in the form of active monomers while functioning in cytokinesis in the filamentous form as indicated by our EM studies (Chen et al., 2020; Ong et al., 2014)). Thus, we have defined a specific biochemical pathway for the tethering, timing, and function of Myo1 in RACF during the cell cycle.

Interestingly, Cdc11 and Shs1 appear to share a common ancestor and their divergence from the common ancestor is restricted to budding yeasts (Takagi et al., 2021). The exclusive conservation of Bni5, Cdc11, and Shs1 in the budding yeast species and the specific requirement of both Cdc11 and Shs1 for the localization of Bni5 at the bud neck suggest their co-evolution. The genome sequence of CC1 region is less conserved than that of C-terminal Ext-CC2-CC3 region (Finnigan et al., 2015a) (this study). However, modeling of protein structures in *Saccharomycetaceae* species still predicts a short coiled-coil region at their N-termini similar to the CC1 of Bni5 in *S. cerevisiae* (e.g., *Ashbya gossypii* Q754P5, data available at https://alphafold.ebi.ac.uk/). Given the more stringent conservation of septins among species than the tails of myosin-IIs (Pan et al., 2007; Wang et al., 2020), it is possible that the divergence of CC1 among budding yeast species is due to the co-evolution of Bni5 and myosin-II tail to fit unique requirements or conditions experienced by each species.

Given the strict conservation of the septin-Bni5-myosin-II pathway in the budding yeast species and its role in promoting RACF in *S. cerevisiae*, thereby mediating the asymmetric inheritance of mitochondrial fitness and mitochondria-tethered protein aggregates (Higuchi et al., 2013; Zhou et al., 2014) (this study), it is likely that this pathway has evolved to control aging or cellular health of all these budding yeasts where polarized growth and asymmetric cell division is the way of life.

Despite the budding yeast-specific feature of the septin-Bni5-myosin-II pathway, the general similarity in the spatial distribution of the flowing actin filaments (in lamellipodia or bud cortex) in relation to the myosin-II (in lamella or at the bud neck) and the conserved nature of septins and myosin-II could still allow the mechanistic analysis of the septin-Bni5-myosin-II pathway in budding yeast to inform similar analyses in other retrograde flow systems. Particularly interesting is the possibility that septins and myosin-II may act together via a cell type-specific linker protein (the functional equivalent of Bni5) to drive retrograde flow in mammalian cells.

## Materials and methods

### Yeast media and culture conditions

Standard culture media and genetic techniques were used (Guthrie and Fink, 1991). Yeast strains were grown routinely at 25°C in synthetic complete (SC) minimal medium lacking specific amino acid(s) and/or uracil or in rich medium YM-1 (Lillie and Pringle, 1980). Stock solutions of 20 mM LatA (in DMSO, FUJIFILM Wako Pure Chemical, Osaka, Japan), 1 mM MitoTracker Red CMXRos (in DMSO, Cell Signaling Technology, Danvers, MA, USA) and 1% (w/v) CW (in distilled water, Sigma, St. Louis, MO, USA) were diluted into media at the indicated final concentrations.

### Constructions of strains

New strains were constructed either by integrating a plasmid carrying a modified gene at a genomic locus or by transferring a deletion or tagged allele of a gene from a plasmid or from one strain to another via PCR amplification and yeast transformation (Lee et al., 2013; Longtine et al., 1998) (see footnotes in **Table S1**).

### Primers and plasmids

All plasmids and PCR primers used in this study were listed in **Table S2** and **Table S3**, respectively. All PCR primers were purchased from Integrated DNA Technologies (Coralville, IA, USA). All new constructs were validated by sequencing performed at the DNA Sequencing Facility, University of Pennsylvania. Plasmids pAG25 (Goldstein and McCusker, 1999), YIp128-CDC3-GFP (Gao et al., 2007), pFA6a-link-yoEGFP-SpHIS5 and pFA6a-link-yoTagRFP-T-CaURA3 (Lee et al., 2013), pFA6a-link-ymScarlet-I-CaURA (Marquardt et al., 2020), pMAL-MYO1-mTD1 (Fang et al., 2010), pRG205MX (Gnugge et al., 2016), proHIS3-ymScarlet-I-TUB1-tTUB1-HPH (Ghanegolmohammadi et al., 2021), pRS316-N-MYO1-GFP (Caviston et al., 2003), and pUG36-GFP-Ecm25-xACT(536–588aa) (Duan et al., 2021) were described previously.

The following plasmids were kindly provided by the indicated colleagues: bWL722 (Markus et al., 2015) (Wei-Lih Lee, Dartmouth College); pCOLA-Duet-[His less]-Shs1 (Garcia et al., 2011), pMVB128, and pMVB133 (Versele et al., 2004) (Jeremy Thorner, University of California, Berkeley); pFA6a-URA3-KanMX6 (Onishi et al., 2013) (John Pringle, Stanford University); pUG34 and pUG36 (Johannes H. Hegemann, Heinrich Heine University Düsseldorf); pYES2-Htt103Q-GFP (Song et al., 2014) (Thomas Nyström, University of Gothenburg); YCp50-MYO1 (Susan Brown, University of Michigan); and YIplac128-pCUP2-gcR7-ymScarlet-I-tADH1 (Takashi Ito, Kyushu University).

The following plasmids were generated for this study: To generate pET-His6-Sumo-Elm1_FL_, pET-His6-Sumo-Elm1_1-420_, or pET-His6-Sumo-Elm1_421-640_, a DNA fragment containing *ELM1(FL)*, *elm1(1-420)*, or *elm1(421-640)* was amplified by PCR using plasmid pUG36-ELM1(FL), pUG36-ELM1(1-420), or pUG36-ELM1(421-640) (Our lab stocks) as the template DNA and the pair of primers Elm1-FL-F and Elm1-FL-R, Elm1-N-F and Elm1-N-R, or Elm1-C-F and Elm1-C-R, respectively. The resultant PCR products were subcloned into BamHI- and SspI-digested pET-His6-Sumo-TEV-LIC (Addgene # 29659) using in-fusion cloning kit (Takara, Kusatsu, Japan). To generate pFA6a-link-GBP-CaURA3, a ∼0.4-kb DNA fragment containing *GBP-6xHis* was amplified by PCR using the plasmid pRS315-INN1-C2-mApple-GBP-6xHis (lab stock) as the template DNA and the pair of primers GBP-F and GBP-R, and then subcloned to replace ∼0.7-kb PacI-AscI region of pFA6a-link-yoTagRFP-T-CaURA3 using in-fusion cloning kit. To generate pFA6a-link-yomApple-GBP-CaURA, a ∼1.7-kb DNA fragment containing *inn1*-*mApple-GBP-6xHis* was amplified by PCR using the plasmid pRS315-INN1-C2-mApple-GBP-6xHis as the template DNA and the pair of primers P45 and P425. The resultant PCR product was digested with PacI and AscI to acquire ∼1.2 kb *mApple-GBP-6xHis* fragment, and then the fragment was subcloned to replace ∼0.7-kb PacI-AscI region of pFA6a-link-yoTagRFP-T-CaURA3 using T4 DNA ligase (New England Biolabs, Ipswich, MA, USA). To generate, pGEX-4T-1-BNI5(1-448), pGEX-4T-1-BNI5(1-40), pGEX-4T-1-BNI5(306-393), pGEX-4T-1-BNI5(340-448), or pGEX-4T-1-BNI5(306-448) a ∼1.4-kb, ∼0.2-kb, ∼0.3-kb, ∼0.4-kb, or ∼0.5-kb DNA fragment containing *BNI5(1-448)*, *bni5(1-40)*, *bni5(306-393)*, *bni5(340-448)*, or *bni5(306-448)* was acquired by BamHI- and XhoI-digestion of pUG36-BNI5(FL), pUG36-BNI5(1-40), pUG36-BNI5(309-393), pUG36-BNI5(340-448), or pUG36-BNI5(306-448), respectively. The resultant DNA fragments were subcloned to replace ∼0.9-kb BamHI-XhoI insert of pGEX-4T-1-SSO1 (lab stock) using T4 DNA ligase. To generate pRG205MX-proBNI5-yEGFP, a DNA fragment containing 800 bp of promoter region of *BNI5* and *yEGFP* followed by linker peptide (SRTSGSPGL) was amplified by PCR using the chromosomal DNA of YEF10276 (Marquardt et al., 2020) as the template and the pair of primers P1581 and P1582. The resultant PCR product was then subcloned into SacI- and XbaI-digested pRG205MX using in-fusion cloning kit. To generate pRG205MX-proBNI5-yEGFP-BNI5(FL), pRG205MX-proBNI5-yEGFP-BNI5(1-40), pRG205MX-proBNI5-yEGFP-BNI5(306-339), pRG205MX-proBNI5-yEGFP-BNI5(306-448), pRG205MX-proBNI5-yEGFP-BNI5(340-448), pRG205MX-proBNI5-yEGFP-BNI5(41-448), and pRG205MX-proBNI5-yEGFP-BNI5(Δ41-305), A DNA fragment containing *BNI5(FL)*, *BNI5(1-40)*, *BNI5(306-339)*, *BNI5(306-448)*, *BNI5(340-448)*, *BNI5(41-448)*, or *BNI5(Δ41-305)* was acquired by EagI and BamHI digestion of pUG36-BNI5(FL), pUG36-BNI5(1-40), pUG36-BNI5(306-339), pUG36-BNI5(306-448), pUG36-BNI5(340-448), pUG36-BNI5(41-448), or pUG36-BNI5(Δ41-305), and then subcloned into EagI- and BamHI-digested pRG205MX-proBNI5-yEGFP using T4 DNA ligase. To generate pUG34-ymScarlet-I, a ∼0.7-kb DNA fragment containing *ymScarlet-I* was amplified by PCR using the plasmid YIplac128-pCUP2-gcR7-ymScarlet-I-tADH1 as the template and the pair of primers P1583 and P1584, and then assembled with XbaI linearized pUG34 (CEN *HIS3* GFP) by gap repair cloning (Oldenburg et al., 1997) to replace the GFP region of pUG34 with ymScarlet-I. To generate pUG34-ymScarlet-BNI5(394-448), a ∼0.2-kb DNA fragment containing *bni5(394-448)* was acquired from BamHI- and XhoI-digested pUG36-(394-448), and then ligated into BamHI- and XhoI digested pUG34-ymScarlet-I. To generate YIp128-proACT1-GFP-ECM25-(536-588AA)-tADH1, a ∼0.2 kb DNA fragment containing *ecm25(536-588)* was amplified by PCR using plasmid pUG36-GFP-Ecm25-xACT(536–588aa) as the template and the pair of primers P254 and P549. The ∼5.8-kb DNA fragment of plasmid vector, which can express gene of interest under *ACT1* promoter, was prepared by inverse PCR using plasmid YIp128-proACT1-PKC1C1-GFP-tADH1 (lab stock) as the template and the pair of primers P255 and P550. Two resultant PCR products were then subcloned using in-fusion cloning kit. The pUG36-BNI5* plasmid series that express different *BNI5* fragments with N-terminally tagged GFP [e.g., pUG36-BNI5(FL)] were constructed by gap repair cloning or digestion and ligation cloning (see footnotes of **Table S2**).

### Live cell imaging and data analysis

Quantitative time-lapse imaging analysis was performed as described previously with slight modifications (Okada et al., 2021a). For time-lapse microscopy, cells were cultured to exponential phase at 25°C in SC medium, briefly sonicated at 15% power for 5 sec to declump (model Q55, Qsonica, Newtown, CT, USA), concentrated by centrifugation, and spotted onto a concanavalin-A coated glass bottom chamber dish. Once cells were attached to the bottom, excess cells were removed by discarding the supernatant and 1mL fresh SC medium was added to the dish. For samples imaged at 37°C, cells were pre-cultured at 37°C for one hour prior to harvesting for time-lapse analysis. Imaging was performed at room temperature (23°C) or 37^°^C controlled by the OKO temperature control system (Okolab, Ottaviano, NA, Italy). Images were acquired by spinning-disk confocal microscopy using a Nikon microscope (model Eclipse Ti2-U, Nikon, Tokyo, Japan) equipped with a 100x/1.49NA oil objective (model CFI Apo TIRF 100x, Nikon) and a confocal scanner unit (model CSU-X1, Yokogawa, Tokyo, Japan). An EMCCD camera (model Evolve 512 Delta, Photometrics, Tucson, AZ, USA) was used for capture. Solid-state lasers for excitation (405 nm for CW, 488 nm for GFP and 561 nm for RFP) were housed in a launch (model ILE-400, Spectral Applied Research, Richmond Hill, ON, Canada). The imaging system was controlled by MetaMorph version 7.10.4.431 (Molecular Devices, San Jose, CA, USA). All time-lapse images, except for actin cables (GFP-xACT), were taken every 1, 1.5, or 2 min with 11 z-stacks with a step-size of 0.8 m. Images of GFP-xACT were taken every 1 sec at a single fixed focal plane. A sum or max projection was created with NIH ImageJ (1.53t). The quantification of fluorescence intensities was performed as described previously (Okada et al., 2021a). In brief, the fluorescent intensity at the division site was calculated by subtracting the intensity in the background area from the that of the division site. To calculate the constriction rate, we manually measured the myosin ring diameter during constriction from max projection images acquired through time-lapse imaging. Then, we calculated the slope of the diameter curve from 3 or 4-time points including midpoint of constriction. To determine the rate of actin cable flow, we manually marked an endpoint of an extending actin cable and measured the distance between endpoints in each consecutive time frames. Data analyses were performed with Microsoft Excel, GraphPad Prism 9.4.1, and R (ver. 3.0.1).

### FRAP

FRAP analysis was performed as described previously (Okada et al., 2019). Imaging samples were prepared as described above (see **Live cell imaging and data analysis**). The imaging system used consists of a spinning-disk confocal scanner unit (model CSU-X1, Yokogawa) and a microscope (model IX81, Olympus, Tokyo, Japan) equipped with a 100×/1.40 NA oil objective (model UPlanSApo 100x/1.40 Oil, Olympus) and an EMCCD camera (model iXon X3 897, Andor Technology, Belfast, UK). MetaMorph ver. 7.8.10.0 (Molecular Devices) was used for hardware control and image acquisition. Diode lasers (488 nm for GFP and 561 nm for RFP) controlled via laser merge module (model LMM5, Spectral Applied Research, Richmond Hill, Ontario, Canada) were used for excitation. Images were taken at room temperature with 11 z-stacks with a step size of 0.7 µm. To induce photobleaching, a diode-pumped 405 nm laser (model DL405-050-O CrystaLaser, Reno, NV, USA) was applied to a defined subcellular region. Maximum projections were created and analyzed with NIH ImageJ. In ImageJ, a polygon was drawn encircling the bleached area to calculate the integrated density within the area over time. Data of recovery curve were analyzed with GraphPad Prism 9. The estimated maximum amount of recovery (max) and half-time of recovery (t_1/2_) were determined using the one phase-association function in the GraphPad.

### Inheritance of Htt103Q protein aggregates

Cells were cultured at 25°C until they reached the exponential growth phase in SC-Ura+2% Raf medium (using raffinose instead of glucose). Subsequently, they were transferred to SC-Ura+2% Gal medium (using galactose instead of glucose) and cultured at 25°C to induce overexpression of Htt103Q. After 4 hours of protein induction, the cells were transferred to SC-Ura medium. For mitochondrial staining, MitoTracker Red CMXRos was added (100 nM) and incubated in the dark for 30 min to stain the mitochondria. Afterward, the cells were washed three times with SC-Ura medium. Imaging was performed as described above (see **Live cell imaging and data analysis**).

### Yeast growth assay

Spot assay was performed to examine cell growth under different conditions. Cells were cultured in SC or YPD medium at 25°C for 18 hours, and cell culture was diluted with fresh YPD medium to the 0.1 OD_600_. The cell suspension was subjected to 10-fold serial dilutions and inoculated as 5 µl of spot onto SC- or YPD-plate. After incubation at 25°C or 37°C for 2 or 3 days, the cell growth on the plate was recorded.

### Purification and in vitro binding of recombinant proteins

For testing Bni5 interactions with Elm1 and Myo1, Rosetta DE3 cells transformed with pGEX-4T1 (GST alone), pGEX-4T1-Bni5-FL (GST-Bni5-full length), pGEX-4T1-Bni5-CC1(aa1-40), pGEX-4T1-Bni5-Ext-CC2(aa306-393), pGEX-4T1-Bni5-CC2-CC3(aa340-448), pGEX-4T1-Bni5-Ext-CC2-CC3(aa306-448), pET-His6-Sumo-TEV-LIC (6xHis-SUMO alone), pET-His6-Sumo-Elm1-N(aa1-420), pET-His6-Sumo-Elm1-C(aa421-640), pMAL-C2 (MBP only) or pMAL-C2-Myo1-mTD1 were grown to an OD600 0.6-1.0 before being induced for 3 hours with 0.3 mM IPTG (Lab Scientific, Highlands, NJ, USA) at 25°C. Cells were then lysed by sonication 6 times for 15 sec each in one of the following buffers each containing the Protease Inhibitors Cocktail tablets (Roche) and 1 mg/mL lysozyme (L6876, Sigma Aldrich): the GST lysis buffer (50 mM Tris-HCl, pH 7.5, 300 mM NaCl, 1.25 mM EGTA, 1 mM DTT and 0.1 % NP-40), the 6xHis-SUMO lysis buffer (50 mM Tris-HCl, pH 7.5, 300 mM NaCl, 1.25 mM EGTA, 1 mM DTT, 0.1% NP-40, and 15 mM imidazole), or the MBP lysis buffer (20 mM Tris-HCl, pH 7.5, 200 mM NaCl, 1.25 mM EGTA, 1 mM DTT, and 0.1% NP-40). The resultant lysates were then centrifuged at 24,000 X g for 30 min at 4°C. The supernatants were then incubated with either Glutathione Sepharose 4B (GE Healthcare, Chicago, IL, USA), Complete His-Tag Purification Resin (Roche, Basel, Switzerland), or Amylose Resin (NEB, 1 L, USA), that had been prewashed with respective lysis buffer, for 1 hour at 4°C. The beads were then washed five times with respective lysis buffer. GST-tagged proteins were kept with Glutathione Sepharose 4B and resuspended in Bead storage buffer (50 mM Tris-HCl, pH 7.5, 5 mM MgCl_2_, 25% glycerol), while 6xHis-SUMO-tagged proteins and MBP-tagged proteins were eluted by the His elution buffer (50 mM Tris-HCl, pH7.5, 300 mM NaCl, 1.25 mM EGTA, 1 mM DTT, 0.1% NP-40, and 300 mM imidazole) or the MBP elution buffer (20 mM Tris-HCl, pH 7.5, 200 mM NaCl, 1.25 mM EGTA, 1 mM DTT, 0.1% NP-40, and 10 mM Maltose), respectively. Protein concentrations were determined by standard curve intensity measurements from Coomassie blue-stained bovine serum albumin (A7617, Sigma Aldrich, St. Louis, MO, USA) of known concentrations.

For the in vitro binding assay, 3.5 μg of GST, GST-Bni5-FL, GST-Bni5-CC1, GST-Bni5-Ext-CC2, GST-Bni5-CC2-CC3, or GST-Bni5-Ext-CC2-CC3 was incubated with 3.5 μg of either His6-Sumo, His6-Sumo-Elm1-C, His6-Sumo-Elm1-N, MBP or MBP-Myo1-mTD1 for 1 hour with rotation in binding buffer (20 mM MOPS, pH7.0, 1 mM EGTA, 150 mM NaCl, 1 mM DTT, 0.1%NP-40). After centrifugation, the beads were then washed five times with fresh binding buffer before being extracted with 30 µL of 2X Laemmli Buffer (Bio-Rad Laboratories, Hercules, CA, USA). 10 μL of the samples were separated via SDS-PAGE and stained with SimplyBlue safe stain while 1 μL of the samples were separated via SDS-PAGE, and then transferred to a PVDF membrane before immunoblotting with the anti-6xHis (1:5000 dilution, ab18184, abcam) antibody or MBP (1:10,000, M1321, Sigma Aldrich) primary antibody, respectively, and HRP-labeled secondary antibody and ECL reagents from the Pierce Fast Western kit (Thermo Scientific).

For testing Bni5 interaction with septin filaments, plasmids pMVB128 (Cdc10, His6-Cdc12) and pMVB133 (Cdc3, Cdc11) with or without pCOLA-Duet-[His less]-Shs1 were co-transformed into *E. coli* strain BL21(DE3) to generate 4-septin or 5-septin complexes, respectively. The transformed cells were grown to an OD600 0.6-1.0 before being induced for 4 hours with 1.0 mM IPTG (Lab Scientific, Highlands, NJ, USA) at 37°C, harvested by centrifugation, washed once with PBS, and stored at -80°C. Frozen cells were lysed in Renz buffer (300 mM NaCl, 2 mM MgCl_2_, 15 mM imidazole, 12% (v/v) glycerol, 50 mM Tris-HCl, pH 7.5, Protease Inhibitors Cocktail tablets, and 1 mg/mL lysozyme) by sonication 10 times for 20 sec each. The resultant lysates were then centrifuged at 24,000 X g for 30 min at 4°C. The supernatants were then incubated with the Complete His-Tag Purification Resin (Roche, Basel, Switzerland) that had been prewashed with Renz buffer, for 1 hour at 4°C. The beads were then washed five times with Renz buffer. Protein complexes were then eluted with the elution buffer (300 mM NaCl, 2 mM MgCl_2_, 15 mM imidazole, 12% (v/v) glycerol, 50 mM Tris-HCl, pH 7.5, and 300 mM imidazole). It is worth noting that GST-tagged proteins underwent the same protein purification process as described as above and were eluted with the GST elution buffer (50 mM Tris-HCl, pH 7.5, 300 mM NaCl, 1.25 mM EGTA, 1 mM DTT and 0.1% NP-40, and 10 mM Glutathione) for binding assay with septin filaments. Protein concentrations were determined by standard curve intensity measurements from Coomassie blue-stained bovine serum albumin of known concentrations.

Septin filament formation was performed using an established protocol (Renz et al., 2013) with some modifications: purified septin complexes in the high salt buffer (300 mM NaCl) was diluted to the low salt buffer (50 mM NaCl) by adding the dilution buffer (2 mM MgCl2, 15 mM imidazole, 12% (v/v) glycerol, 50mM Tris-HCl, pH 7.5) and then kept at 4°C for 16 hours to allow filament assembly. Septin complexes and filaments were then concentrated using Centrifugal Filter (Amicon^R^ Ultra-15, Sigma Aldrich).

For the in vitro binding assay, 1.5 μg of GST, GST-Bni5-FL, GST-Bni5-CC1, GST-Bni5-Ext-CC2, GST-Bni5-CC2-CC3, or GST-Bni5-Ext-CC2-CC3 was incubated with 1.5 μg of the concentrated 4-septin [Cdc11-Cdc12(His6)-Cdc3-Cdc10-Cdc10-Cdc3-Cdc12(His6)-Cdc11] or 5-septin [a mixture of Cdc11-Cdc12(His6)-Cdc3-Cdc10-Cdc10-Cdc3-Cdc12(His6)-Cdc11 and Shs1-Cdc12(His6)-Cdc3-Cdc10-Cdc10-Cdc3-Cdc12(His6)-Shs1] complexes/filaments in the binding buffer (50 mM NaCl, 2 mM MgCl2, 15 mM imidazole, 12% (v/v) glycerol, 50mM Tris-HCl, pH 7.5) with rotation for 1 hour at room temperature. The reactions were subjected to centrifugation at 100,000 X g for 1 hour at 4°C to pellet the septin filaments and the associated proteins. The pellets were washed once with fresh binding buffer before being extracted with 35 µL of 2X Laemmli Buffer. 10 μL of the samples were separated via SDS-PAGE and stained with SimplyBlue safe stain while 1 μL or 0.2 μL of the samples were separated via SDS-PAGE, and then transferred to a PVDF membrane before immunoblotting with the anti-GST (1:3000 dilution, ab92, abcam) or anti-SHS1 (1:5000, a gift from Doug Kellogg) primary antibody, and visualized as described above.

### Quantification and statistical analysis

For the statistical analyses of intensities at the division site (related to **Figs. 4 K, 6 D, 6 E, 7 A, and S3 D**), protein turnover kinetics (**Fig. 7 B**), rates of signal drop (**Fig. 7 F**), RACF (**Fig. 8 C**), and Myo1 constriction (**Fig. 9 C**), and pattern of Htt103Q inheritance (**Fig. 8 F**), a two-sided unpaired t-test (assuming unequal variances) was performed. For the correlation analyses of protein accumulation kinetics (**Fig. S5 D**) and protein turnover kinetics (**Fig. S1 C**), a Pearson correlation coefficient between mean values from selected time points (-24 to +15 min) and (0 to 360 sec) was calculated, respectively. “n” refers to the number of cells analyzed unless indicated otherwise.

## ABBREVIATIONS

AMR: actomyosin ring
Ext: extended sequence required for CC2 localization FL full-length
FRAP: fluorescence-recovery-after-photobleaching GBP GFP-binding protein
HDR: hourglass-to-double ring
LatA: latrunculin A
mTD1: minimal targeting domain 1
MID: middle disordered region
RACF: retrograde actin cable flow
WT: wild-type

## SUPPLENTAL MATERIAL

Five supplemental figures are presented, each accompanied by its corresponding figure legend. Fig. S1 depicts the septin accumulation kinetics in WT and *elm1Δ* cells, Myo1 FRAP analysis, and the correlation coefficients between the turnover rates of N- or C-tagged Bni5, septins, Elm1, and Myo1 at the division site. Fig. S2 depicts the evolutionary conservation of Bni5 and its specific domains among fungal species. Fig. S3 shows the functional relationship between the CC1 of Bni5 and the mTD1 of Myo1. Fig. S4 depicts the ability of different Bni5 C-terminal fragments to co-localize with the septins, function as a dosage suppressor of a septin mutant, and interact with septin filaments in vitro and also shows the presence of Cdc11-containing filaments in *shs1Δ* cells. Fig. S5 illustrates the growth and morphological phenotypes of *bni5Δ* in different laboratory strains of *S. cerevisiae,* the change in accumulation kinetics of Myo1 and Chs2 in the absence of Bni5, the impact of Bni5-Myo1 interaction on protein localization, and the synthetically enhanced effect on Myo1 localization between *bni5Δ* and a *myo1* truncation allele (*myo1-1797*).

## DATA AVAILABILITY STATEMENT

The data are available from the primary corresponding author (Erfei Bi via) upon reasonable request.

## ACKNOWLEDGEMENTS

We thank Charles Boone, Scott D Emr, Christopher G Burd, Wei-Lih Lee, Jeremy Thorner, John Pringle, Johannes H. Hegemann, Thomas Nyström, Susan Brown, and Takashi Ito for plasmids and strains; Douglas Kellogg for antiShs1 antibody; Mikael Garabedian and the former and present members of the Bi lab, especially Xiaodong Fang, Carsten Wloka, Yuwen Dong, and Junya Hayase, for stimulating discussions and critically reading the manuscript. The authors declare no competing financial interests. This work was supported by National Institutes of Health grants (GM115420 and GM116876 to E.B.)

## AUTHER CONTRIBUTIONS

Conceptualization, H.O. and E.B.; Methodology, H.O. and E.B.; Investigation, H.O., X.C., K.W., and J.M.; Writing, H.O. and E.B.; Supervision, E.B.

