## Supplemental Figures S1-S5 and Tables S1-S3 for "Bni5 tethers myosin-II to septins to enhance retrograde actin flow and the robustness of cytokinesis"

**SUPPLEMENTAL FIGURE LEGENDS**

**Figure S1. Accumulation and turnover kinetics of Bni5 and its associated proteins at the bud neck. Related to Figure 2.**

(A) Kinetics of Cdc3-mCherry in WT and *elm1*Δ strains carrying the indicated GFP-tagged *BNI5*. The same strains were used in **Fig. 2 C** and **2 D**. Bold lines and associated shaded bands represent mean and SD values, respectively.

(B) FRAP analysis on Myo1-GFP. Half of the ring from each cell was photo-bleached, and fluorescence recovery in the bleached region (gray symbol) and unbleached region (green symbol) was followed over time. Top: Lines, symbols, and error bars represent regression curves, mean values, and SD values, respectively. Bottom: Montages of cells were created from time-lapse series taken with a 10-sec interval. Strain used is YEF11385 (*MYO1-GFP mScarlet-TUB1*).

(C) Correlation coefficients between the turnover rates of N- or C-tagged Bni5, septins (Cdc10), Elm1, and Myo1 at the division site. Values used for this analysis are from **Figs. 1 G, 2 F, and S1 B**.

**Figure S2. Evolutional conservation of distinct regions in Bni5. Related to Figure 3.**

Fungal BLAST analysis was performed in the SGD (https://www.yeastgenome.org/blast-fungal) using the *S. cerevisiae* Bni5 sequence as a query to search datasets of indicated fungal species at the default setting. Green boxes indicate a region with significant homology. The presence of the *BNI5* homolog was determined by a database search of PhylomeDB (http://phylomedb.org/). The dendrogram shown is modified from {Cuomo, 2010 #10341}.

**Figure S3. Impact of the CC1 of Bni5 and the mTD1 of Myo1 on the localization of both proteins at the bud neck. Related to Figure 4.**

(A) Kinetics of GFP-Bni5(FL) and GFP-Bni5(ΔCC1) during bud formation. Bold lines and associated shaded bands represent mean and SD values, respectively. Strains used are as follows: YEF11546 [*GFP-BNI5(FL) ymScarlet-TUB1*], and YEF11600 [*GFP-bni5(41-448) ymScarlet-TUB1*].

(B) Bni5(FL) rescues Myo1 localization at the division site before cytokinesis in *bni5*Δ cells. Montages of GFP-Bni5(FL) or GFP with respect to Myo1-mScarlet in *bni5*Δ cells were created from time-lapse series taken with a 2.5-min interval. Strains used are as follows: YEF11029 (*bni5*Δ *MYO1-ymScarlet* [*GFP-BNI5(FL)*]) and YEF11033 (*bni5*Δ *MYO1-ymScarlet [GFP]*).

(C) Accumulation kinetics of GFP-Bni5(FL) and GFP-Bni5(CC1) shortly before and during cytokinesis. Bold lines and associated shaded bands represent mean and SD values, respectively. Strains used are as follows: YEF11546 [*GFP-BNI5(FL) ymScarlet-TUB1*] and YEF11547 [*GFP-BNI5(1-40) ymScarlet-TUB1*].

(D) Impact of deleting the CC1 of Bni5 on the bud neck localization of Myo1-mTD1-GFP. Left: Cells cultured in SC-Ura medium until the exponential growth phase were subjected to imaging. Arrowheads represent bud neck signal of Myo1-mTD1-GFP. Strains used are as follows: YEF11771 (*myo1-mTD1-GFP bni5Δ* [*URA3 BNI5(FL)*]), YEF11774 (*myo1-mTD1-GFP bni5Δ* [*URA3 bni5(41-448)*]), and YEF11775 (*myo1-mTD1-GFP bni5Δ* [*URA3*]). Right: The intensity of Myo1-mTD1-GFP at the bud neck in cells with a small or medium bud (S/G2 cells) was scored. Green circles, blue bars, and gray error bars represent all data points, mean values, and SD values, respectively.

(E) Forced tethering of Bni5(CC1) to the septins at the bud neck using the GBP-GFP system. Cells in the exponential growth phase were subjected to imaging. Strains used are as follows: YEF11215 (*CDC11-GFP bni5(1-40)-mApple-GBP*), YEF11250 (*CDC11 bni5(1-40)-mApple-GBP*). YEF11214 (*CDC11-GFP bni5Δ::mApple-GBP*), and YEF11249 (*CDC11 bni5Δ::mApple-GBP*).

**Figure S4. The CC2 domain and adjacent regions of Bni5 are responsible for the interaction and function with the septins. Related to Figure 5.**

(A) The normally expressed (or uninduced) Ext, CC2, or CC3 of Bni5 by themselves do not localize to the bud neck. Cells cultured in the SC-Ura medium until the exponential growth phase were subjected to imaging. Strains used are as follows: YEF11053 (*bni5Δ CDC3-mCherry* [*GFP-BNI5(FL)*]), YEF11181 (*bni5Δ CDC3-mCherry* [*GFP-bni5(306-339)*]), YEF11041 (*bni5Δ CDC3-mCherry* [*GFP-bni5(340-393)*]), and YEF11042 (*bni5Δ CDC3-mCherry* [*GFP-bni5(394-448)*]).

(B) Overexpressed (or induced) Ext localizes to the bud neck in an Elm1-independent but Cdc11 and Shs1-dependent manner. Expression of Bni5 fragments was induced by culturing in SC-Ura-Met medium for five hours. Arrowheads indicate the bud neck localization of GFP-Ext. Strains used are as follows: YEF11181, YEF11041, YEF11042, YEF11048 (*bni5Δ CDC3-mCherry* [*GFP*]), YEF11196 (*bni5Δ elm1Δ CDC3-mCherry* [*pUG36-BNI5(306-339)*]), and YEF11926 (*bni5Δ shs1Δ cdc11Δ CDC3-mCherry* [*pUG36-BNI5(306-339)*]).

(C) The ability of Bni5 fragments to function as a dosage suppressor of the temperature-sensitive septin mutant (*cdc12-6*). Different fragments of Bni5 were analyzed using the pUG36-Bni5* plasmid series (**Table S2**). Left: Alleles in sky blue or black indicates the suppression or non-suppression of the growth defect at the restrictive temperature, respectively. Bold lines and dashed lines indicate the presence and absence of the region of Bni5 in relevant alleles. Right: spot assay result documented after three days of incubation at 25°C or 32°C. Strains used are summarized in **Table S1**; all strains with denotation with [z] except YEF11362 and YEF11363.

(D) Western blot using an anti-Shs1 antibody to show the presence of Shs1 in the 5-septin, but not the 4-septin, complexes used in our in vitro binding experiments.

(E) Western blots using an anti-GST antibody to demonstrate the ability of different Bni5 fragments to bind septin filaments in vitro. Due to poor transfer of large-sized proteins from polyacrylamide gel to PVDF membrane, the GST-FL (Bni5) could be observed only when more samples were loaded for the Western blot analysis (left).

(F) The presence of Cdc11 at the bud neck in *shs1Δ* cells. Cells of the strain YEF11436 (*CDC11-mCherry shs1Δ*) were grown to the exponential phase and then subjected to imaging.

**Figure S5. Bni5 increases the robustness of the AMR against genetic perturbation during cytokinesis. Related to Figure 9.**

(A) Deletion of *BNI5* does not cause growth defects in various laboratory strains. The result of the spot assay on YPD plates was documented after two days of incubation at 25°C or 37°C. Strains used are as follows: YEF11851 (W303 *bni5*Δ), YEF11852 (SEY6210 *bni5*Δ), and YEF11853 (BY4742 *bni5*Δ).

(B) Deletion of *BNI5* does not cause defects in cytokinesis or cell morphology. Cells were cultured in YM-1 medium at 25°C or 37°C for 24 hours and subjected to imaging. Strains used are the same as (A).

(C) Accumulation kinetics of the Myo1-mTD1Δ vs. Myo1 in *bni5*Δ cells during cytokinesis. Intensities of indicated proteins at the bud neck were measured from time-lapse series taken with a 1.5-min interval. Bold lines and associated shaded bands represent mean and SD values, respectively. Strains used are as follows: YEF11441 (*MYO1-GFP mScarlet-TUB1*), YEF11442 (*myo1-mTD1*Δ*-GFP mScarlet-TUB1*), and YEF11443 (*bni5*Δ *MYO1-GFP mScarlet-TUB1*).

(D) Correlation coefficients between the accumulation kinetics of indicated proteins during cytokinesis. Nodes and red and brown lines between nodes represent indicated proteins, a significant correlation at $p<0.001$ and $p<0.01$, respectively. Numbers associated with the lines represent the correlation coefficient between nodes. Values used for analysis are from **Figs**. **S3 C and S5 C**.

(E) Accumulation kinetics of the Chs2 in WT and *bni5Δ* cells during cytokinesis. Intensities of Chs2-GFP at the bud neck were measured from time-lapse series taken with a 1.5-min interval. Bold lines and associated shaded bands represent mean and SD values, respectively. Strains used are as follows: YEF10692 (*CHS2-GFP mScarlet-TUB1*) and YEF11390 (*bni5*Δ *CHS2-GFP mScarlet-TUB1*).

(F) Synthetically enhanced effect on Myo1 localization by *bni5Δ* and the tail truncation of Myo1. Left: Accumulation kinetics of Myo1 or tail-truncated Myo1 in WT and *bni5*Δ cells during cytokinesis. Intensities of indicated proteins at the bud neck were measured from time-lapse series taken with a 1.5-min interval. Bold lines and associated shaded bands represent mean and SD values, respectively. Right: Montages of indicated proteins at the bud neck were created from selected frames. Strains used are as follows: YEF11529 (*GFP-MYO1 mScarlet-TUB1*), YEF11524 [*GFP-myo1(1-1797) mScarlet-TUB1)*], YEF11601 (*bni5*Δ *GFP-MYO1 mScarlet-TUB1*), and YEF11530 [*bni5Δ GFP-myo1(1-1797) mScarlet-TUB1)*].

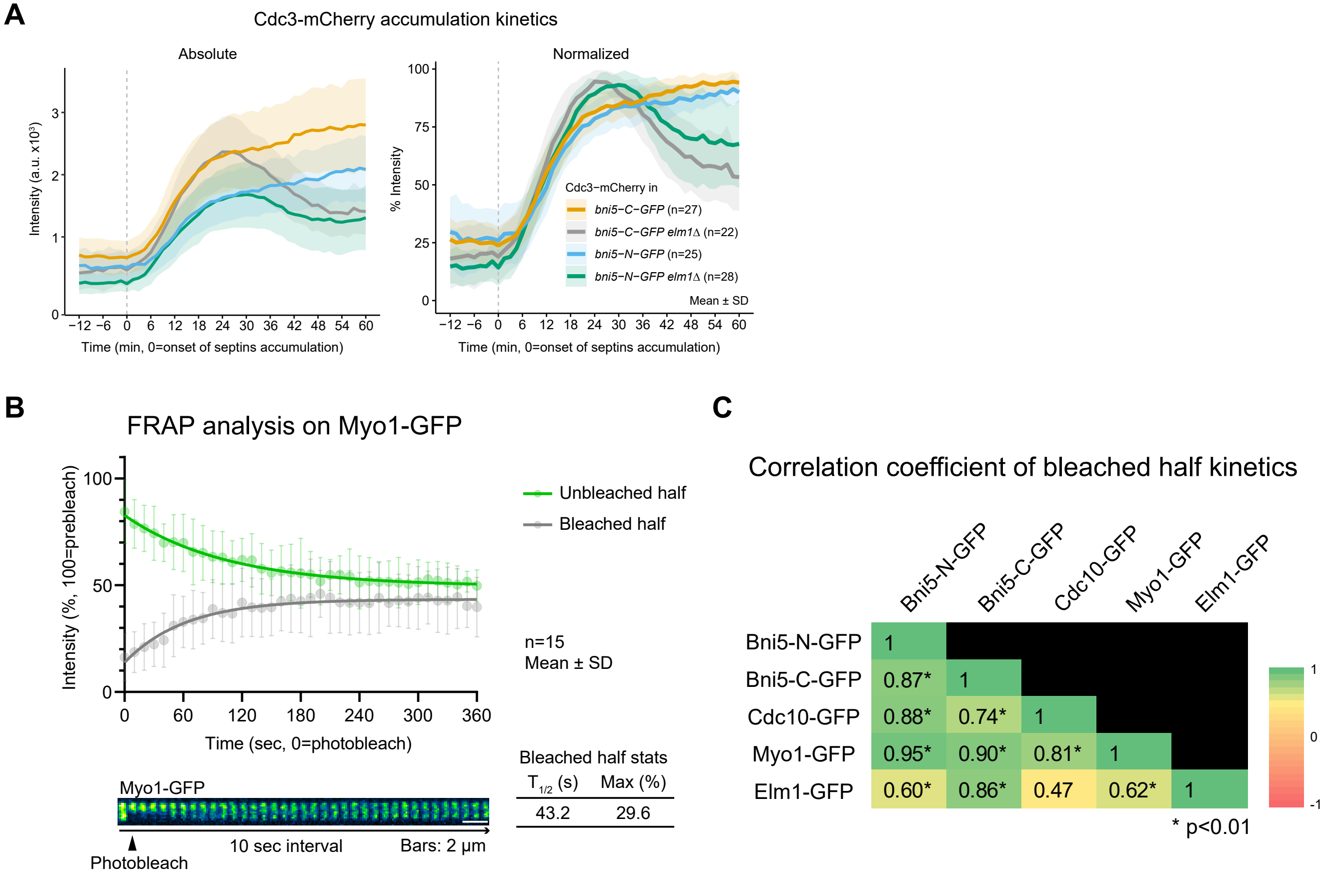


Figure S1. (Okada et al.)

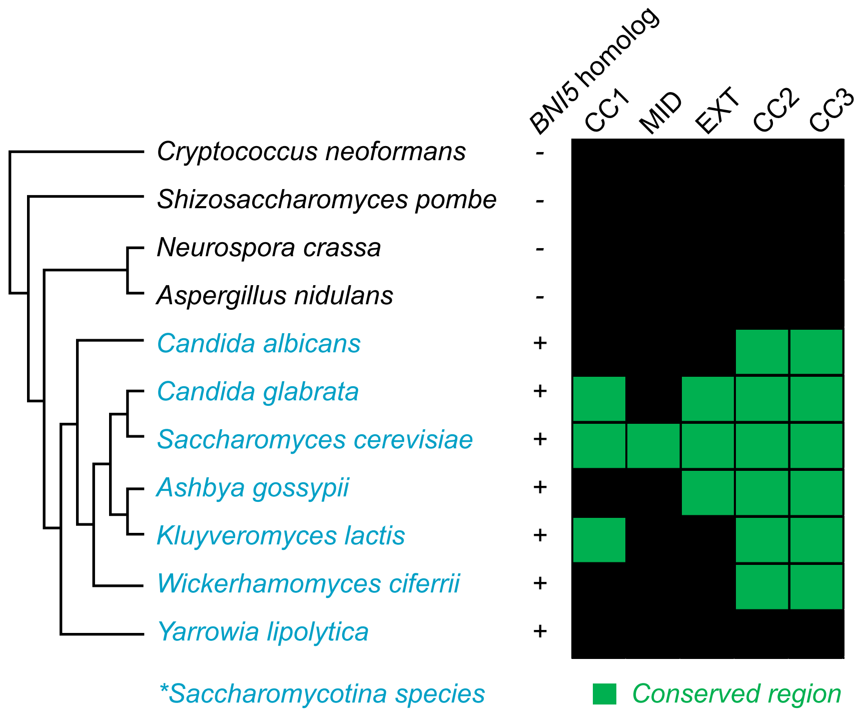


Figure S2. (Okada et al.)

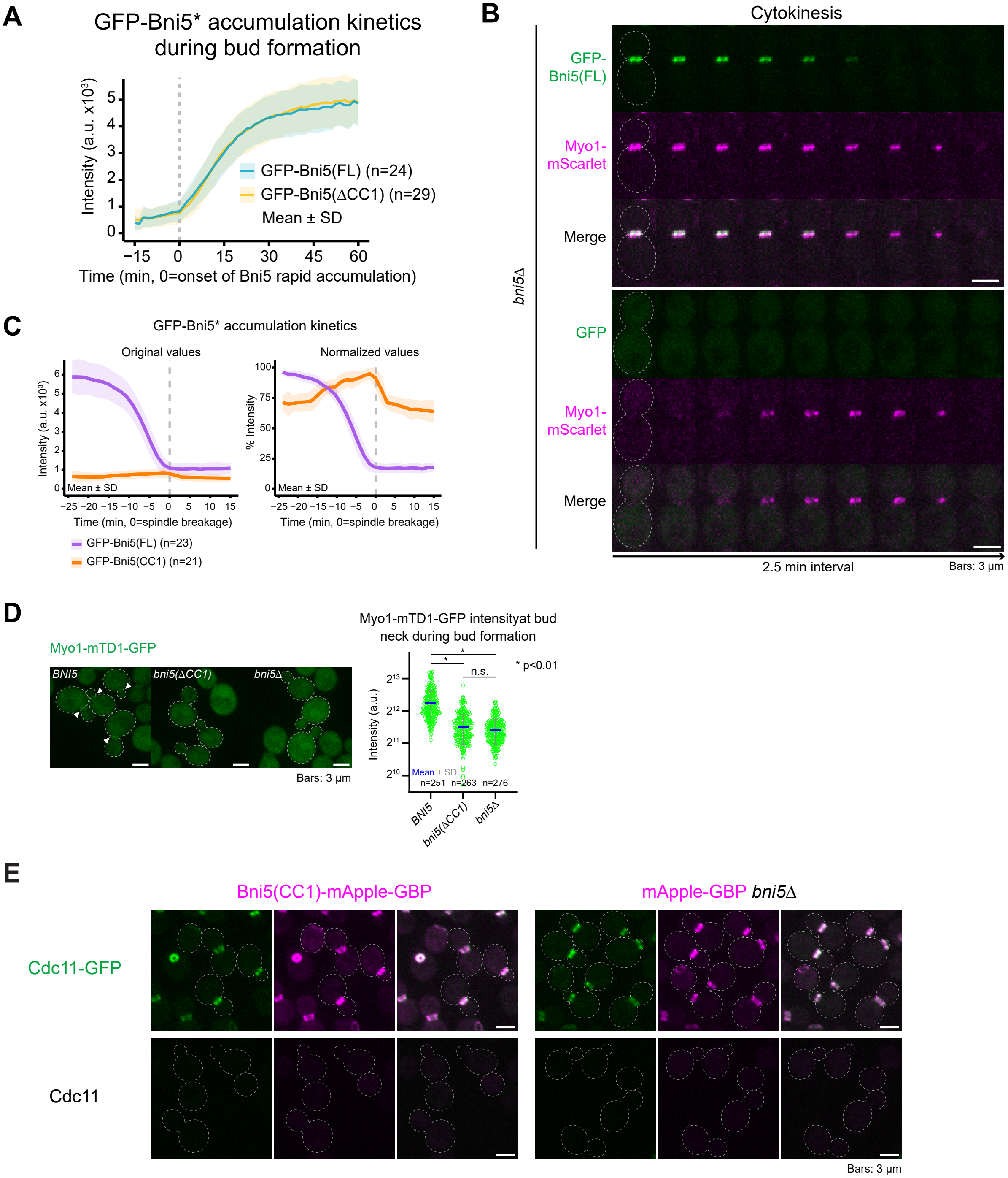


Figure S3. (Okada et al.)

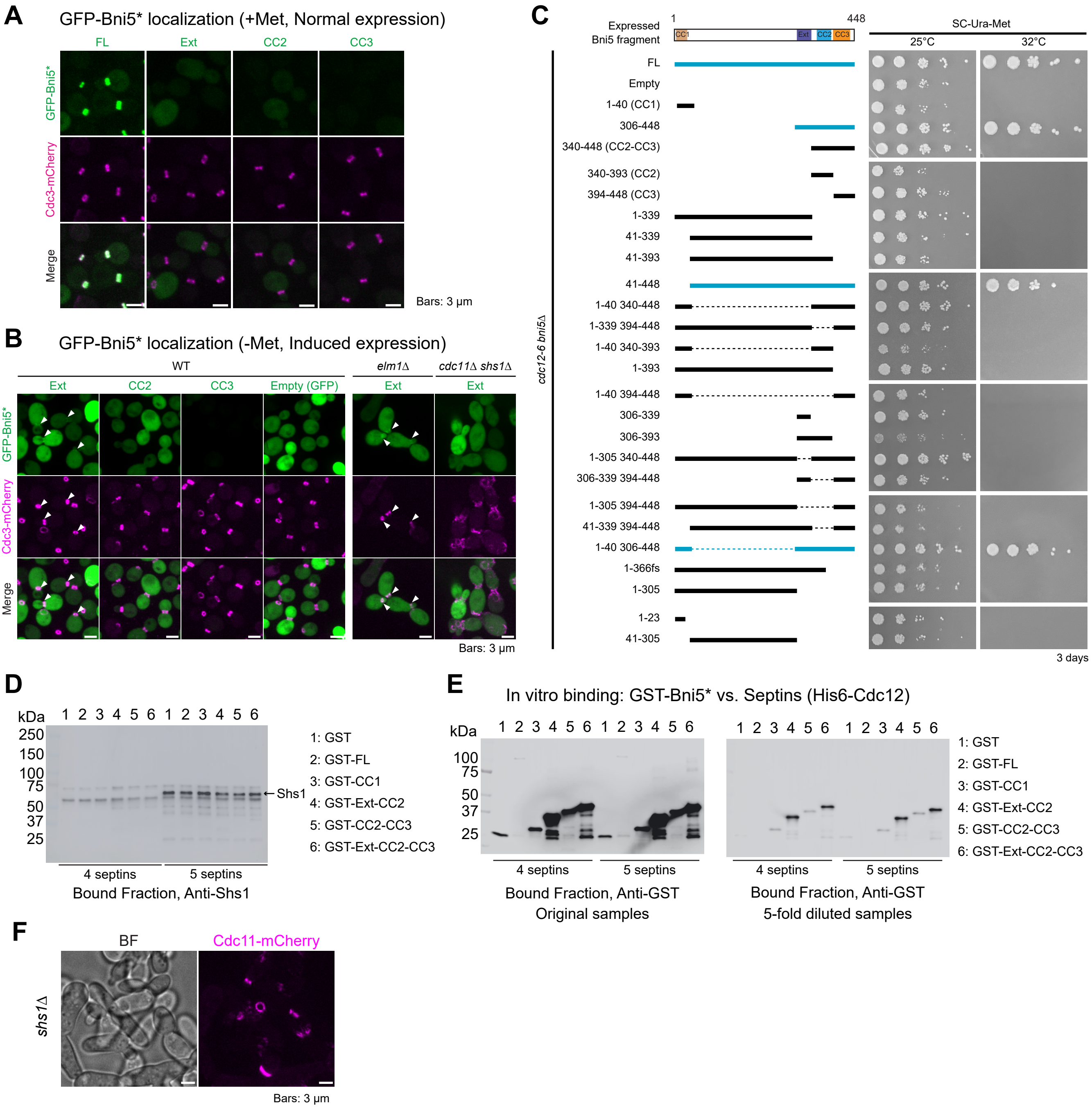


Figure S4. (Okada et al.)

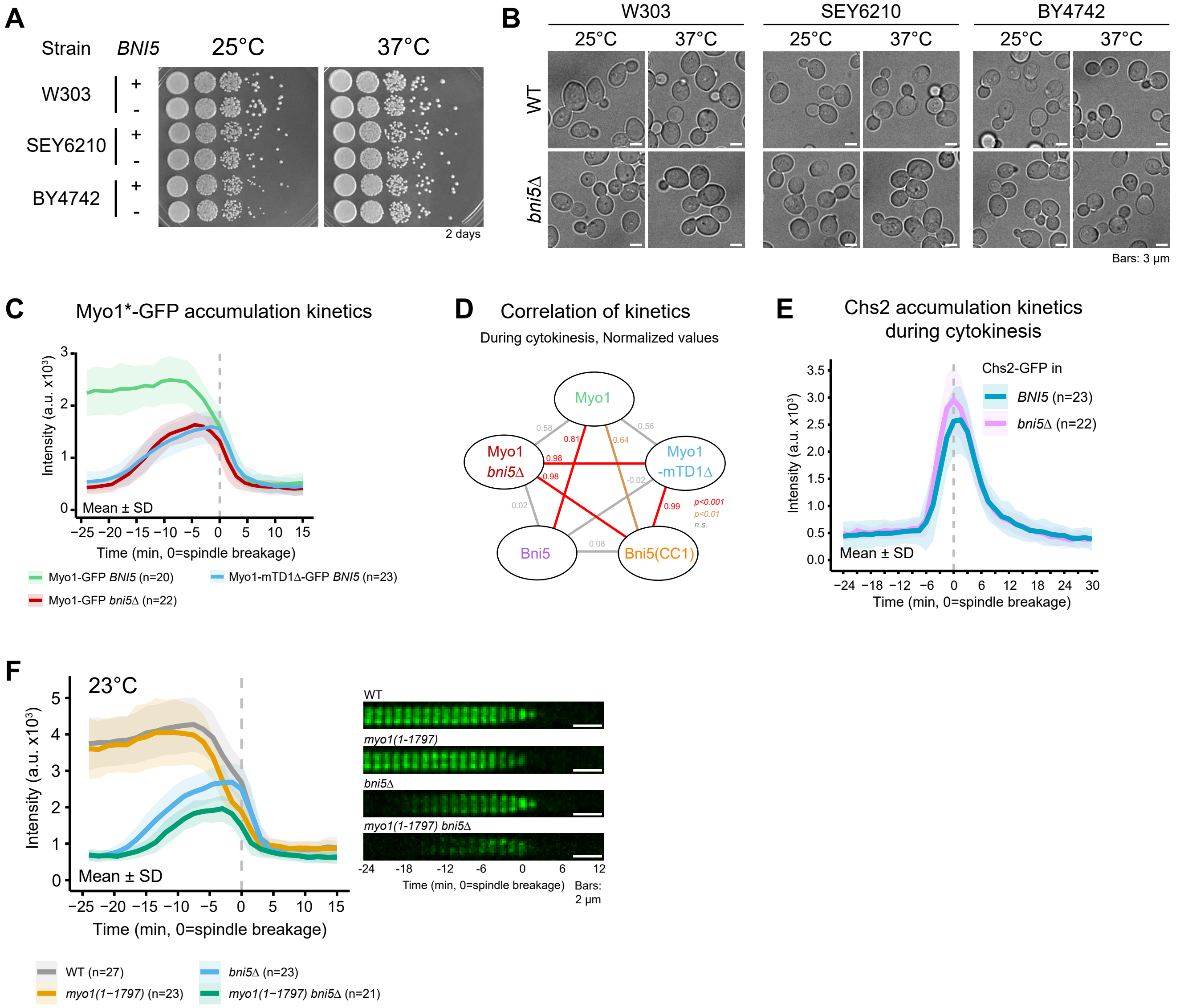


Figure S5. (Okada et al.)

| Strain | Genotype | Source |
|---|---|---|
| YEF473A | *MAT*a *trp1-Δ63 leu2-Δ1 ura3-52 his3-Δ200 lys2-801* | (Bi and Pringle, 1996) |
| W303 | *MAT*a *leu2-3,112 trp1-1 can1-100 ura3-1 ade2-1 his3-11,15* | Charles Boone |
| SEY6210 | *MAT*α *leu2-3,112 ura3-52 his3-Δ200 trp1-Δ901 suc2-Δ9 lys2-801; GAL* | Scott D Emr |
| BY4742 | *MAT*α *his3Δ1 leu2Δ0 lys2Δ0 ura3Δ0* | Christopher G Burd |
| YEF743 (M-17) | *MAT*a *cdc12-6 leu2 ura3* | (Caviston et al., 2003) |
| Masa1243 | As YEF473A except *myo1Δ::URA3-KanMX6* [CEN *HIS3 MYO1-GFP*] | (Fang et al., 2010) |
| XDY 258 | As YEF473A except *myo1-mTD1-GFP bni5Δ::His3MX6* | (Fang et al., 2010) |
| YEF4857 | As YEF473A except *CDC11-mCherry-His3MX6* | Lab stock |
| YEF5797 | As YEF473A except *cdc10Δ::KanMX6 CDC3::CDC3-mCherry-LEU2* | (Wloka et al., 2011) |
| YEF5799 | As YEF473A except *shs1Δ::KanMX6 CDC3::CDC3-mCherry-LEU2* | Lab stock |
| YEF5804 | As YEF473A except *CDC3::CDC3-mCherry-LEU2* | (Fang et al., 2010) |
| YEF6108 | As YEF473A except *MYO1-GFP-KanMX CDC3::CDC3-mCherry-LEU2* | Lab stock |
| YEF6349 | As YEF473A except *myo1Δ::His3MX6* CDC3::CDC3-mCherry-LEU2 | (Wloka et al., 2013) |
| YEF7066 | As YEF473A except *cdc11Δ::TRP1 CDC3::CDC3-mCherry-LEU2* | Lab stock |
| YEF8390 | As YEF473A except *TUB1::HPH-proHIS3-mRuby2-TUB1* | (Okada et al., 2021b) |
| YEF8391 | As YEF473A except *TUB1::HPH-proHIS3-mRuby2-TUB1 MLC2-mApple-URA3MX* | (Okada et al., 2021b) |
| YEF8437 | As YEF473A except *ELM1-GFP-His3MX6 TUB1::HPH-proHIS3-mRuby2-TUB1* | This study[a] |
| YEF8937 | As YEF473A except [CEN *URA3 MYO1*] | This study[b] |
| YEF8950 | As YEF473A except *myo1Δ::NatMX6* [CEN *URA3 MYO1*] | This study[c] |
| YEF9290 | As YEF473A except *TUB1::HPH-pHIS3-mRuby2-TUB1 BNI5-GFP-His3MX6* | This study[d] |
| YEF9336 | As YEF473A except *CDC3::CDC3-mCherry-LEU2 Bni5-GFP-His3MX6* | This study[e] |
| YEF9369 | As YEF473A except *elm1*Δ*::KanMX6 CDC3::CDC3-mCherry-LEU2 Bni5-GFP-His3MX6* | This study[f] |
| YEF9473 | As YEF473A except *GFP-MYO1* | This study[g] |
| YEF9654 | As YEF473A except *bni5Δ::NatMX6* | This study[h] |
| YEF10170 | As YEF473A except *myo1Δ::NatMX6 leu2::proACT1-GFP-ECM25-(536-588)-LEU2* [CEN *URA3 MYO1*] | This study[i] |
| YEF10173 | As YEF473A except *bni5Δ::NatMX6 leu2::proACT1-GFP-ECM25-(536-588)-LEU2* | This study[i] |
| YEF10201 | As YEF473A except *leu2::proACT1-GFP-ECM25-(536-588)-LEU2* | This study[i] |

| | | |
|---|---|---|
| YEF10243 | As YEF473A except *bni5NΔ::URA3-KanMX6* | (Marquardt et al., 2020) |
| YEF10276 | As YEF473A except *GFP-BNI5* | (Marquardt et al., 2020) |
| YEF10293 | As YEF473A except *GFP-BNI5 CDC3::CDC3-mCherry-LEU2* | This study[j] |
| YEF10296 | As YEF473A except *CDC11-GFP-His3MX6* | This study[k] |
| YEF10315 | As YEF473A except *elm1Δ::KanMX6 GFP-BNI5 CDC3::CDC3-mCherry-LEU2* | This study[g] |
| YEF10440 | As YEF473A except *ELM1-GFP-His3MX6 CDC3::CDC3-mCherry-LEU2* | (Marquardt et al., 2023) |
| YEF10667 | As YEF473A except *TUB1::HPH-proHIS3-mScarlet-I-TUB1* | This study[l] |
| YEF10688 | As YEF473A except *CHS2-GFP-TRP1* | This study[m] |
| YEF10692 | As YEF473A except *CHS2-GFP-TRP1 TUB1::HPH-proHIS3-mScarlet-I-TUB1* | This study[l] |
| YEF10885 | As YEF473A except *CDC10-mScarlet-I-URA3MX* | This study[n] |
| YEF10958 | As YEF473A *MYO1-mScarlet-I-KanMX* | This study[o] |
| YEF10993 | As YEF473A except *bni5Δ::NatMX6 MYO1-mScarlet-I-KanMX* | This study[p] |
| YEF10994 | As YEF473A except *bni5Δ::NatMX6 CDC3::CDC3-mCherry-LEU2* | This study[j] |
| YEF11020 | As YEF473A except *bni5Δ::NatMX6 MYO1-mScarlet-I-KanMX* [CEN U*RA3 GFP-bni5(306-448)*] | This study[q] |
| YEF11021 | As YEF473A except *bni5Δ::NatMX6 MYO1-mScarlet-I-KanMX* [CEN *URA3 GFP-bni5(340-448)*] | This study[q] |
| YEF11022 | As YEF473A except *bni5Δ::NatMX6 MYO1-mScarlet-I-KanMX* [CEN *URA3 GFP-bni5(340-393)*] | This study[q] |
| YEF11023 | As YEF473A except *bni5Δ::NatMX6 MYO1-mScarlet-I-KanMX* [CEN *URA3 GFP-bni5(394-448)*] | This study[q] |
| YEF11024 | As YEF473A except *bni5Δ::NatMX6 MYO1-mScarlet-I-KanMX* [CEN *URA3 GFP-bni5(1-339)*] | This study[q] |
| YEF11025 | As YEF473A except *bni5Δ::NatMX6 MYO1-mScarlet-I-KanMX* [CEN *URA3 GFP-bni5(41-339)*] | This study[q] |
| YEF11026 | As YEF473A except *bni5Δ::NatMX6 MYO1-mScarlet-I-KanMX* [CEN *URA3 GFP-bni5(41-393)*] | This study[q] |
| YEF11027 | As YEF473A except *bni5Δ::NatMX6 MYO1-mScarlet-I-KanMX* [CEN *URA3 GFP-bni5(41-448)*] | This study[q] |
| YEF11028 | As YEF473A except *bni5Δ::NatMX6 MYO1-mScarlet-I-KanMX* [CEN *URA3 GFP-bni5(Δ41-339)*] | This study[q] |
| YEF11029 | As YEF473A except *bni5Δ::NatMX6 MYO1-mScarlet-I-KanMX* [CEN *URA3 GFP-BNI5(FL)*] | This study[q] |
| YEF11030 | As YEF473A except *bni5Δ::NatMX6 MYO1-mScarlet-I-KanMX* [CEN *URA3 GFP-bni5(1-366*-frameshift)*] | This study[q] |

| | | |
|---|---|---|
| YEF11031 | As YEF473A except *bni5Δ::NatMX6 MYO1-mScarlet-I-KanMX* [CEN *URA3 GFP-bni5(1-305)*] | This study[q] |
| YEF11032 | As YEF473A except *bni5Δ::NatMX6 MYO1-mScarlet-I-KanMX* [CEN *URA3 GFP-bni5(1-23)*] | This study[q] |
| YEF11033 | As YEF473A except *bni5Δ::NatMX6 MYO1-mScarlet-I-KanMX* [CEN *URA3 GFP*] | This study[q] |
| YEF11034 | As YEF473A except *bni5Δ::NatMX6 MYO1-mScarlet-I-KanMX* [CEN *URA3 GFP-bni5(Δ340-393)*] | This study[q] |
| YEF11035 | As YEF473A except *bni5Δ::NatMX6 MYO1-mScarlet-I-KanMX* [CEN *URA3 GFP-bni5(1-40 340-393)*] | This study[q] |
| YEF11036 | As YEF473A except *bni5Δ::NatMX6 MYO1-mScarlet-I-KanMX* [CEN *URA3 GFP-bni5(1-393)*] | This study[q] |
| YEF11037 | As YEF473A except *bni5Δ::NatMX6 MYO1-mScarlet-I-KanMX* [CEN *URA3 GFP-bni5(Δ41-393)*] | This study[q] |
| YEF11038 | As YEF473A except *bni5Δ::NatMX6 CDC3::CDC3-mCherry-LEU2* [CEN *URA3 GFP-bni5(1-40)*] | This study[r] |
| YEF11039 | As YEF473A except *bni5Δ::NatMX6 CDC3::CDC3-mCherry-LEU2* [CEN *URA3 GFP-bni5(306-448)*] | This study[r] |
| YEF11040 | As YEF473A except *bni5Δ::NatMX6 CDC3::CDC3-mCherry-LEU2* [CEN *URA3 GFP-bni5(340-448)*] | This study[r] |
| YEF11041 | As YEF473A except *bni5Δ::NatMX6 CDC3::CDC3-mCherry-LEU2* [CEN *URA3 GFP-bni5(340-393)*] | This study[r] |
| YEF11042 | As YEF473A except *bni5Δ::NatMX6 CDC3::CDC3-mCherry-LEU2* [CEN *URA3 GFP-bni5(394-448)*] | This study[r] |
| YEF11043 | As YEF473A except *bni5Δ::NatMX6 CDC3::CDC3-mCherry-LEU2* [CEN *URA3 GFP-bni5(1-339)*] | This study[r] |
| YEF11044 | As YEF473A except *bni5Δ::NatMX6 CDC3::CDC3-mCherry-LEU2* [CEN *URA3 GFP-bni5(41-339)*] | This study[r] |
| YEF11045 | As YEF473A except *bni5Δ::NatMX6 CDC3::CDC3-mCherry-LEU2* [CEN *URA3 GFP-bni5(41-393)*] | This study[r] |
| YEF11046 | As YEF473A except *bni5Δ::NatMX6 CDC3::CDC3-mCherry-LEU2* [CEN *URA3 GFP-bni5(41-448)*] | This study[r] |
| YEF11047 | As YEF473A except *bni5Δ::NatMX6 CDC3::CDC3-mCherry-LEU2* [CEN *URA3 GFP-bni5(Δ41-339)*] | This study[r] |
| YEF11048 | As YEF473A except *bni5Δ::NatMX6 CDC3::CDC3-mCherry-LEU2* [CEN *URA3 GFP*] | This study[r] |
| YEF11049 | As YEF473A except *bni5Δ::NatMX6 CDC3::CDC3-mCherry-LEU2* [CEN *URA3 GFP-bni5(Δ340-393)*] | This study[r] |
| YEF11050 | As YEF473A except *bni5Δ::NatMX6 CDC3::CDC3-mCherry-LEU2* [CEN *URA3 GFP-bni5(1-40 340-393)*] | This study[r] |
| YEF11051 | As YEF473A except *bni5Δ::NatMX6 CDC3::CDC3-mCherry-LEU2* [CEN *URA3 GFP-bni5(1-393)*] | This study[r] |

| | | |
|---|---|---|
| YEF11052 | As YEF473A except *bni5Δ::NatMX6 CDC3::CDC3-mCherry-LEU2* [CEN *URA3 GFP-bni5(Δ41-393)*] | This study[r] |
| YEF11053 | As YEF473A except *bni5Δ::NatMX6 CDC3::CDC3-mCherry-LEU2* [CEN *URA3 GFP-BNI5(FL)*] | This study[r] |
| YEF11054 | As YEF473A except *bni5Δ::NatMX6 CDC3::CDC3-mCherry-LEU2* [CEN *URA3 GFP-bni5(1-366*-frameshift)*] | This study[r] |
| YEF11055 | As YEF473A except *bni5Δ::NatMX6 CDC3::CDC3-mCherry-LEU2* [CEN *URA3 GFP-bni5(1-305)*] | This study[r] |
| YEF11056 | As YEF473A except *bni5Δ::NatMX6 CDC3::CDC3-mCherry-LEU2* [CEN *URA3 GFP-bni5(1-23)*] | This study[r] |
| YEF11058 | As YEF473A except *bni5Δ::NatMX6 MYO1-mScarlet-I-KanMX* [CEN *URA3 GFP-bni5(1-40)*] | This study[q] |
| YEF11088 | As YEF473A except *CDC11-GFP-His3MX6 MYO1-mScarlet-I-KanMX* | This study[p] |
| YEF11114 | As YEF473A except *myo1Δ::His3MX6 CDC3::CDC3-mCherry-LEU2* [CEN *URA3 GFP-bni5(1-40)*] | This study[s] |
| YEF11117 | As YEF473A except *cdc10Δ::KanMX6 CDC3::CDC3-mCherry-LEU2* [CEN *URA3 GFP-bni5(306-448)*] | This study[t] |
| YEF11121 | As YEF473A except *shs1Δ::KanMX6 CDC3::CDC3-mCherry-LEU2* [CEN *URA3 GFP-bni5(306-448)*] | This study[t] |
| YEF11176 | As YEF473A except *bni5Δ::NatMX6 MYO1-mScarlet-I-KanMX* [CEN *URA3 GFP-bni5(306-339)*] | This study[q] |
| YEF11177 | As YEF473A except *bni5Δ::NatMX6 MYO1-mScarlet-I-KanMX* [CEN *URA3 GFP-bni5(306-393)*] | This study[q] |
| YEF11178 | As YEF473A except *bni5Δ::NatMX6 MYO1-mScarlet-I-KanMX* [CEN *URA3 GFP-bni5(Δ41-305)*] | This study[q] |
| YEF11179 | As YEF473A except *bni5Δ::NatMX6 MYO1-mScarlet-I-KanMX* [CEN *URA3 GFP-bni5(Δ306-339)*] | This study[q] |
| YEF11180 | As YEF473A except *bni5Δ::NatMX6 MYO1-mScarlet-I-KanMX* [CEN *URA3 GFP-bni5(306-339 394-448)*] | This study[q] |
| YEF11181 | As YEF473A except *bni5Δ::NatMX6 CDC3::CDC3-mCherry-LEU2* [CEN *URA3 GFP-bni5(306-339)*] | This study[r] |
| YEF11182 | As YEF473A except *bni5Δ::NatMX6 CDC3::CDC3-mCherry-LEU2* [CEN *URA3 GFP-bni5(306-393)*] | This study[r] |
| YEF11183 | As YEF473A except *bni5Δ::NatMX6 CDC3::CDC3-mCherry-LEU2* [CEN *URA3 GFP-bni5(Δ41-305)*] | This study[r] |
| YEF11184 | As YEF473A except *bni5Δ::NatMX6 CDC3::CDC3-mCherry-LEU2* [CEN *URA3 GFP-bni5(Δ306-339)*] | This study[r] |
| YEF11185 | As YEF473A except *bni5Δ::NatMX6 CDC3::CDC3-mCherry-LEU2* [CEN *URA3 GFP-bni5(306-339 394-448)*] | This study[r] |
| YEF11186 | As YEF473A except *bni5Δ::NatMX6 MYO1-mScarlet-I-KanMX* [CEN *URA3 GFP-bni5(Δ306-393)*] | This study[q] |

| | | |
|---|---|---|
| YEF11187 | As YEF473A except *bni5Δ::NatMX6 CDC3::CDC3-mCherry-LEU2* [CEN *URA3 GFP-bni5(Δ306-393)*] | This study[r] |
| YEF11190 | As YEF473A except *bni5Δ::NatMX6 elm1Δ::KanMX6 CDC3::CDC3-mCherry-LEU2* [CEN *URA3 GFP-BNI5(FL)*] | This study[u] |
| YEF11193 | As YEF473A *except bni5Δ::NatMX6 elm1Δ::KanMX6 CDC3::CDC3-mCherry-LEU2* [CEN *URA3 GFP-bni5(306-448)*] | This study[u] |
| YEF11194 | As YEF473A except *bni5Δ::NatMX6 elm1Δ::KanMX6 CDC3::CDC3-mCherry-LEU2* [CEN *URA3 GFP-bni5(340-448)*] | This study[u] |
| YEF11196 | As YEF473A except *bni5Δ::NatMX6 elm1Δ::KanMX6 CDC3::CDC3-mCherry-LEU2* [CEN *URA3 GFP-bni5(306-339)*] | This study[u] |
| YEF11197 | As YEF473A except *bni5Δ::NatMX6 elm1Δ::KanMX6 CDC3::CDC3-mCherry-LEU2* [CEN *URA3 GFP-bni5(306-393)*] | This study[u] |
| YEF11211 | As YEF473A except *CDC11-GFP-His3MX6 MYO1-mScarlet-I-KanMX bni5Δ::GBP-URA3MX* | This study[v] |
| YEF11212 | As YEF473A except *CDC11-GFP-His3MX6 MYO1-mScarlet-I-KanMX bni5(1-40)-GBP-URA3MX* | This study[u] |
| YEF11214 | As YEF473A except *CDC11-GFP-His3MX6 bni5Δ::mApple-GBP-URA3MX* | This study[u] |
| YEF11215 | As YEF473A except *CDC11-GFP-His3MX6 bni5(1-40)-mApple-GBP-URA3MX* | This study[u] |
| YEF11247 | *cdc12-6 bni5Δ::NatMX6* | This study[w] |
| YEF11248 | As YEF473A except *CDC11-GFP-His3MX6 MYO1-mScarlet-I-KanMX bni5Δ::NatMX6* | This study[w] |
| YEF11249 | As YEF473A except *bni5Δ::mApple-GBP-URA3MX* | This study[x] |
| YEF11250 | As YEF473A except *bni5(1-40)-mApple-GBP-URA3MX* | This study[x] |
| YEF11254 | As YEF473A except *bni5Δ::GBP-URA3MX* | This study[x] |
| YEF11255 | As YEF473A except *bni5(1-40)-GBP-URA3MX* | This study[x] |
| YEF11275 | As YEF473A except *bni5Δ::NatMX6 MYO1-mScarlet-I-KanMX* [CEN *URA3 GFP-bni5(41-339 394-448)*] | This study[q] |
| YEF11276 | As YEF473A except *bni5Δ::NatMX6 CDC3::CDC3-mCherry-LEU2* [CEN *URA3 GFP-bni5(41-339 394-448)*] | This study[r] |
| YEF11277 | As YEF473A except *bni5Δ::NatMX6 elm1Δ::KanMX6 CDC3::CDC3-mCherry-LEU2* | This study[y] |
| YEF11310 | As YEF473A except *cdc11Δ::TRP1 CDC3::CDC3-mCherry-LEU2* [CEN *URA3 GFP-bni5(306-448)*] | This study[t] |
| YEF11312 | *cdc12-6 bni5Δ::NatMX6* [CEN *URA3 GFP-BNI5(FL)*] | This study[z] |
| YEF11313 | *cdc12-6 bni5Δ::NatMX6* [CEN *URA3 GFP*] | This study[z] |

| | | |
|---|---|---|
| YEF11314 | *cdc12-6 bni5Δ::NatMX6* [CEN *URA3 GFP-bni5(1-40)*] | This study[z] |
| YEF11315 | *cdc12-6 bni5Δ::NatMX6* [CEN *URA3 GFP-bni5(306-448)*] | This study[z] |
| YEF11316 | *cdc12-6 bni5Δ::NatMX6* [CEN *URA3 GFP-bni5(340-448)*] | This study[z] |
| YEF11317 | *cdc12-6 bni5Δ::NatMX6* [CEN *URA3 GFP-bni5(340-393)*] | This study[z] |
| YEF11318 | *cdc12-6 bni5Δ::NatMX6* [CEN *URA3 GFP-bni5(394-448)*] | This study[z] |
| YEF11319 | *cdc12-6 bni5Δ::NatMX6* [CEN *URA3 GFP-bni5(1-339)*] | This study[z] |
| YEF11320 | *cdc12-6 bni5Δ::NatMX6* [CEN *URA3 GFP-bni5(41-339)*] | This study[z] |
| YEF11321 | *cdc12-6 bni5Δ::NatMX6* [CEN *URA3 GFP-bni5(41-393)*] | This study[z] |
| YEF11322 | *cdc12-6 bni5Δ::NatMX6* [CEN *URA3 GFP-bni5(41-448)*] | This study[z] |
| YEF11323 | *cdc12-6 bni5Δ::NatMX6* [CEN *URA3 GFP-bni5(Δ41-339)*] | This study[z] |
| YEF11324 | *cdc12-6 bni5Δ::NatMX6* [CEN *URA3 GFP-bni5(Δ340-393)*] | This study[z] |
| YEF11325 | *cdc12-6 bni5Δ::NatMX6* [CEN *URA3 GFP-bni5(1-40 340-393)*] | This study[z] |
| YEF11326 | *cdc12-6 bni5Δ::NatMX6* [CEN *URA3 GFP-bni5(1-393)*] | This study[z] |
| YEF11327 | *cdc12-6 bni5Δ::NatMX6* [CEN *URA3 GFP-bni5(Δ41-393)*] | This study[z] |
| YEF11328 | *cdc12-6 bni5Δ::NatMX6* [CEN *URA3 GFP-bni5(306-339)*] | This study[z] |
| YEF11329 | *cdc12-6 bni5Δ::NatMX6* [CEN *URA3 GFP-bni5(306-393)*] | This study[z] |
| YEF11330 | *cdc12-6 bni5Δ::NatMX6* [CEN *URA3 GFP-bni5(Δ306-339)*] | This study[z] |
| YEF11331 | *cdc12-6 bni5Δ::NatMX6* [CEN *URA3 GFP-bni5(306-339 394-448)*] | This study[z] |
| YEF11332 | *cdc12-6 bni5Δ::NatMX6* [CEN *URA3 GFP-bni5(Δ306-393)*] | This study[z] |
| YEF11333 | *cdc12-6 bni5Δ::NatMX6* [CEN *URA3 GFP-bni5(41-339 394-448)*] | This study[z] |
| YEF11360 | As YEF473A except *URA3-proTEF1-IQG1* | This study[aa] |
| YEF11361 | *cdc12-6 bni5Δ::NatMX6* [CEN *URA3 GFP-bni5(Δ41-305)*] | This study[z] |
| YEF11362 | *cdc12-6 bni5Δ::NatMX6* [CEN *URA3 BNI5*] | This study[z] |
| YEF11363 | *cdc12-6 bni5Δ::NatMX6* [CEN *URA3 BNI5-C-GFP*] | This study[z] |
| YEF11379 | As YEF473A except *myo1-mTD1Δ* | This study[ab] |
| YEF11380 | As YEF473A except *GFP-IQG1* | This study[ac] |
| YEF11385 | As YEF473A except *TUB1::HPH-proHIS3-mScarlet-I-TUB1 MYO1-GFP*$^{Envy}$*-URA3MX* | This study[ad] |
| YEF11390 | As YEF473A except *bni5Δ::NatMX6 CHS2-GFP-TRP1 TUB1::HPH-proHIS3-mScarlet-I-TUB1* | This study[w] |
| YEF11403 | *cdc12-6 bni5Δ::NatMX6* [CEN *URA3 GFP-bni5(1-366*-frameshift)*] | This study[z] |
| YEF11404 | *cdc12-6 bni5Δ::NatMX6* [CEN *URA3 GFP-bni5(1-305)*] | This study[z] |
| YEF11405 | *cdc12-6 bni5Δ::NatMX6* [CEN *URA3 GFP-bni5(1-23)*] | This study[z] |
| YEF11408 | As YEF473A except *myo1-mTD1Δ CDC3::CDC3-mCherry-LEU2* | This study[j] |

| | | |
|---|---|---|
| YEF11412 | As YEF473A except *GFP-IQG1 TUB1::HPH-proHIS3-mScarlet-I-TUB1* | This study[l] |
| YEF11417 | As YEF473A except *MYO1-GFP$^{Envy}$-URA3MX* | This study[ad] |
| YEF11418 | As YEF473A except *myo1-mTD1Δ-GFP$^{Envy}$-URA3MX* | This study[ad] |
| YEF11436 | As YEF473A except *CDC11-mCherry-His3MX6 shs1Δ::TRP1* | This study[ae] |
| YEF11439 | As YEF473A except *shs1Δ::KanMX6 cdc11Δ::TRP1 CDC3::CDC3-mCherry-LEU2* | This study[af] |
| YEF11441 | As YEF473A except *MYO1-GFP$^{Envy}$-URA3MX TUB1::HPH-proHIS3-mScarlet-I-TUB1* | This study[l] |
| YEF11442 | As YEF473A except *myo1-mTD1Δ-GFP$^{Envy}$-URA3MX TUB1::HPH-proHIS3-mScarlet-I-TUB1* | This study[l] |
| YEF11443 | As YEF473A except *bni5Δ::NatMX6 TUB1::HPH-proHIS3-mScarlet-I-TUB1 MYO1-GFP$^{Envy}$-URA3MX* | This study[w] |
| YEF11464 | As YEF473A except *bni5Δ::NatMX6 GFP-IQG1 TUB1::HPH-proHIS3-mScarlet-I-TUB1* | This study[w] |
| YEF11489 | As YEF473A except *myo1-mTD1Δ leu2::proACT1-GFP-ECM25-(536-588)-LEU2* | This study[i] |
| YEF11501 | As YEF473A except *GFP-myo1(1-1797)-KanMX* | This study[ag] |
| YEF11520 | As YEF473A except *bni5Δ::NatMX6 MYO1-mScarlet-I-KanMX* [CEN *URA3 GFP-bni5(41-305)*] | This study[q] |
| YEF11521 | As YEF473A except *bni5Δ::NatMX6 CDC3::CDC3-mCherry-LEU2* [CEN *URA3 GFP-bni5(41-305)*] | This study[r] |
| YEF11522 | As YEF473A except *bni5Δ::NatMX6 TUB1::HPH-proHIS3-mScarlet-I-TUB1* | This study[l] |
| YEF11524 | As YEF473A except *GFP-myo1(1-1797)-KanMX TUB1::HPH-proHIS3-mScarlet-I-TUB1* | This study[l] |
| YEF11529 | As YEF473A except GFP-MYO1 TUB1::HPH-proHIS3-mScarlet-I-TUB1 | This study[l] |
| YEF11530 | As YEF473A except *bni5Δ::NatMX6 GFP-myo1(1-1797)-KanMX TUB1::HPH-proHIS3-mScarlet-I-TUB1* | This study[w] |
| YEF11546 | As YEF473A except *bni5Δ::NatMX6 TUB1::HPH-proHIS3-mScarlet-I-TUB1 leu2::proBNI5-GFP-BNI5(FL)-LEU2MX* | This study[ah] |
| YEF11547 | As YEF473A except *bni5Δ::NatMX6 TUB1::HPH-proHIS3-mScarlet-I-TUB1 leu2::proBNI5-GFP-BNI5(1-40)-LEU2MX* | This study[ah] |
| YEF11549 | As YEF473A except *bni5Δ::NatMX6 TUB1::HPH-proHIS3-mScarlet-I-TUB1 leu2::proBNI5-GFP-bni5(306-339)-LEU2MX* | This study[ah] |
| YEF11550 | *cdc12-6 bni5Δ::NatMX6* [CEN *URA3 GFP-bni5(41-305)*] | This study[z] |
| YEF11551 | As YEF473A except *bni5Δ::NatMX6 TUB1::HPH-proHIS3-mScarlet-I-TUB1 leu2::proBNI5-GFP-bni5(340-448)-LEU2MX* | This study[ah] |
| YEF11552 | As YEF473A except *bni5Δ::NatMX6 TUB1::HPH-proHIS3-mScarlet-I-TUB1 leu2::proBNI5-GFP-bni5(306-448)-LEU2MX* | This study[ah] |

| | | |
|---|---|---|
| YEF11553 | As YEF473A except *bni5Δ::NatMX6 TUB1::HPH-proHIS3-mScarlet-I-TUB1 leu2::proBNI5-GFP-bni5(Δ41-305)-LEU2MX* | This study[ah] |
| YEF11600 | As YEF473A except *bni5Δ::NatMX6 TUB1::HPH-proHIS3-mScarlet-I-TUB1 leu2::proBNI5-GFP-bni5(41-448)-LEU2MX* | This study[ah] |
| YEF11601 | As YEF473A except *bni5Δ::NatMX6 GFP-MYO1 TUB1::HPH-proHIS3-mScarlet-I-TUB1* | This study[w] |
| YEF11624 | As YEF473A except *TUB1::HPH-proHIS3-mScarlet-I-TUB1 BNI5-GFP-His3MX6* | This study[f] |
| YEF11634 | As YEF473A except *CDC10-mScarlet-I-URA3MX bni5Δ::NatMX6* | This study[w] |
| YEF11640 | As YEF473A except *CDC10-mScarlet-I-URA3MX bni5Δ::NatMX6 TUB1::HPH-proHIS3-Venus-TUB1* | This study[ai] |
| YEF11744 | As YEF473A except *bni5Δ::NatMX6 MYO1-mScarlet-I-KanMX* [CEN *URA3 BNI5-C-GFP*] | This study[q] |
| YEF11745 | As YEF473A except *bni5Δ::NatMX6 CDC3::CDC3-mCherry-LEU2* [CEN *URA3 BNI5-C-GFP*] | This study[r] |
| YEF11750 | As YEF473A except *CDC3::CDC3-mCherry-LEU2 CDC10-GFP-His3MX6* | This study[aj] |
| YEF11753 | As YEF473A except *TUB1::HPH-proHIS3-mScarlet-I-TUB1 CDC10-GFP-His3MX6* | This study[aj] |
| YEF11757 | As YEF473A except *myo1-mTD1Δ CDC3::CDC3-mCherry-LEU2* [CEN *URA3 GFP-bni5(1-40)*] | This study[ak] |
| YEF11771 | As YEF473A except *myo1-mTD1-GFP bni5Δ::His3MX6* [CEN *URA3 BNI5(FL)*] | This study[al] |
| YEF11772 | As YEF473A except *CDC10-mScarlet-I-URA3MX bni5Δ::NatMX6 TUB1::HPH-proHIS3-Venus-TUB1 leu2::proBNI5-GFP-BNI5(FL)-LEU2MX* | This study[am] |
| YEF11773 | As YEF473A except *CDC10-mScarlet-I-URA3MX bni5Δ::NatMX6 TUB1::HPH-proHIS3-Venus-TUB1 leu2::proBNI5-GFP-bni5(Δ41-305)-LEU2MX* | This study[am] |
| YEF11774 | As YEF473A except *myo1-mTD1-GFP bni5Δ::His3MX6* [CEN *URA3 bni5(41-448)*] | This study[al] |
| YEF11775 | As YEF473A except *myo1-mTD1-GFP bni5Δ::His3MX6* [CEN *URA3*] | This study[al] |
| YEF11790 | As YEF473A except *shs1Δ::KanMX6 cdc11Δ::TRP1 CDC3::CDC3-mCherry-LEU2* [CEN *URA3 GFP-bni5(306-448)*] | This study[an] |
| YEF11821 | As YEF473A except *bni5Δ::NatMX6 leu2::proBNI5-GFP-BNI5(FL)-LEU2MX* | This study[ao] |
| YEF11823 | As YEF473A except *bni5Δ::NatMX6 leu2::proBNI5-GFP-bni5(306-393)-LEU2MX* | This study[ao] |

| | | |
|---|---|---|
| YEF11824 | As YEF473A except *bni5Δ::NatMX6 leu2::proBNI5-GFP-bni5(340-448)-LEU2MX* | This study[ao] |
| YEF11825 | As YEF473A except *bni5Δ::NatMX6 leu2::proBNI5-GFP-bni5(306-448)-LEU2MX* | This study[ao] |
| YEF11826 | As YEF473A except *bni5Δ::NatMX6 leu2::proBNI5-GFP-bni5(306-393)-LEU2MX* [CEN *HIS3 mScarlet-I-bni5(394-448)*] | This study[ap] |
| YEF11827 | As YEF473A except *bni5Δ::NatMX6 leu2::proBNI5-GFP-bni5(306-393)-LEU2MX* [CEN *HIS3 mScarlet-I*] | This study[ap] |
| YEF11828 | As YEF473A except *bni5Δ::NatMX6 leu2::proBNI5-GFP-bni5(306-448)-LEU2MX* [CEN *HIS3 mScarlet-I-bni5(394-448)*] | This study[ap] |
| YEF11829 | As YEF473A except *bni5Δ::NatMX6 leu2::proBNI5-GFP-bni5(306-448)-LEU2MX* [CEN *HIS3 mScarlet-I*] | This study[ap] |
| YEF11832 | As YEF473A except *bni5Δ::NatMX6* [CEN *HIS3 mScarlet-I-bni5(394-448)*] | This study[ap] |
| YEF11834 | As YEF473A except *bni5Δ::NatMX6* [CEN *HIS3 mScarlet-I-bni5(394-448)*] [CEN *URA3 GFP-BNI5(FL)*] | This study[aq] |
| YEF11835 | As YEF473A except *bni5Δ::NatMX6* [CEN *HIS3 mScarlet-I-bni5(394-448)*] [CEN *URA3 GFP-bni5(1-393)*] | This study[aq] |
| YEF11836 | As YEF473A except *bni5Δ::NatMX6* [CEN *HIS3 mScarlet-I-bni5(394-448)*] [CEN *URA3 GFP-bni5(1-339)*] | This study[aq] |
| YEF11837 | As YEF473A except *bni5Δ::NatMX6* [CEN *HIS3 mScarlet-I-bni5(394-448)*] [CEN *URA3 GFP-bni5(41-393)*] | This study[aq] |
| YEF11838 | As YEF473A except *bni5Δ::NatMX6* [CEN *HIS3 mScarlet-I-bni5(394-448)*] [CEN *URA3 GFP-bni5(306-339)*] | This study[aq] |
| YEF11839 | As YEF473A except *bni5Δ::NatMX6* [CEN *HIS3 mScarlet-I-bni5(394-448)*] [CEN *URA3 GFP-bni5(306-393)*] | This study[aq] |
| YEF11840 | As YEF473A except *bni5Δ::NatMX6* [CEN *HIS3 mScarlet-I-bni5(394-448)*] [CEN *URA3 GFP-bni5(340-448)*] | This study[aq] |
| YEF11841 | As YEF473A except *bni5Δ::NatMX6* [CEN *HIS3 mScarlet-I-bni5(394-448)*] [CEN *URA3 GFP-bni5(340-393)*] | This study[aq] |
| YEF11842 | As YEF473A except *bni5Δ::NatMX6* [CEN *HIS3 mScarlet-I-bni5(394-448)*] [CEN *URA3 GFP-bni5(394-448)*] | This study[aq] |
| YEF11843 | As YEF473A except *bni5Δ::NatMX6* [CEN *HIS3 mScarlet-I-bni5(394-448)*] [CEN *URA3 GFP-bni5(306-448)*] | This study[aq] |
| YEF11844 | As YEF473A except *bni5Δ::NatMX6* [CEN *HIS3 mScarlet-I-bni5(394-448)*] [CEN *URA3*] | This study[aq] |
| YEF11851 | As W303 except *bni5Δ::NatMX6* | This study[w] |
| YEF11852 | As SEY6210 except *bni5Δ::NatMX6* | This study[w] |
| YEF11853 | As BY4742 except *bni5Δ::NatMX6* | This study[w] |
| YEF11920 | As YEF473A except *bni5Δ::NatMX6* [2µ *URA3 HTT103Q-GFP*] | This study[ar] |

| | | |
|---|---|---|
| YEF11921 | As YEF473A except [2µ *URA3 HTT103Q-GFP*] | This study[ar] |
| YEF11925 | As YEF473A *shs1Δ::KanMX6 cdc11Δ::TRP1 bni5Δ::NatMX6 CDC3::CDC3-mCherry-LEU2* | This study[w] |
| YEF11926 | As YEF473A *shs1Δ::KanMX6 cdc11Δ::TRP1 bni5Δ::NatMX CDC3::CDC3-mCherry-LEU2* [CEN *URA3 GFP-bni5(306-339)*] | This study[as] |
| YEF11949 | As YEF473A except *TOM20-mScarlet-I-KanMX* [2µ *URA3 HTT103Q-GFP*] | This study[at] |

**Table S1. Strains used in this study**

[a] A DNA fragment carrying *ELM1-GFP-His3MX6* was amplified by PCR using the chromosomal DNA from YEF8250 (Marquardt et al., 2020) as the template DNA and the pair of primers P1119 and P1140, and then transformed into YEF8390 to generate YEF8437.

[b] A low-copy plasmid YCp50-MYO1 was transformed into YEF473A.

[c] A DNA fragment carrying *myo1Δ::NatMX* was amplified by PCR using the plasmid pAG25 as the template and the pair of primers P1227 and P1228 and then transformed into the YEF8937.

[d] A DNA fragment carrying *BNI5-GFP-His3MX6* was amplified by PCR using the plasmid pFA6a-link-yoEGFP-SpHIS5 as the template and the pair of primers P1166 and P1167 and then transformed into the YEF8857 (*TUB1::HPH-pHIS3-mRuby2-TUB1*, lab stock).

[e] A DNA fragment carrying *BNI5-GFP-His3MX6* was amplified by PCR using the chromosomal DNA from YEF9290 as the template and the pair of primers P1168 and Bni5 381bp DS TAG R or of primers P1526 and P1167 and then transformed into the YEF5804 or YEF10667 to generate YEF9336 or YEF11624, respectively.

[f] A DNA fragment carrying *elm1*Δ*::KanMX6* was amplified by PCR using the chromosomal DNA from YEF9246 (lab stock) or YEF8393 (Marquardt et al., 2020) as the template and the pair of primers P1139 and P1140 and then transformed into the YEF9336 or YEF10293 to generate YEF9369 or YEF10315, respectively.

[g] pRS316-N-MYO1-GFP was digested with SalI and CalI and then transformed into Masa1243. The transformation mixture was plated on SC-His plate. After 5-FOA and G418 selection, cells were grown on YPD to remove the cover plasmid (pUG23-MYO1).

[h] A DNA fragment carrying *bni5Δ::NatMX6* was amplified by PCR using the plasmid pAG25 as the template and the pair of primers P1067 and P1087 and then transformed into the YEF473A.

[i] EcoRV-digested plasmid YIp128-proACT1-GFP-ECM25-(536-588AA)-tADH1 was integrated into *leu2* locus of YEF8950, YEF9654, YEF473A, or YEF11379 to generate YEF10170, YEF10173, YEF10201, or YEF11489, respectively.

[j] BglII-digested plasmid Yip128-CDC3-mCherry was integrated into *CDC3* locus of YEF10276, YEF9654, or YEF11379 to generate YEF10293, YEF10994, or YEF11408, respectively.

[k] A DNA fragment carrying *CDC11-GFP-HiseMX6* was amplified by PCR using the gDNA of YEF4940 (lab stock) as the template and the pair of primers P1121 and P1122 and then transformed into YEF473A.

[l] XbaI-digested plasmid proHIS3-ymScarlet-I-TUB1-tTUB1-HPH was integrated into *TUB1* locus of YEF473A, YEF10688, YEF11380, YEF11417, YEF11418, YEF9654, YEF11501, or YEF9473 to generate YEF10667, YEF10692, YEF11412, YEF11441, YEF11442, YEF11522, YEF11524, or YEF11529, respectively.

[m] A DNA fragment carrying *CHS2-GFP-TRP1* was amplified by PCR using the chromosomal DNA from YEF6040 as the template and the pair of primers P121 and P410, and then transformed into the YEF473A.

[n] A DNA fragment carrying *CDC10-mScarlet-I-URA3MX* was amplified by PCR using the plasmid pFA6a-link-ymScarlet-I-CaURA as the template and the pair of primers P121 and P410, and then transformed into the YEF473A.

[o] A DNA fragment carrying *MYO1-mScarlet-I-KanMX* was amplified by PCR using the plasmid pFA6a-link-ymScarlet-I-Kan as the template and the pair of primers P226 and P517, and then transformed into the YEF473A.

[p] A DNA fragment carrying *MYO1-mScarlet-I-KanMX* was amplified by PCR using the chromosomal DNA of YEF10958 as the template and the pair of primers Myo1-249-DS-TAA and Myo1-159-UP-TAA and then transformed into the YEF9654 or YEF10296 to generate YEF10993 or YEF11088, respectively.

[q] A low-copy plasmid from pUG36-BNI5* plasmid series (see genotype column and **Table S2**) was transformed into YEF10993.

[r] A low-copy plasmid from pUG36-BNI5* plasmid series (see genotype column and **Table S2**) was transformed into YEF10994.

[s] A low-copy plasmid pUG36-BNI5(1-40) was transformed into YEF6349.

[t] A low-copy plasmid pUG36-BNI5(306-448) was transformed into YEF5797, YEF5799, or YEF7066 to generate YEF11117, YEF11121, or YEF11310, respectively.

[u] A low-copy plasmid from pUG36-BNI5* plasmid series (see genotype column and **Table S2**) was transformed into YEF11277.

[v] A DNA fragment carrying *bni5Δ::GBP-URA3MX* was amplified by PCR using the chromosomal DNA from YEF11254 as the template and the pair of primers P1527 and Bni5 381bp DS TAG R, and then transformed into the YEF11088 or YEF10296 to generate YEF11211 or YEF11214, respectively. Similarly, A DNA fragment carrying *bni5(1-40)-GBP-URA3MX* was amplified by PCR using the chromosomal DNA from YEF11255 as the template and the pair of primers P1527 and Bni5 381bp DS TAG R, and then transformed into the YEF11088 or YEF10296 to generate YEF11212 or YEF11215, respectively.

[w] A DNA fragment carrying *bni5Δ::NatMX6* was amplified by PCR using the chromosomal DNA from YEF9654 as the template and the pair of primers P1628 and P1087 and then transformed into the YEF743, YEF11088, YEF10692, YEF11385, YEF11412, YEF11524, YEF11529, YEF10885, W303, SEY6210, BY4742, or YEF11439 to generate YEF11247, YEF11248, YEF11390, YEF11443, YEF11464, YEF11530, YEF11601, YEF11634, YEF11851, YEF11852, YEF11853, or YEF11925, respectively.

[x] A DNA fragment carrying *bni5Δ::mApple-GBP-URA3MX* or *bni5(1-40)-mApple-GBP-URA3MX*, was amplified by PCR using the plasmid pFA6a-link-yomApple-GBP-CaURA as the template and the pair of primers P1546 and P1167 or P1545 and P1167, and then transformed into the YEF473A. Similarly, a DNA fragment carrying *bni5Δ::GBP-URA3MX* or *bni5(1-40)-GBP-URA3MX* was amplified by PCR using the plasmid pFA6a-link-GBP-CaURA3 as the template and the pair of primers P1546 and P1167 or P1545 and P1167, and then transformed into the YEF473A.

[y] A DNA fragment carrying *elm1Δ::KanMX6* was amplified by PCR using the chromosomal DNA from YEF7515 (lab stock) as the template and the pair of primers P1125 and P1140, and then transformed into the YEF10994.

[z] A low-copy plasmid from pUG-BNI5* plasmid series (see genotype column and **Table S2**) was transformed into YEF11247.

[aa] A DNA fragment carrying *URA3-proTEF1* was amplified by PCR using the plasmid pFA6a-URA-KanMX6 as the template and the pair of primers P1631 and P1632, and then transformed into the YEF473A. The cassette was inserted right before start codon of *IQG1*.

[ab] Two DNA fragments carrying distinct halves of *MYO1* gene; 1) chromosomal region from ~250 bp upstream of *MYO1* start codon until aa990 coding region followed by aa1181-1193 coding region, and 2) aa976-990 region followed by aa1181 coding region to ~130 bp downstream region of *MYO1* were amplified by PCR using the chromosomal DNA from YEF473A as the template DNA and the pair of primers P1518 and P1578 or P1579 and P1580, respectively. Resultant PCR products were mixed and then transformed into Masa1243. After 5-FOA and G418 selection, cells were grown on YPD to remove the cover plasmid (pUG23-MYO1).

[ac] A DNA fragment carrying *GFP* followed by SRTSGGSGGTGG linker and flanked by 40 bp of either upstream or downstream of *IQG1* start codon was amplified by PCR using the plasmid YIp128-proHIS3-yEGFP-TPM1-tADH1 as the template DNA and the pair of primers P1633 and P1634, and then transformed into YEF11360 to replace *URA3-proTEF1* cassette.

[ad] A DNA fragment carrying *MYO1-GFP-URA3MX* was amplified by PCR using the chromosomal DNA from YEF9064 (lab stock) as the template DNA and the pair of primers Myo1-159-UP-TAA and Myo1-249-DS-TAA, and then transformed into YEF10667, YEF473A, and YEF11379 to generate YEF11385, YEF11417, and YEF11418, respectively.

[ae] A DNA fragment carrying *shs1Δ::TRP1* was amplified by PCR using the chromosomal DNA from YEF7506 (lab stock) as the template DNA and the pair of primers Y178 and P1118, and then transformed into YEF4857.

[af] A DNA fragment carrying *cdc11Δ::TRP1* was amplified by PCR using the chromosomal DNA from YEF11309 as the template DNA and the pair of primers P1626 and P1122, and then transformed into YEF5799.

[ag] A DNA fragment carrying *myo1(1-1797)-KanMX* was amplified by PCR using the chromosomal DNA from YEF10459 (lab stock) as the template DNA and the pair of primers Myo1-4801F and P1580, and then transformed into YEF9473.

[ah] AscI-digested plasmid pRG205MX-proBNI5-yEGFP-BNI5(FL), pRG205MX-proBNI5-yEGFP-BNI5(1-40), pRG205MX-proBNI5-yEGFP-BNI5(306-339), pRG205MX-proBNI5-yEGFP-BNI5(340-448), pRG205MX-proBNI5-yEGFP-BNI5(306-448), pRG205MX-proBNI5-yEGFP-BNI5(Δ41-305), or pRG205MX-proBNI5-yEGFP-BNI5(41-448) was integrated into *leu2* locus of YEF11522 to generate YEF11546, YEF11547, YEF11549, YEF11551, YEF11552, YEF11553, or YEF11600, respectively.

[ai] XbaI-digested plasmid bWL722 was integrated into *TUB1* locus of YEF11634.

[aj] A DNA fragment carrying *CDC10-GFP-His3MX6* was amplified by PCR using the chromosomal DNA from YEF8831 (lab stock) as the template DNA and the pair of primers P812 and P813, and then transformed into YEF5804 and YEF10667 to generate YEF11750 and YEF11753, respectively.

[ak] A low-copy plasmid pUG36-BNI5(1-40) was transformed into YEF11408.

[al] A low-copy plasmid pUG36-BNI5(FL)-w/oGFP, pUG36-BNI5(41-448)-w/oGFP, or pUG36-w/oGFP was transformed into YEF11758 to generate YEF11757, YEF11711, or YEF11774, respectively.

[am] AscI-digested plasmid pRG205MX-proBNI5-yEGFP-BNI5(FL) or pRG205MX-proBNI5-yEGFP-BNI5(Δ41-305) was integrated into *leu2* locus of YEF11640 to generate YEF11772 or YEF11773, respectively.

[an] A low-copy plasmid pUG36-BNI5(306-448) was transformed into YEF11439.

[ao] AscI-digested pRG205MX-proBNI5-yEGFP-BNI5(FL), pRG205MX-proBNI5-yEGFP-BNI5(306-393), pRG205MX-proBNI5-yEGFP-BNI5(340-448), or pRG205MX-proBNI5-yEGFP-BNI5(306-448) was integrated into *leu2* locus of YEF9654 to generate YEF11821, YEF11823, YEF11824, or YEF11825, respectively.

[ap] A low-copy plasmid pUG34-ymScarlet-BNI5(394-448) was transformed into YEF11823, YEF11825, or YEF11832 to generate YEF11826, YEF11828, or YEF11832, respectively. A low-copy plasmid pUG34-ymScarlet-I was transformed into YEF11823 or YEF11825 to generate YEF11827 or YEF11829, respectively.

[aq] A low-copy plasmid from pUG36-BNI5* plasmid series (see genotype column and **Table S2**) was transformed into YEF11832.

[ar] A high-copy plasmid pYES2-Htt103Q-GFP was transformed into YEF9654 or YEF473A to generate YEF11920 or YEF11921, respectively.

[as] A low-copy plasmid pUG36-BNI5(306-339) was transformed into YEF11925.

[at] A DNA fragment carrying *TOM20-mScarlet-I-KanMX* was amplified by PCR using the chromosomal DNA from YEF11941 (lab stock) as the template DNA and the pair of primers P1308 and P1309, and then transformed into YEF11921.

| Plasmid | Source | Identifier |
|---|---|---|
| bWL722 (pHIS3p:Venus-Tub1+3'UTR::HPH) | (Markus et al., 2015) | |
| pAG25 | (Goldstein and McCusker, 1999) | |
| pCOLA-Duet-[His less]-Shs1 | (Garcia et al., 2011) | |
| pET His6 Sumo TEV LIC | Scott Gradia | Addgene #29659 |
| pET-His6-Sumo-$Elm1_{FL}$ | This study | BiLab collection#E2952 |
| pET-His6-Sumo-$Elm1_{1\text{-}420}$ | This study | BiLab collection#E2951 |
| pET-His6-Sumo-$Elm1_{420\text{-}640}$ | This study | BiLab collection#E2973 |
| pFA6a-link-GBP-CaURA3 | This study | BiLab collection#E2506 |
| pFA6a-link-ymScarlet-I-CaURA | (Marquardt et al., 2020) | BiLab collection#E2500 |
| pFA6a-link-yoEGFP-SpHIS5 | (Lee et al., 2013) | Addgene #44836 |
| pFA6a-link-yomApple-GBP-CaURA | This study | BiLab collection#E2498 |
| pFA6a-link-yoTagRFP-T-CaURA3 | (Lee et al., 2013) | Addgene #44877 |
| pFA6a-URA-KanMX6 | (Onishi et al., 2013) | |
| pGEX-4T-1-BNI5(1-448) | This study | BiLab collection#E2946 |
| pGEX-4T-1-BNI5(1-40) | This study | BiLab collection#E2947 |
| pGEX-4T-1-BNI5(306-393) | This study | BiLab collection#E2948 |
| pGEX-4T-1-BNI5(340-448) | This study | BiLab collection#E2949 |
| pGEX-4T-1-BNI5(306-448) | This study | BiLab collection#E2950 |
| pMAL-MYO1-mTD1 | (Fang et al., 2010) | BiLab collection#E2593 |
| pMVB128 | (Versele et al., 2004) | |
| pMVB133 | (Versele et al., 2004) | |
| pRG205MX | (Gnugge et al., 2016) | Addgene #64535 |
| pRG205MX-proBNI5-yEGFP | This study | BiLab collection#E2773 |
| pRG205MX-proBNI5-yEGFP-BNI5(Δ41-305) | This study | BiLab collection#E2793 |
| pRG205MX-proBNI5-yEGFP-BNI5(1-40) | This study | BiLab collection#E2781 |
| pRG205MX-proBNI5-yEGFP-BNI5(306-339) | This study | BiLab collection#E2784 |
| pRG205MX-proBNI5-yEGFP-BNI5(306-448) | This study | BiLab collection#E2792 |
| pRG205MX-proBNI5-yEGFP-BNI5(340-448) | This study | BiLab collection#E2791 |
| pRG205MX-proBNI5-yEGFP-BNI5(41-448) | This study | BiLab collection#E2813 |
| pRG205MX-proBNI5-yEGFP-BNI5(FL) | This study | BiLab collection#E2780 |
| proHIS3-ymScarlet-I-TUB1-tTUB1-HPH | (Ghanegolmohammadi et al., 2021) | BiLab collection#E2614 |
| pRS316-N-MYO1-GFP | (Caviston et al., 2003) | |
| pUG34 | J. H. Hegemann | |
| pUG34-ymScarlet-BNI5(394-448) | This study | BiLab collection#E2711 |
| pUG34-ymScarlet-I | This study | BiLab collection#E2699 |
| pUG36 | J. H. Hegemann | |
| pUG36-BNI5(Δ306-339) | This study[a] | BiLab collection#E2697 pUG36-BNI5* plasmid series |

| | | |
|---|---|---|
| pUG36-BNI5(Δ306-393) | This study[b] | BiLab collection#E2704 pUG36-BNI5* plasmid series |
| pUG36-BNI5(Δ340-393) | This study[c] | BiLab collection#E2677 pUG36-BNI5* plasmid series |
| pUG36-BNI5(Δ41-305) | This study[d] | BiLab collection#E2696 pUG36-BNI5* plasmid series |
| pUG36-BNI5(Δ41-339) | This study[d] | BiLab collection#E2675 pUG36-BNI5* plasmid series |
| pUG36-BNI5(Δ41-393) | This study[d] | BiLab collection#E2680 pUG36-BNI5* plasmid series |
| pUG36-BNI5(1-23) | This study[e] | BiLab collection#E2070 pUG36-BNI5* plasmid series |
| pUG36-BNI5(1-305) | This study[f] | BiLab collection#E2069 pUG36-BNI5* plasmid series |
| pUG36-BNI5(1-339) | This study[g] | BiLab collection#E2671 pUG36-BNI5* plasmid series |
| pUG36-BNI5(1-366fs) | This study[h] | BiLab collection#E2068 pUG36-BNI5* plasmid series |
| pUG36-BNI5(1-393) | This study[i] | BiLab collection#E2679 pUG36-BNI5* plasmid series |
| pUG36-BNI5(1-40 340-393) | This study[j] | BiLab collection#E2678 pUG36-BNI5* plasmid series |
| pUG36-BNI5(1-40) | This study[k] | BiLab collection#E2666 pUG36-BNI5* plasmid series |
| pUG36-BNI5(306-339 394-448) | This study[l] | BiLab collection#E2698 pUG36-BNI5* plasmid series |
| pUG36-BNI5(306-339) | This study[m] | BiLab collection#E2694 pUG36-BNI5* plasmid series |
| pUG36-BNI5(306-393) | This study[n] | BiLab collection#E2695 pUG36-BNI5* plasmid series |
| pUG36-BNI5(306-448) | This study[o] | BiLab collection#E2667 pUG36-BNI5* plasmid series |
| pUG36-BNI5(340-393) | This study[p] | BiLab collection#E2669 pUG36-BNI5* plasmid series |
| pUG36-BNI5(340-448) | This study[q] | BiLab collection#E2668 pUG36-BNI5* plasmid series |
| pUG36-BNI5(394-448) | This study[r] | BiLab collection#E2670 pUG36-BNI5* plasmid series |
| pUG36-BNI5(41-305) | This study[s] | BiLab collection#E2785 pUG36-BNI5* plasmid series |
| pUG36-BNI5(41-339 394-448) | This study[t] | BiLab collection#E2710 pUG36-BNI5* plasmid series |

| | | |
|---|---|---|
| pUG36-BNI5(41-339) | This study[u] | BiLab collection#E2672<br>pUG36-BNI5* plasmid series |
| pUG36-BNI5(41-393) | This study[v] | BiLab collection#E2673<br>pUG36-BNI5* plasmid series |
| pUG36-BNI5(41-448) | This study[w] | BiLab collection#E2674<br>pUG36-BNI5* plasmid series |
| pUG36-BNI5(41-448)-w/oGFP | This study[x] | BiLab collection#E2942<br>pUG36-BNI5* plasmid series |
| pUG36-BNI5(FL) | This study[y] | BiLab collection#E2066<br>pUG36-BNI5* plasmid series |
| pUG36-BNI5(FL)-w/oGFP | This study[z] | BiLab collection#E2709<br>pUG36-BNI5* plasmid series |
| pUG36-BNI5-C-GFP | This study[aa] | BiLab collection#E2727<br>pUG36-BNI5* plasmid series |
| pUG36-w/oGFP | This study[ab] | BiLab collection#E2943<br>pUG36-BNI5* plasmid series |
| pUG36-GFP-Ecm25-xACT(536–588aa) | (Duan et al., 2021) | |
| pYES2-Htt103Q-GFP | (Song et al., 2014) | |
| YCp50-MYO1 (CEN *URA3 MYO1*) | Susan Brown | |
| YGPM28d13 | Yeast Genomic Tiling Collection | |
| YIp128-CDC3-mCherry | (Gao et al., 2007) | BiLab collection#E1914 |
| YIp128-proACT1-GFP-ECM25-(536-588AA)-tADH1 | This study | BiLab collection#E2483 |
| YIp128-proHIS3-yEGFP-TPM1-tADH1 | (Okada et al., 2021b) | BiLab collection#E2497 |

**Table S2. Plasmids used in this study**

[a] Constructed by recombination-mediated plasmid construction (Oldenburg et al., 1997) (referred to as gap repair cloning hereafter). Two DNA fragments containing *bni5(1-305)* or *bni5(340-448)* was amplified by PCR from the plasmid pUG36-BNI5(FL) as the template and the pair of primers P1530 and P1573 or P1574 and P680, respectively. Resultant PCR products were mixed and then assembled with EcoRI linearized pUG36 in yeast cells.

[b] Constructed as [a] except primer P1575 was used instead of P1574 to amplify *bni5(394-448)* region instead of *bni5(340-448)* region.

[c] Constructed by gap repair cloning. A DNA fragment containing *bni5(394-448)* was amplified by PCR from the plasmid pUG36-BNI5(394-448) as the template and the pair of primers P1542 and P680. Resultant PCR product was then assembled with AflII and HindIII linearized pUG36-BNI5(1-339).

[d] Constructed by inverse PCR with gap repair cloning. A DNA fragment containing entire plasmid except *bni5(41-305)*, *bni5(41-339)*, or *bni5(41-393)* region was amplified by PCR from pUG36-BNI5(FL) as the template and the pair of primers P1539 and P1572, P1539 and P1540, or P1539 and P1569, respectively. Resultant PCR products were then self-assembled in yeast cells to generate pUG36-BNI5(Δ41-305), pUG36-BNI5(Δ41-339), and pUG36-BNI5(Δ41-393), respectively.

[e] This allele was identified by screening for synthetic lethality with *hof1Δ* (Nishihama et al., 2009). In this allele, there is an *amber* mutation (the codon CAG for Q24 was substituted to TAG, stop codon). A DNA fragment containing this allele was amplified from YEF6928 (lab stock) then assembled into pUG36 by gap repair cloning.

[f] This allele was allele identified by screening for synthetic lethality with *hof1Δ* (Nishihama et al., 2009). In this allele, there is an *amber* mutation at the codon for Q306. A DNA fragment containing this allele was amplified from YEF6927 (lab stock) then assembled into pUG36 by gap repair cloning.

[g] Constructed by gap repair cloning. A DNA fragment containing *bni5(1-339)* was amplified by PCR from the plasmid pUG36-BNI5(FL) as the template and the pair of primers P1530 and P1538. Resultant PCR product was then assembled with EcoRI linearized pUG36 in yeast cells.

[h] This allele was identified by screening for synthetic lethality with *hof1Δ* (Nishihama et al., 2009). In this allele, deletion of G from the codon AGA for R367 caused frameshift, encoding for six extra residues NMIIYP followed by stop codon. A DNA fragment containing this allele was amplified from YEF6926 (lab stock) then assembled into pUG36 by gap repair cloning.

[i] Constructed as [g] except *bni5(1-393)* region was amplified by the pair of primers P1530 and P1535.

[j] Constructed as [g] except *bni5(1-40 340-393)* region was amplified from plasmid pUG36-BNI5(Δ41-339) by the pair of primers P1530 and P1535.

[k] Constructed as [g] except *bni5(1-40)* region was amplified by the pair of primers P1530 and P1531.

[l] Constructed by gap repair cloning. Two DNA fragments containing *bni5(306-339)* or *bni5(394-448)* was amplified by PCR from the plasmid pUG36-BNI5(306-448) as the template and the pair of primers P1530 and P1541 or P1542 and P680, respectively. Resultant PCR products were mixed and then assembled with EcoRI linearized pUG36 in yeast cells.

[m] Constructed as [g] except *bni5(306-339)* region was amplified from plasmid pUG36-BNI5(306-448) by the pair of primers P1530 and P1538.

[n] Constructed as [g] except *bni5(306-393)* region was amplified from plasmid pUG36-BNI5(306-448) by the pair of primers P1530 and P1535.

[o] Constructed as [g] except *bni5(306-448)* region was amplified from plasmid YGPM28d13 by the pair of primers P1532 and P1533.

[p] Constructed as [g] except *bni5(340-393)* region was amplified from plasmid YGPM28d13 by the pair of primers P1534 and P1535.

[q] Constructed as [g] except *bni5(340-448)* region was amplified from plasmid YGPM28d13 by the pair of primers P1534 and P1533.

[r] Constructed as [g] except *bni5(394-448)* region was amplified from plasmid YGPM28d13 by the pair of primers P1536 and P1533.

[s] A ~0.8 kb DNA fragment containing *bni5(288-305-amber)* was acquired by EcoRI and EagI digestion of pUG36-BNI5(1-305) and ligated into EcoRI and EagI digested pUG36-BNI5(41-339).

[t] A ~0.4 kb DNA fragment containing *bni5(288-339 394-448)* was acquired by EcoRI and XhoI digestion of pUG36-BNI5(Δ340-393) and ligated into EcoRI and XhoI digested pUG36-BNI5(41-339).

[u] Constructed as [g] except *bni5(41-339)* region was amplified from plasmid YGPM28d13 by the pair of primers P1537 and P1538.

[v] Constructed as [g] except *bni5(41-393)* region was amplified from plasmid YGPM28d13 by the pair of primers P1537 and P1535.

[w] Constructed as [g] except *bni5(41-448)* region was amplified from plasmid YGPM28d13 by the pair of primers P1537 and P1533.

[x] A ~1.3 kb DNA fragment containing *BNI5(41-448)* was acquired by BamHI and XhoI digestion of pRG205MX-proBNI5-yEGFP-BNI5(41-448) and ligated into ~5.4kb vector backbone of pUG36-BNI5(FL)-w/oGFP acquired by BamHI and XhoI digestion.

[y] Constructed as WT control for research of *bni5* alleles identified by screening for synthetic lethality with *hof1Δ* (Nishihama et al., 2009). A DNA fragment containing *BNI5* was assembled into pUG36 by gap repair cloning.

[z] To remove the region encoding GFP from pUG36-BNI5(FL), a ~1.4 kb DNA fragment containing *BNI5(FL)* was acquired by SpeI and XhoI digestion of pUG36-BNI5(FL) and ligated into ~5.4kb vector backbone of the same plasmid digested with XbaI and XhoI. SpeI and XbaI generated compatible cohesive ends.

[aa] Constructed by gap repair cloning. A DNA fragment containing *BNI5-C-GFP* was amplified by PCR from the chromosomal DNA of YEF9290 as the template and the pair of primers P1166 and P1587. Resultant PCR product was then assembled with ClaI linearized pUG36-BNI5(FL)-w/oGFP in yeast cells.

[ab] The GFP region was removed from pUG36 as described in [z]. SpeI and XbaI digested pUG36 was self-assembled by ligation.

2

| Name | Sequence | Identifier |
|---|---|---|
| Bni5 381bp DS TAG R | ACAAAGTTAGCAGGGTTATCGC | N/A |
| Bni5-F tag Check | GTCTGAGCTGGGCAGTATTG | P1168 |
| Bni5-F1 | TGGTGATGCTATGTTAGTGTGAAATAGAACAACAGAA ACGCGGATCCCCGGGTTAATTAA | P1067 |
| Bni5-F5 | TGATCGTGGCGAAAATGGCCAATTTTGGATTGGAACTA AAGGTGACGGTGCTGGTTTA | P1166 |
| Bni5-R1(Nat) | TAATTTATAAATATTTATAACAACCCTTGGCGTAATGTA ATTCGAGCTCGTTTTCGACAC | P1087 |
| Bni5-R3 | TAATTTATAAATATTTATAACAACCCTTGGCGTAATGTA ATCGATGAATTCGAGCTCG | P1167 |
| CDC10+304-3' | GGCACAATCCCTAACCAAAC | P813 |
| CDC10-nt(668-687)-5' | ACAGAAGTGTTAGATCTATC | P812 |
| Cdc11-F-tag check | GTCATCGTCCACCACAACAAG | P1121 |
| Cdc11-R-check | CGATAATGACGATCCACACAAG | P1122 |
| Elm1-C-F | TACTTCCAATCCAATATTATGTCATCGTCCAGTGTG | N/A |
| Elm1-C-R | GCTCGAATTCGGATCCCTATATTTGACCATTATCTGCAA AG | N/A |
| Elm1-F1 | TTTTTTGAACGCCAGGTTAACAATAATTACTTAGCATG AACGGATCCCCGGGTTAATTAA | P1125 |
| Elm1-FL-F | TACTTCCAATCCAATATTATGTCACCTCGACAGCTTATA CCG | N/A |
| Elm1-FL-R | GCTCGAATTCGGATCCCTATATTTGACCATTATCTGCAA AG | N/A |
| Elm1-F-tag check | CCTAAAGAGAACGGGAACAGAAC | P1119 |
| Elm1-N-F | TACTTCCAATCCAATATTATGTCACCTCGACAGCTT | N/A |
| Elm1-N-R | GCTCGAATTCGGATCCTTAAATTTGACTGTGATTCCTA GAA | N/A |
| Elm1-R-check | GATTTCGCGACACAGTGG | P1140 |
| F045-INN1-Seq-2 | GATAAAAACATAGAAAGG | P45 |
| F123-CHS2-300up-from-Stop | GAATTGTGATGATTTGGATGC | P121 |
| F228-MYO1-tag-F5 | AAAAATATTGATAGTAACAATGCACAGAGTAAAATTTT CAGTGGTGACGGTGCTGGTTTA | P226 |
| F260-yeGFP-XbaI | TCTAGAATGTCTAAAGGTGAAG | P254 |
| F261-ECM25(aa536-588)-tADH1 | GAACGTTTACAAGGTTAAGGCGCGCCACTTCTAAATA AGCGAATTTCTTATG | P255 |
| Felm1-check | GAGGAACTTACTTGATCCTTCTTGAAG | P1139 |
| GBP-F | GACGGTGCTGGTTTAATGGCACAAGTTCAATTGGTTG AA | N/A |

| | | |
|---|---|---|
| GBP-R | TTTAGAAGTGGCGCGTTAATGGTGGTGATGGTGATGA GA | N/A |
| M13(-20)Forward | GTAAAACGACGGCCAGTGAA | P680 |
| Myo1-159-UP-TAA | CTAGCGAATAAAAATAGAAGCGA | N/A |
| Myo1-249-DS-TAA | GATACGGGGTGAAAGAGTT | N/A |
| Myo1-4801F | AAAGCTGAAACAAACTTAAA | N/A |
| Myo1-F1 | GAAGATCATAACAAAGTTAGACAGGACAACAACAGC AATACGGATCCCCGGGTTAATTAA | P1227 |
| Myo1-R1 | AAAGGATATAAAGTCTTCCAAATTTTTAAAAAAAAGTT CGGAATTCGAGCTCGTTTAAAC | P1228 |
| P1518-F-Myo1-250up-Start | AACGCTGAGATAGCTTTTCTTAC | P1518 |
| P1526-F-Bni5-(220) | AGAACAAGTTAGACGCTGAACTTG | P1526 |
| P1527-F-Bni5-250up-ATG | TGGCAAATGTATGTAGCACTTCTC | P1527 |
| P1530-F-GFP(aa150)-in-pUG36 | TCACAATGTTTACATCATGGCTGAC | P1530 |
| P1531-R-pUG36-link-Bni5(40)-Stop | ATGACTCGAGGTCGACGGTATCGATAAGCTTGATATCT TAAGCTTCGTCCTCCGCAGGTT | P1531 |
| P1532-F-pUG36-link-ATG-Bni5(306) | GTACAAATCTAGAACTAGTGGATCCCCCGGGCTGCAG ATGCAGAAAAAAATGGCGAATTT | P1532 |
| P1533-R-pUG36-link-Bni5(448)-Stop | ATGACTCGAGGTCGACGGTATCGATAAGCTTGATATCT TATTTAGTTCCAATCCAAAATT | P1533 |
| P1534-F-pUG36-link-ATG-Bni5(340) | GTACAAATCTAGAACTAGTGGATCCCCCGGGCTGCAG ATGTCACTAAATAAGTATGATTC | P1534 |
| P1535-R-pUG36-link-Bni5(393)-Stop | ATGACTCGAGGTCGACGGTATCGATAAGCTTGATATCT TAAGAACCCTGTGCATTCATCT | P1535 |
| P1536-F-pUG36-link-ATG-Bni5(394) | GTACAAATCTAGAACTAGTGGATCCCCCGGGCTGCAG ATGTTATCTATGGAAGACGGAAA | P1536 |
| P1537-F-pUG36-link-ATG-Bni5(41) | GTACAAATCTAGAACTAGTGGATCCCCCGGGCTGCAG ATGGTCGAAGATAATGTCAAGGA | P1537 |
| P1538-R-pUG36-link-Bni5(339)-Stop | ATGACTCGAGGTCGACGGTATCGATAAGCTTGATATCT TAAGATTTACGACTTCCGTTTC | P1538 |
| P1539-R-Bni5(40) | AGCTTCGTCCTCCGCAGGTTCAACTAATTGTAAATTTT CG | P1539 |
| P1540-F-Bni5(40)-Bni5(340) | AACCTGCGGAGGACGAAGCTTCACTAAATAAGTATGA TTCGCCAGTCTCCTCTCCTATCA | P1540 |
| P1541-R-Bni5(339) | AGATTTACGACTTCCGTTTCTTGAATTGGAATTGCTGC TC | P1541 |
| P1542-F-Bni5(339)-Bni5(394) | GAAACGGAAGTCGTAAATCTTTATCTATGGAAGACGG AAAGAGACTACATAGAGCCGTAG | P1542 |
| P1545-F-Bni5(40)-F5 | CGAAAATTTACAATTAGTTGAACCTGCGGAGGACGAA GCTGGTGACGGTGCTGGTTTA | P1545 |

| | | |
|---|---|---|
| P1546-F-Bni5(1)-F5 | TGATGCTATGTTAGTGTGAAATAGAACAACAGAAACG ATGGGTGACGGTGCTGGTTTA | P1546 |
| P1569-F-Bni5(40)-Bni5(394) | AACCTGCGGAGGACGAAGCTTTATCTATGGAAGACGG AAAGAGACTACATAGAGCCGTAG | P1569 |
| P1572-F-Bni5(40)-Bni5(306) | AACCTGCGGAGGACGAAGCTCAGAAAAAAATGGCGA ATTTCGAGACACGACGCCCTACAA | P1572 |
| P1573-R-Bni5(305) | TCCATCACCATCTGAATTAAAGTGATTTAT | P1573 |
| P1574-F-Bni5(305)-Bni5(340) | TTAATTCAGATGGTGATGGATCACTAAATAAGTATGATT CGCCAGTCTCCTCTCCTATCA | P1574 |
| P1575-F-Bni5(305)-Bni5(394) | TTAATTCAGATGGTGATGGATTATCTATGGAAGACGGA AAGAGACTACATAGAGCCGTAG | P1575 |
| P1578-R-MYO1-delta-mTD1 | TTCCAATTCTTTAATTGATATTTGCTTTATTAGTTCGTTA TTTTTTGAGTGTAATTTCTC | P1578 |
| P1579-F-MYO1-delta-mTD1 | AACTAATAAAGCAAATATCAATTAAAGAATTGGAAGCT CGGTTGTCACAGGAAATATCC | P1579 |
| P1580-R-MYO1-130down-stop | ACTTAGTATATAACGCTCGTGTCGTC | P1580 |
| P1581-F-BNI5pro-SacI-pRG305MX | TATATTTCTTTTCGCGAGCTCGCATTGGGAGTCTATCAT AACGTATTTATATATCCTCAT | P1581 |
| P1582-R-EGFP-link-pRG305MX | CAGCCCGGGGGATCCACTAGTTCTAGACTTGTATAATT CATCCATGCCCAACGTTATACC | P1582 |
| P1583-F-pUG3x-ymScarlet | TTACCCCCATCCATACTCTAGAATGGTGTCTAAAGGTG AAGCCGTTATCA | P1583 |
| P1584-R-pUG3x-ymScarlet | AGCCCGGGGGATCCACTAGTTCTAGACTTGTACAACT CATCCATACCACCAGTAGAATG | P1584 |
| P1626-F-Cdc11-520up-from-ATG | AATCACTAGTATTCTTATTCCATG | P1626 |
| P1628-F-Bni5-802up-from-ATG | TCGCATTGGGAGTCTATCATAACG | P1628 |
| P1631-F-Iqg1-ura-TEFpro-F1 | GCACCAGTTCAATTATATGTAACAAGGTGGTGCAAAA ACACGGATCCCCGGGTTAATTAA | P1631 |
| P1632-R-Iqg1-ura-TEFpro | TATTGCCTGGTTTCGAAGGAGAGCCTGAATATGCTGTC ATGGTTGTTTATGTTCGGATGT | P1632 |
| P1633-F-GFP-to-Iqg1-ura-TEFpro | GCACCAGTTCAATTATATGTAACAAGGTGGTGCAAAA ACAATGTCTAAAGGTGAAGAATT | P1633 |
| P1634-R-GFP-to-Iqg1-ura-TEFpro | TATTGCCTGGTTTCGAAGGAGAGCCTGAATATGCTGTC ATACCACCTGTTCCTCCGCTAC | P1634 |
| R093-CHS2-300down-from-Stop | TCAAAAGCTCTTGATGCCCA | P410 |
| R108-GBP-with-AscI-term | GTCATGGCGCGCCTTAATGGTGGTGATGGTG | P425 |
| R200-MYO1-tag-R3 | TAATGCATATTCTCATTCTGTATATACAAAACATCTCATC ATTCGATGAATTCGAGCTCG | P517 |
| R238-ECM25(aa536-588)-TAA | TTAACCTTGTAAACGTTCTTCGTAC | P549 |
| R239-GFP-proACT1 | CCTTTAGACATTCTAGATGTTAATTCAGTAAATTTTCGA TCTTGGGAAG | P550 |

| | | |
|---|---|---|
| Relm1-check | GATTTCGCGACACAGTGG | P1140 |
| Shs1-R-check | GCTTTACTTTCTGACCTTCG | P1118 |
| Shs1-AMP-520 | ACCACCTTTTTCCATACGA | Y178 |
| Tom20-checking F | CTATGTCATAACTCTCGTCCAGAATG | P1309 |
| Tom20-R3 | GAAACAAAAACGGAGAAAAAAAGCAAGCAAAATGTTACTCTCGATGAATTCGAGCTCG | P1308 |

**Table S3. Oligonucleotides used in this study**
